# Multilayer Network Analysis across Cortical Depths in Resting-State 7T fMRI

**DOI:** 10.1101/2023.12.23.573208

**Authors:** Parker Kotlarz, Kaisu Lankinen, Maria Hakonen, Tori Turpin, Jonathan R. Polimeni, Jyrki Ahveninen

## Abstract

In graph theory, “multilayer networks” represent systems involving several interconnected topological levels. One example in neuroscience is the stratification of connections between different cortical depths or “laminae”, which is becoming non-invasively accessible in humans using ultra-high-resolution functional MRI (fMRI). Here, we applied multilayer graph theory to examine functional connectivity across different cortical depths in humans, using 7T fMRI (1-mm^3^ voxels; 30 participants). Blood oxygenation level dependent (BOLD) signals were derived from five depths between the white matter and pial surface. We compared networks where the inter-regional connections were limited to a single cortical depth only (“layer-by-layer matrices”) to those considering all possible connections between areas and cortical depths (“multilayer matrix”). We utilized global and local graph theory features that quantitatively characterize network attributes including network composition, nodal centrality, path-based measures, and hub segregation. Detecting functional differences between cortical depths was improved using multilayer connectomics compared to the layer-by-layer versions. Superficial depths of the cortex dominated information transfer and deeper depths drove clustering. These differences were largest in frontotemporal and limbic regions. fMRI functional connectivity across different cortical depths may contain neurophysiologically relevant information; thus, multilayer connectomics could provide a methodological framework for studies on how information flows across this stratification.

## Introduction

Investigating brain activity and function through network analyses has become an integral methodological foundation of neuroscience. Connectomics has yielded significant advances in understanding brain structure and function (Farahani et al., 2019; Milano et al., 2019). Modeling the brain as a system of nodes (brain regions) connected by edges (mathematical relationships)— often using graph theory—can be used to provide insight into brain characteristics and topological properties (Rubinov & Sporns, 2010). Brain networks can be derived from structural neuroimaging such as MRI or diffusion tensor imaging (DTI) (structural connectomics) (Griffa et al., 2013; Meoded et al., 2020; Yeh et al., 2021) or functional neuroimaging such as fMRI, EEG, or magnetoencephalography (MEG) (functional connectomics) (De Vico Fallani et al., 2014; Matthews & Hampshire, 2016; Sadaghiani et al., 2022; Smith et al., 2013; Xia & He, 2017). Both structural and functional connectomics have been used to understand disease models (Benito-Leon et al., 2019; Fleischer et al., 2019; Jacob et al., 2020; Kotlarz et al., 2022), aid in surgical mapping (Ahsan et al., 2020; Dadario et al., 2021; Gleichgerrcht et al., 2020; Hart et al., 2016), and characterize therapeutic effects of neuropsychiatric treatments (Caeyenberghs et al., 2017; Chen et al., 2020; Filippi et al., 2023; Lei et al., 2021; Tavakol et al., 2019; Yun & Kim, 2021).

An emerging field within connectomics, and more broadly graph theory, is the exploration of multilayer networks (Boccaletti et al., 2014; Kivela et al., 2014). Multilayer networks are composed of individual layers of networks with interconnecting edges between different layers. Connections across layers can be solely between homologous nodes (multiplex) or between nodes regardless of layer or nodal position (multilayer). Multilayer connectomics enables the study of multifaceted and multimodal neuroimaging data, with the different groups of data divided into distinct layers of the connectivity matrix (Betzel & Bassett, 2017; De Domenico, 2017; Vaiana & Muldoon, 2018). For example, multilayer networks can be derived using correlations between different frequency bands of MEG recordings to identify the interplay between frequencies (Buldu & Porter, 2018). Additionally, different modalities such as MEG, fMRI, and diffusion MRI can be combined to identify patterns in brain processing (Breedt et al., 2023) or pathological dysfunction (Casas-Roma et al., 2022) that were not found in traditional single-layer analysis. Thus, multilayer connectomics allows for the incorporation of multidimensional neuroimaging data and can identify relationships between distinct neuroimaging techniques and analyses.

One potential application of multilayer connectomics is understanding the hierarchical organization of the cerebral cortex. Neuroanatomical (Felleman & Van Essen, 1991; Rockland & Pandya, 1979; Zeki, 2018) and electrophysiological (Schroeder & Foxe, 2002; Schroeder et al., 2001) studies in animal models have identified that laminar input/output patterns can inform about bottom-up (feedforward) or top-down (feedback) processes between cortical regions. Despite its high resolution, a limitation of laminar electrophysiological recordings in comprehensive connectivity analyses is that the coverage area is typically very small. At the same time, mapping anatomical connections using fiber tracing has limited options for mapping of the post-synaptic targets (Rockland, 2019). Intracortical analyses of high resolution fMRI data have the benefit that the coverage can be extended to the entire cerebral cortex. Advancements in high-resolution fMRI (≤ 1 mm^3^ voxel size) have enabled sampling of functional signals from different depths of the cortical gray matter (Finn et al., 2019; L. Huber et al., 2021; Norris & Polimeni, 2019; Polimeni et al., 2018). However, there are multiple challenges and unanswered questions for the feasibility of using cortical depth profiles of fMRI signals (Norris & Polimeni, 2019). Because deoxygenated blood also drains up to the cortical surface through the intracortical diving venules, fMRI voxels intersecting the superficial layers could also be affected by deeper neuronal activations (Markuerkiaga et al., 2016b). Despite this limitation, studies using this emerging methodology have attempted to identify feedback and feedforward relationships non-invasively in the human brain (Chai et al., 2021; De Martino et al., 2015; Fracasso et al., 2018; Gau et al., 2020; Klein et al., 2018; Kok et al., 2016; Lankinen et al., 2023; Lawrence et al., 2019; Moerel et al., 2018; Moerel et al., 2019; Muckli et al., 2015; Wu et al., 2018), akin to micro-scale recordings in animal models.

Even with the advent of high-resolution functional neuroimaging, multilayer connectomics have mostly focused on anatomical networks derived from structural MRI and DTI (Shamir & Assaf, 2021; Shamir & Assaf, 2023) due to their direct relationship to cortical architecture. For example, DTI and histological samples identified that cortical areas with similar laminar structure were more likely to be connected (Wei et al., 2019). Additionally, even in functional laminar studies, previous works have predominantly utilized task-based studies in pre-defined brain regions (Chai et al., 2021; De Martino et al., 2015; Finn et al., 2019; Fracasso et al., 2018; Gau et al., 2020; Klein et al., 2018; Kok et al., 2016; Lankinen et al., 2023; Lawrence et al., 2019; Moerel et al., 2018; Moerel et al., 2019; Muckli et al., 2015; Polimeni et al., 2010; Wu et al., 2018). Consequently, the functional components of the whole-brain cortical depths continue to be underexplored.

In contrast to task-based studies, which primarily focus on specific cortical areas (Finn et al., 2021), resting-state analysis enables whole-brain investigation of laminar organization (L. Huber et al., 2021; L. R. Huber et al., 2021). These resting-state connections have been shown to reflect anatomical connectivity (Adachi et al., 2012; Honey et al., 2009; Turk et al., 2016; van den Heuvel et al., 2016) and task-based networks (Di et al., 2013; Hermundstad et al., 2013). Thus, network differences within laminar resting-state fMRI networks represent functional differences between cortical depths. This work explores the laminar structure of the cortex using high-resolution resting-state fMRI and multilayer connectomics. We use a dual-pipeline approach in comparing the information extracted from layer-by-layer vs. multilayer connectomics to test whether there are connectivity differences between cortical depths. We demonstrate the validity of multilayer functional laminar connectomics through showing that cortical depths have distinct graph theory characteristics that are more clearly identifiable through multilayer connectomics compared to the traditional single layer methodology.

## Methods

### Participants

Thirty healthy adults (mean age ± standard deviation = 32.4 ± 10 years, 15 women, all right-handed) were recruited using an internal online recruiting platform. Participants were screened for vision problems, hearing problems, cognition-altering medications, and exclusions for MRI (metal in the body). Twenty-eight of the participants were native English-speakers. Informed consent was obtained from all participants, and MRI safety screening forms were completed before each scan. The study design, protocol, and consent were approved by the Mass General Brigham Institutional Review Board.

### Image Acquisition

Participants were measured in sets of 7.9-min resting-state fMRI scans occurring on different days (3 to 4 sessions per participant). Twenty-three participants were measured in twelve resting-state scans. Seven participants had between ten to eighteen resting-state scans (10 scans: *n* = 1; 11 scans: *n* = 1; 13 scans: *n* = 2; 14 scans: *n* = 2, 18 scans: *n* = 1) (**Figure S1**). The participants were instructed to avoid movement during the scans and keep their eyes open and fixated on a fixation cross projected on a screen viewed through a mirror. The average duration of the sessions was around two hours. Breathing and heart rate were recorded using the built-in Siemens system at a sampling rate of 400 Hz. Inhalation and exhalation were measured with the Siemens respiratory-effort transducer attached to a respiratory belt. The heart rate was recorded using Siemens photoplethysmogram transducers on the participant’s index finger.

The functional and structural neuroimaging data was acquired using a 7T whole-body MRI scanner (MAGNETOM Terra, Siemens, Erlangen, Germany) with a home-built custom-built 64-channel array coil (Mareyam et al., 2020). To reduce participant head motion inside the scanner, MRI-compatible paddings were placed around the head and neck. In each imaging session, T_1_- weighted anatomical images were measured using a 0.75-mm isotropic multi-echo MPRAGE pulse sequence (van der Kouwe et al., 2008; Zaretskaya et al., 2018) with repetition time (TR) = 2530 ms, four echoes with echo time (TE) of 1.72, 3.53, 5.34, and 7.15 ms, 7° flip angle, 240 × 240 mm^2^ field of view (FoV), and 224 sagittal slices. To help with pial surface placement by avoiding dura mater, T_2_-weighted anatomical images (voxel size = 0.83 x 0.83 x 0.80 mm, TR = 9000 ms, TE = 269 ms, flip angle = 120°, FoV = 225 x 225 mm^2^, 270 sagittal slices) were acquired for twenty-eight out of thirty participants in one of the imaging sessions. Resting-state functional imaging was collected using a T_2_*-weighted blipped-CAIPI (Setsompop et al., 2012) simultaneous multi-slice (SMS) echo planar imaging (EPI) sequence using multi-band RF pulses (Setsompop et al., 2012) with 4× acceleration factor in phase-encoding direction, 3× acceleration factor in slice-encoding direction, TR = 2800 ms, TE = 27.0 ms, isotropic 1-mm^3^ voxels, 78° flip angle, 192 × 192 mm^2^ FoV, 132 axial slices, anterior-to-posterior phase encoding direction, 1446 Hz/pixel bandwidth, 0.82 ms nominal echo spacing, and fat suppression. In addition, to de-warp the functional data, an EPI scan was collected with identical parameters but with an opposite phase-encoding polarity (posterior-to-anterior, PA-EPI) relative to the functional scans. For four participants with missing PA-EPI scans, the data were de-warped using a gradient-echo field map (TR = 1040 ms, TE = 4.71 ms and 5.73 ms, isotropic 1.3-mm^3^ voxels; flip angle = 75°, FoV = 240 × 240 mm^2^, 120 slices, bandwidth = 303 Hz/pixel).

### MRI Preprocessing

First, SPM12 (http://www.fil.ion.ucl.ac.uk/spm/, [SPM12-spm_preproc_run.m]; bias field correction, full-width at half-maximum, FWHM: 18 mm, sampling distance: 2 mm, bias regularization: 1E−4) and customized MATLAB scripts were used to correct the bias field of the structural T_1_ and T_2_ images. Next, recon-all of FreeSurfer 6.0 (Fischl, 2012) with an extension for submillimeter 7 T data (Zaretskaya et al., 2018) was used to automatically create cortical reconstructions for each participant. An average of multiple T_1_-weighted anatomical volumes (3 to 4 per participant) alongside a T_2_-weighted volume were used in the reconstruction to enhance the quality of the cortical surfaces. Nine intermediate surfaces were created between the white matter and pial surfaces with fixed relative distances, of which five were selected for the laminar analysis (described below). Lastly, the surfaces generated by recon-all were corrected manually for inaccuracies with Recon Edit of Freeview.

For the functional data, slice-timing and motion corrections were first implemented in FreeSurfer 7.1 (Fischl, 2012). De-warping was then used to correct for geometric distortions caused by susceptibility-induced off-resonance fields. In de-warping, the off-resonance distortion field was estimated using the functional data and the PA-EPI scan collected with reversed a phase-encode blip; thus, the distortions are reversed in direction in respect to the scans [FreeSurfer: topup, applytopup] (Andersson et al., 2003; Smith et al., 2004). For four participants that were missing the PA-EPI scan used above, the distortion field was estimated using the B_0_ field map scan in FreeSurfer 6.0 [FreeSurfer-epidewarp]. The respiratory and heart rate artifacts were corrected using the RETROspective Image CORrection (RETROICOR) algorithm (3^rd^ order heart rate, respiratory, and multiplicative terms) (Glover et al., 2000). Three participants were missing heart rate data and, therefore, only respiratory recordings were used in RETROICOR. In addition, RETROICOR was not applied to five participants with missing respiratory and heart rate data. Functional data were then co-registered with the structural images using Boundary-Based Registration in FreeSurfer 6.0 (Greve & Fischl, 2009). By projecting each intersecting voxel onto the corresponding surface vertex using trilinear interpolation, the fMRI timeseries were then resampled onto the pial and white matter surfaces, and nine cortical depths between them.

From the nine intracortical surfaces, five alternating depths were selected starting closest to pial surface (depths 1 to 5, superficial to deep) (**Figure 1A**). The outside surfaces (pial and white matter) were excluded to avoid partial volume effects from the cerebrospinal fluid and white matter, respectively. Additionally, depths included were alternated to minimize potential for partial volume overlap between surfaces that would bias the correlation matrix generation. To measure this overlap, the distance from each voxel centroid (within cortical volume) to the cortical surface (white matter/pial surface) was calculated. The relative distance was defined so that the depth at pial surface was zero, and one at the white matter border. Next, voxels intersecting each layer were picked and plotted with respect to their relative distances in a histogram. **Figure S2** illustrates that taking every other layer limits the overlap between the layers, and thus leaking of information to adjacent layers. Additionally, to explore if tSNR impacted connectivity matrix generation, the average tSNR per each cortical depth was calculated for each layer (**Figure S3**).

**Figure 1.**
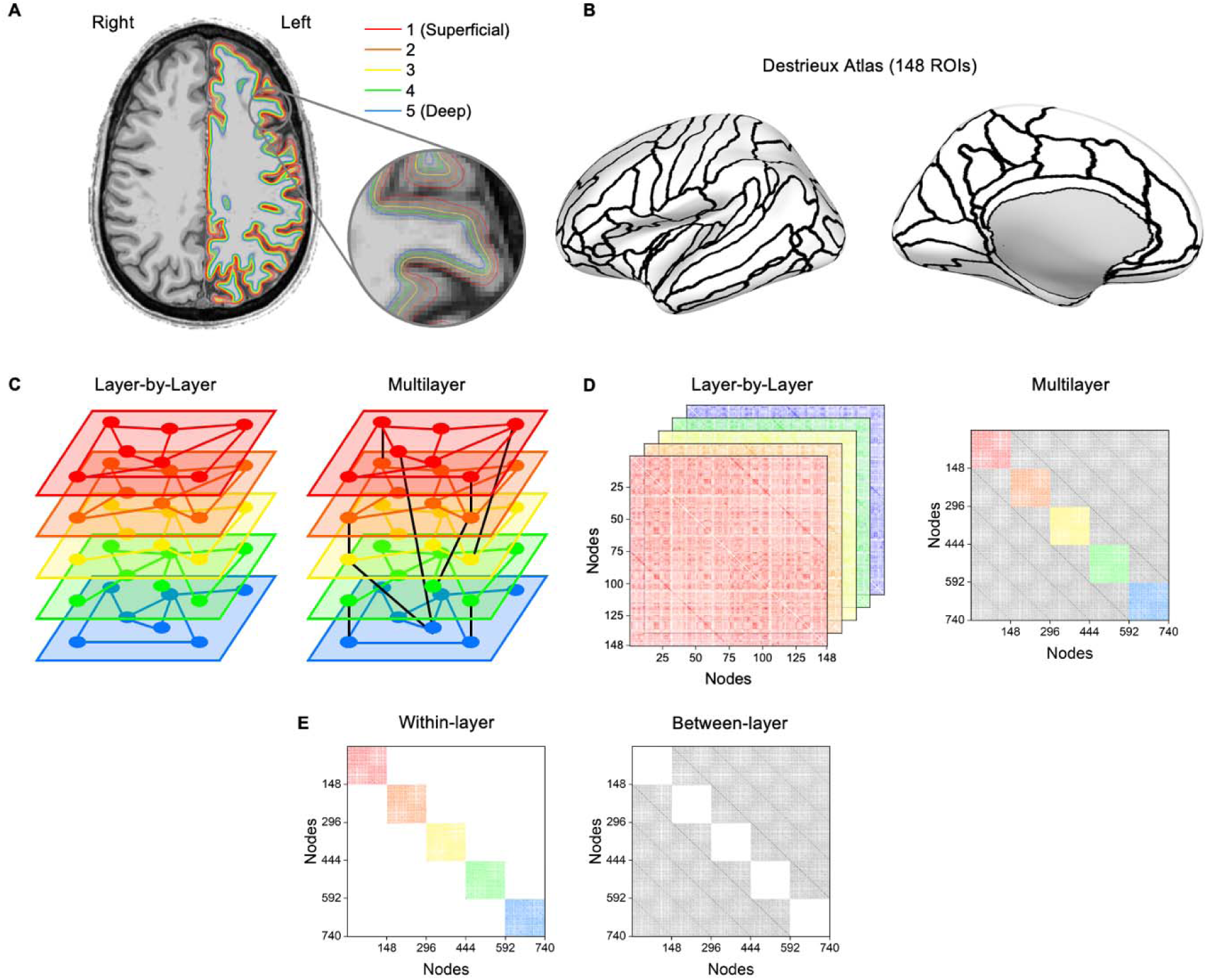
Multilayer connectomic pipeline to analyze functional connectivity across different cortical depths. Here, “layers” refer to the dimensions of the connectivity matrices which represent fMRI signals gathered from different “cortical depths.” **(A)** The cortex was uniformly divided into five surfaces at different depths, as seen above projected on a 0.75-mm isotropic-resolution anatomical T_1_-weighted image. **(B)** The brain was parcellated into 148 regions-of-interest (ROIs) (74 per hemisphere) based on the Destrieux atlas in FreeSurfer (Destrieux et al., 2010; Fischl et al., 2004). The ROIs are shown on an inflated left-hemisphere cortical surface. **(C)** Schematic showing the difference between a layer-by-layer network and a multilayer network. In the layer-by-layer approach, each layer (network) is independent of other layers while in the multilayer approach, the layers are inter-connected. A sparser multilayer network is shown for visualization purposes. **(D)** Example matrix construction from both the layer-by-layer and multilayer approaches. While both approaches use matrices derived from Pearson correlations from the different layers, the multilayer approach generates a supra-adjacency matrix that also has correlations between different layers (shown in grayscale). **(E)** Example matrix construction for within-layer and between-layer matrices. For within-layer matrices, each sub-matrix is extracted individually for analysis. White areas represent connections excluded from the analysis.

### Matrix Generation and Processing

Two parallel matrix processing pipelines were used to generate individual independent adjacency matrices for the layer-by-layer approach while creating one supra-adjacency matrix for the multilayer approach (**Figures 1C and 1D**). The layer-by-layer approach creates an independent network for each cortical depth while the multilayer approach results in five interconnected networks that combines all cortical depths.

One important distinction is between the terminology “depth” and “layer.” Here, depth refers to the anatomical depth in the cortex while layer refers to a specific network derived from a cortical depth. This distinction is critical to avoid equating a network layer with an associated cerebral cortical layer.

The brain was parcellated into 148 regions-of-interest (ROIs) (74 per hemisphere) based on the Destrieux atlas in FreeSurfer (Destrieux et al., 2010; Fischl et al., 2004) (**Figure 1B**). A detailed list of parcellations can be found in the **Table S1**.

Resting-state time series for each participant were concatenated across runs, leading to the following number of time points: 2028 for *n* = 23 participants; 1690 time points: *n* = 1; 1859 time points: *n* = 1; 2197 time points: *n* = 2; 2366 time points: *n* = 2, and 3042 time points: *n* = 1. Concatenated time series were detrended and filtered using a second-order Butterworth filter [high-pass: 0.01 Hz, low-pass: 0.1 Hz, MATLAB-filtfilt].

For the **layer-by-layer approach**, Pearson correlations were derived between ROIs within the same depth, resulting in 10878 pairwise correlations from 148 nodes (ROIs) after removing 148 self-correlations (number of correlations = (nodes^2^ – diagonal nodes) / 2). Pearson correlation coefficients were normalized using Fisher’s z-transformation resulting in five 148-by-148 symmetric weighted connectivity matrices for each participant, i.e., one matrix for each cortical depth (**Figure 1D**). Pearson correlation was used as opposed to partial correlations since partial correlations have been shown to perform poorly in networks with a large number of ROIs (Smith et al., 2011), the larger impact of noise and time series length on partial correlations (Liegeois et al., 2020; Matkovic et al., 2023), and the popularity of Pearson correlations in previous studies (Casas-Roma et al., 2022; Wang et al., 2014).

For the **multilayer approach**, Pearson correlations were derived between ROIs between and within all depths, resulting in 273430 pairwise correlations from 740 nodes (140 ROIs times 5 depths) after removing 740 self-correlations. Pearson correlation coefficients were then normalized using Fisher’s z-transformation with the final product being a 740 by 740 symmetric weighted connectivity matrix for each participant (**Figure 1D**).

For both approaches, individual matrices were normalized and thresholded at 2% intervals ranging from 2 to 40% graph density (ratio of edges present to total number of possible edges) to understand measure differences over a wide range of thresholds. Thresholding is required to minimize the effect spurious correlations and consider only positive correlations.

We also examined the account of within- and between-layer connections only in the context of the complete multilayer matrix. To this end, we draw two additional types of sub-matrices from the multilayer matrix, selectively concentrating on either their within-layer aspect (here, termed **multilayer within-layer**) or the between-layer aspects (termed **multilayer between-layer**) only. The **multilayer within-layer** matrices were derived by normalizing the supra-adjacency matrix, thresholding the matrix, and then extracting the nodes included in each individual layer (i.e. nodes 1 to 148 for layer 1), creating a 148-by-148 weighted connectivity matrix. The **multilayer between-layer** matrices were, in turn, derived by normalizing the supra-adjacency matrix, thresholding the matrix, and then zeroing the five diagonal matrices (from each cortical depth) composing within-layer connections, thus resulting in only between-layer connections. (It is worth noting that since the within-layer and between-layer connectivity matrices were extracted after thresholding, analysis that requires normalization, i.e., non-thresholded matrices, could not be conducted in the context of this analysis; **Figure 1E**.)

### Edge Consistency and Variability

Connections (edges) within and between layers were explored to understand edge consistency and variability between participants. Edge consistency (Finn et al., 2015) was calculated by selecting the top five percent of edges with the lowest standard deviation in un-thresholded multilayer networks. In contrast, edge variability (Menon & Krishnamurthy, 2019) was calculated by selected the top five percent of edges with the highest standard deviation across participants. In both cases, edges in each layer were then summed and divided by the total number of significant edges (edges in the top five percent) to identify the percentage of significant edges in each layer.

### Matrix Similarity

Matrix similarity was used to understand how matrices differed across layers. Thresholded (2– 40%) and normalized matrices were compared using cosine similarity,

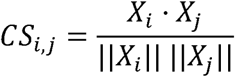

where X_i_ and X_j_ are vectors of the upper triangular elements of two adjacency matrices [MATLAB-pdist2], with values ranging from −1 (maximal dissimilarity) to +1 (maximal similarity). Cosine similarity was shown to distinguish between matrices better than traditionally used Pearson correlation (Cabral et al., 2017; Menon & Krishnamurthy, 2019). Using both the layer-by-layer approach and within-layer matrices, each layer was compared to the other layers. Additionally, to understand how matrix generation differs between methods, the same layer was compared across layer-by-layer and within-layer approaches.

### Connectomic Analysis

Global and nodal measures were calculated in MATLAB using the Brain Connectivity Toolbox (MATLAB Version R2022b) (Rubinov & Sporns, 2010) on the Massachusetts Life Sciences Center Compute Cluster (DELL R440 servers with two Intel Xeon Silver 4214R twelve core CPUs). Global measures characterize the entire network while nodal measures characterize attributes of specific node (ROI). Nodal measures can also be averaged to create a global measure. Measures can be grouped into four general categories to describe their overall network characterization: composition, centrality, integration, and segregation. **Composition** measures describe the topology of the network while **centrality** measures detail specific nodal importance for network function. **Integration** measures examine how information flows through the network and **segregation** measures explore how the network is divided into functional components. Therefore, different measures can be used to understand different characteristics of the network. For example, decreased nodal and global average strength was found in maltreated children indicating decreased overall brain connectivity (Puetz et al., 2017) while decreased clustering coefficient and global efficiency in patients with Parkinson’s disease can signify deficits in brain network integration and segregation (Schill et al., 2023). **Table 1** denotes the measures used in this work, and detailed explanation of each measure can be found in Rubinov and Sporns (Rubinov & Sporns, 2010). Additionally, small-worldness, a global quantifier that examines how “random” a network is organized, was also calculated on layer-by-layer networks since noisier data will appear more “random” (Humphries & Gurney, 2008). Small-worldness was not calculated on within-layer and between-layer connectivity matrices since the calculation requires normalization.

**Table 1.** List of network measures used in this work organized by functional category. (n) denotes a nodal measures.

|  |  |  |  |
| --- | --- | --- | --- |
| Composition | Centrality | Integration | Segregation |
| - Largest Cluster Size | - Betweenness | - Characteristic Path | - Maximized |
| - Graph Density | Centrality (n) | Length | Modularity Q |
| - Degree (n) | - Eigenvector | - Radius | - Transitivity |
| - Strength (n) | Centrality (n) | - Diameter | - Clustering |
|  | - Participation | - Global Efficiency | Coefficient (n) |
|  | Coefficient (n) | - Assortativity | - Local Efficiency (n) |

### Statistical Analysis

To avoid the bias of selecting a single threshold, area-under-the-curve (AUC) analysis was conducted to create a threshold-independent measure. The measures in **Table 1** were calculated at each threshold (from 2 to 40% graph density in 2% intervals). The measure values at each threshold were then plotted against their threshold, and the area underneath the generated curve was calculated using a trapezoidal integration method [MATLAB-trapz].

For each global measure (and averaged nodal measure), the AUC value for each layer for all participants was compared using a one-way analysis of variance (ANOVA) [MATLAB-anova1]. A one-way ANOVA was also used to compare each nodal measure to find differences at each specific ROI (node). For the multilayer network and between-layer measures, only nodal values (both averaged and individual) were compared since global measures for the multilayer network (and thus between-layer measures) contain effects from every layer. For global values, a False Discovery Rate (FDR, Benjamini-Hochberg) correction (alpha = 0.05) was applied to account for multiple comparisons (Benjamini & Hochberg, 1995; Groppe, 2024). Additionally, for nodal values (non-averaged), a FRD (Bonferroni-Holm method) correction (alpha = 0.01) was applied to account for multiple comparisons (Groppe, 2023; Holm, 1979).

### Cortical Thickness Validation

One potential confounding factor using whole-brain laminar analysis is that different brain regions have different cortical thicknesses (Barbas, 2015; Ding et al., 2009; Zachlod et al., 2020). Thus, comparing cortical thickness values of significant brain regions (defined above) can help evaluate whether our findings may be influenced by cortical thickness. Subsequently, cortical thickness values for each ROI for each participant were extracted using FreeSurfer and averaged across all subjects (Fischl, 2012). The distribution of significant nodes versus non-significant nodes for each nodal measure and pipeline with greater than ten significant nodes were compared using a t-test [MATLAB-ttest2].

## Results

### Matrix Similarity and Edge Comparison

Edges from the multilayer matrix were compared to understand differences and similarities between participants and to see if the laminar connectomic methodology can distinguish different participants. High edge consistency indicates a similar connectivity pattern between participants while a high edge variability increases the ability to distinguish been participants. **Figure 2A** and **Table 2** shows the percentage of consistent edge strengths between participants that each layer contains from the multilayer matrix. Layer 1 (derived from the depth closest to pial surface) has the largest number of consistent edges (29.5*2%*) with connections between layer 1 and layer 5 (7.06%) being the most consistent between participants. In contrast, **Figure 2B** shows the edge variability between participants with the highest variability found in layer 5 (closest to white matter) overall (33.65%) and in within layer connections (11.86%). It is important to clarify that edge consistency and variability are not mutually related, even though they provide complementary results above.

**Figure 2.**
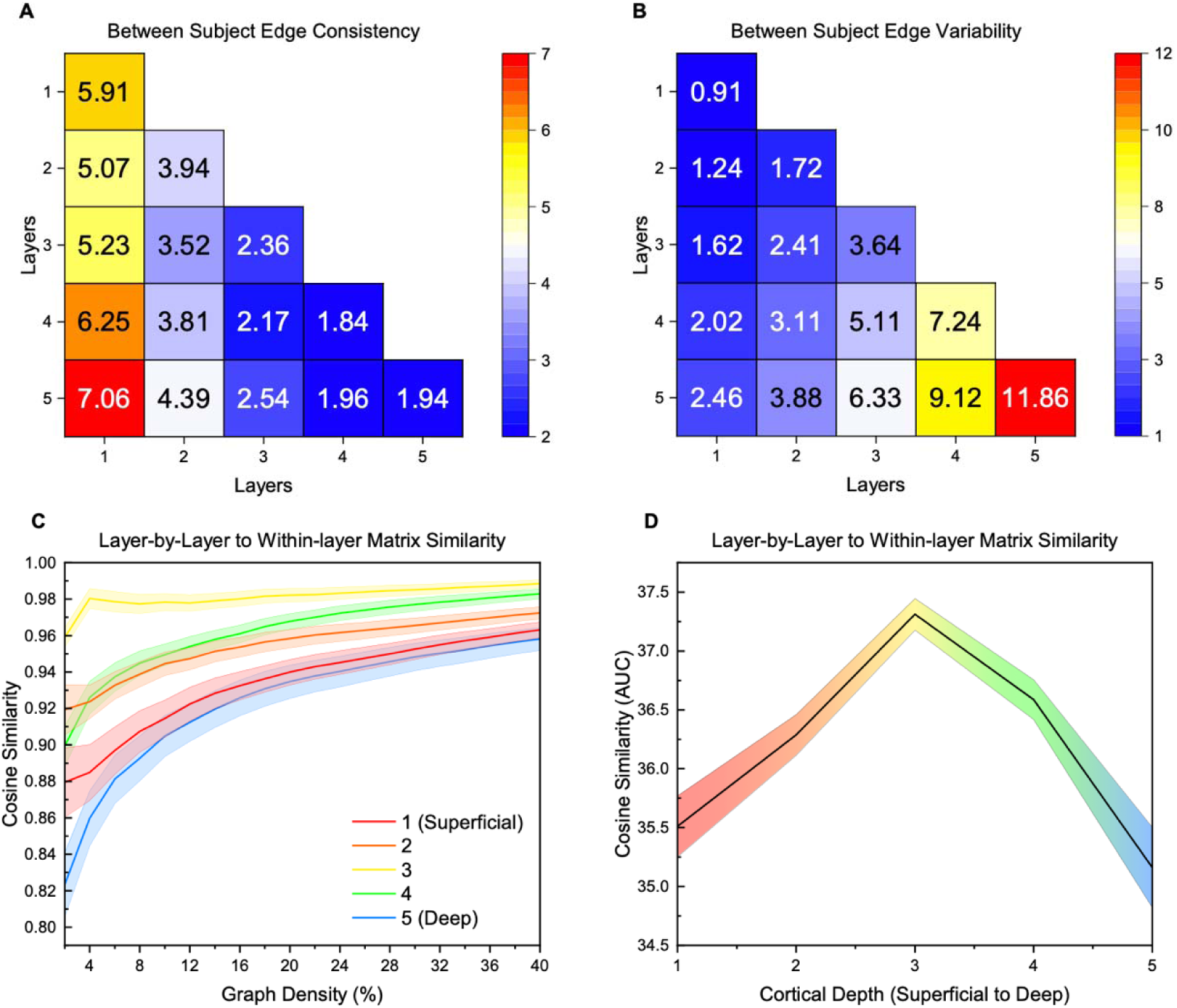
**(A)** Edge consistency between each participant (multilayer matrix). The edges of each layer of the multilayer matrix were compared to find the edge strengths that had the lowest 5% standard deviation between participants. Higher values indicate a higher percentage of consistent edges, indicating consistent features between participants for those connections. **(B)** Edge variability between each participant (multilayer matrix). The edges of each layer of the multilayer matrix were compared to find the edge strengths that had the highest 5% standard deviation between participants. Higher values indicate a higher percentage of variable edges, indicating variable features between participants for those connections. **(C)** Cosine similarity between layer-by-layer and within-layer matrices (multilayer approach). Within participant, the matrix generation methods were compared using cosine similarity across a range of thresholds at each layer. Cosine similarity values range from −1 (maximal dissimilarity) to +1 (maximal similarity). The mean value at each threshold is plotted while the shaded region indicates the standard error. **(D)** Area-under-the-curve (AUC) measure in comparing layer-by-layer and within-layer matrix generation methods. Linear interpolation was used for visualization. The AUC from **(C)** is calculated using trapezoidal approximation. Higher values indicate higher similarity between methods while lower values indicate lower similarity between methods. The mean AUC value at each layer is plotted while the shaded region indicates the standard error.

**Table 2.** Edge consistency and variability percentages for each layer derived from the multilayer matrix. Bolded values show the highest percentage for each measure.

| Layer | Edge Consistency (%) | Edge Variability (%) |
| --- | --- | --- |
| 1 (Superficial) | **29.52** | 8.25 |
| 2 | 20.73 | 12.36 |
| 3 | 15.82 | 19.11 |
| 4 | 16.03 | 26.60 |
| 5 (Deep) | 17.89 | **33.65** |

After edge analysis, connectivity matrices were compared within participants to understand the two competing matrix generation methodologies: layer-by-layer vs. within-layer (multilayer) approach. Cosine similarity was used to examine similarities between different connectivity matrices. In comparing within participant matrices across layers (**Figure S4**), layers were found to be similar with the most distant layers (layer 1 to layer 5) having the lowest similarity in both layer-by-layer matrices and within-layer matrices. Matrices within the same layer and within participant were also compared across the matrix generation methods (**Figure 2C and 2D**). Layer 3 was the most consistent across the two methodologies while the peripheral layers (layer 1 and layer 5) differed the most between methods.

### Single Layer Results

#### Global

Global network measures were calculated for layer-by-layer matrices. AUC values for each global measure can be found summarized in **Table S2**. Network diameter (*p* = 0.049) significantly increased from the most superficial layer 1 to the deepest layer 5 (**Figure 3A**). Largest cluster size (*p* = 0.0024), average betweenness centrality (*p* = 0.028), average local efficiency (*p* = 0.046), and eigenvector centrality (p = 0.028) significantly decreased from layer 1 to layer 5 (**Figure 3B–E**), with a peak in layer 2 for average betweenness centrality and average local efficiency. There were no significant differences in modularity, transitivity, characteristic path length, global efficiency, radius, assortativity, average degree centrality, average strength, average clustering coefficient, and average participation coefficient (**Figure S5–6).** Graph density and average degree centrality were constant across layers due to both measures being a direct function of thresholding (**Figure S5–6**). Additionally, small-worldness showed a general trend of decreasing with depth; however, there was no significant differences between layers (*p* = 0.2651) and small-worldness was greater than one (indicating a small-world network) for all thresholds except 40% graph density (**Figure S7**).

**Figure 3.**
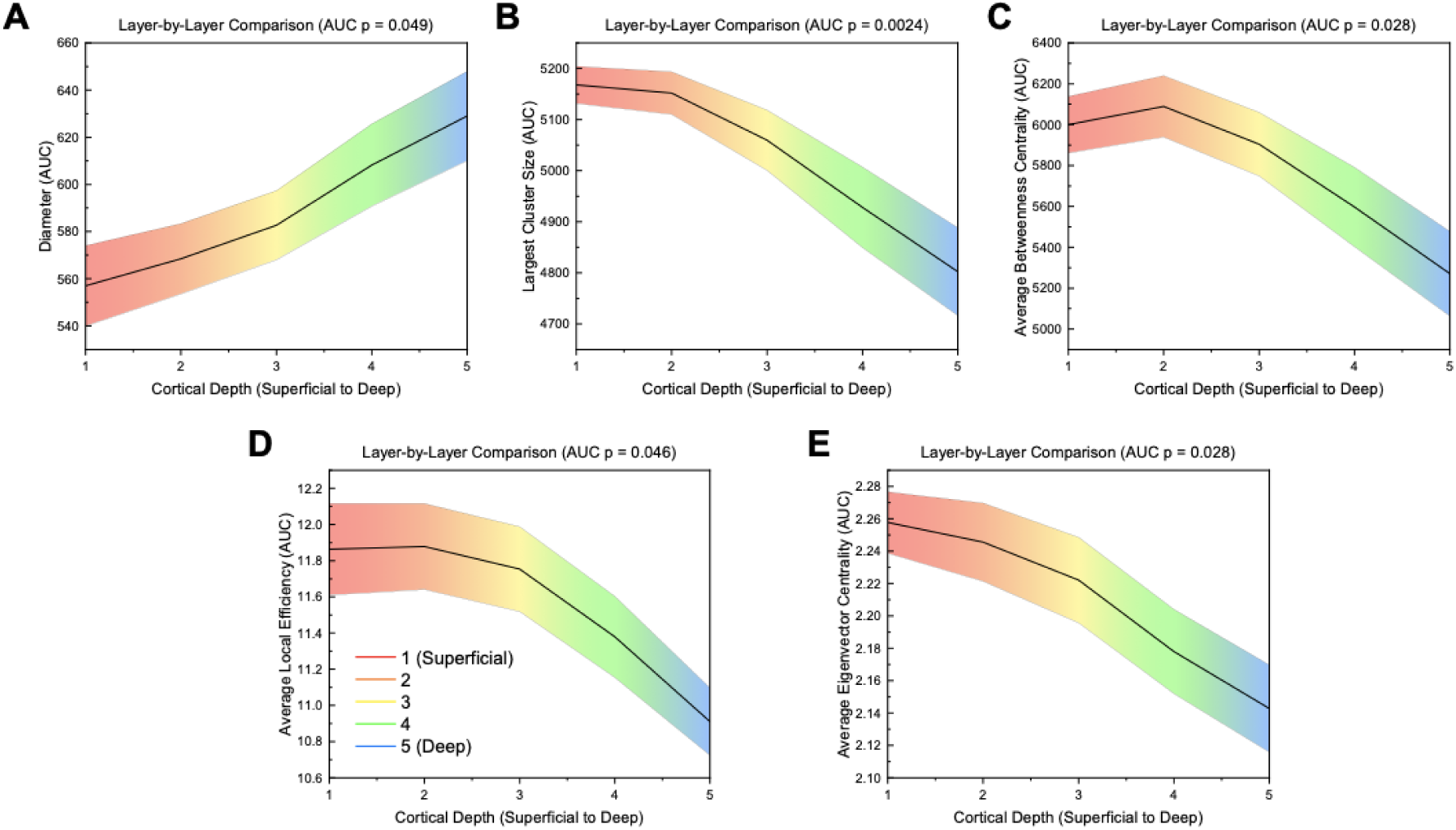
Area-under-the-curve (AUC) values across different layers for significant global measures (*p* ≤ 0.05) for layer-by-layer analysis. Significance was calculated using a one-way ANOVA with an FDR correction (alpha = 0.05). Linear interpolation was used for visualization. The mean value across participants at each layer is plotted while the shaded region indicates the standard error. P values shown are FDR corrected (Benjamini-Hochberg method, alpha = 0.05).

#### Nodal

**Table 3** shows the number of nodes in brain regions in layer-by-layer matrices with significant differences between layers (FDR correction with alpha = 0.01) (See **Table S3** for specific values and regions). Degree centrality, strength, and eigenvector centrality had the greatest number of significant nodes (4/148) (**Table 3**). The limbic region had more significant nodes than all other regions for each measure, except for clustering coefficient which was tied with the temporal region (one significant node for each region). In all measures, the right hemisphere had more significant nodes than the left hemisphere (**Table 3**). In general, the most superficial layers (1 and 2) had the highest value for significant nodes (**Table 3, TableS3, Figure S8–S9**). Significant nodes were distributed across node thickness levels (**Figure S10–S11**).

**Table 3.** Number of significant nodes within each brain region for layer-by-layer analysis. Significance was calculated from the area-under-the curve (AUC) values using a one-way ANOVA with an FDR correction (alpha = 0.01) to account for multiple comparisons (Groppe, 2023; Holm, 1979). Details of nodal mapping to each region can be found in Table S1. Orange: hemisphere with the highest number of nodes; Yellow: measure with the highest number of nodes; Green: region within each measure with the highest number of nodes.

| Network Measure | Hemisphere | All Regions | Frontal | Limbic | Occipital | Parietal | Temporal |
| --- | --- | --- | --- | --- | --- | --- | --- |
| Degree Centrality | Left: | 1/74 | 0/24 | 1/14 | 0/10 | 0/10 | 0/16 |
|  | Right: | 3/74 | 0/24 | 2/14 | 0/10 | 0/10 | 1/16 |
|  | Both: | 4/148 | 0/48 | 3/28 | 0/20 | 0/20 | 1/32 |
| Strength | Left: | 1/74 | 0/24 | 1/14 | 0/10 | 0/10 | 0/16 |
|  | Right: | 3/74 | 0/24 | 2/14 | 0/10 | 0/10 | 1/16 |
|  | Both: | 4/148 | 0/48 | 3/28 | 0/20 | 0/20 | 1/32 |
| Eigenvector Centrality | Left: | 1/74 | 0/24 | 1/14 | 0/10 | 0/10 | 0/16 |
|  | Right: | 3/74 | 0/24 | 2/14 | 0/10 | 0/10 | 1/16 |
|  | Both: | 4/148 | 0/48 | 3/28 | 0/20 | 0/20 | 1/32 |
| Betweenness | Left: | 0/74 | 0/24 | 0/14 | 0/10 | 0/10 | 0/16 |
| Centrality | Right: | 3/74 | 1/24 | 2/14 | 0/10 | 0/10 | 0/16 |
|  | Both: | 3/148 | 1/48 | 2/28 | 0/20 | 0/20 | 0/32 |
| Clustering | Left: | 0/74 | 0/24 | 0/14 | 0/10 | 0/10 | 0/16 |
| Coefficient | Right: | 2/74 | 0/24 | 1/14 | 0/10 | 0/10 | 1/16 |
|  | Both: | 2/148 | 0/48 | 1/28 | 0/20 | 0/20 | 1/32 |
| Local | Left: | 1/74 | 0/24 | 1/14 | 0/10 | 0/10 | 0/16 |
| Efficiency | Right: | 3/74 | 0/24 | 2/14 | 0/10 | 0/10 | 1/16 |
|  | Both: | 4/148 | 0/48 | 3/28 | 0/20 | 0/20 | 1/32 |
| Participation | Left: | 1/74 | 0/24 | 1/14 | 0/10 | 0/10 | 0/16 |
| Coefficient | Right: | 2/74 | 0/24 | 2/14 | 0/10 | 0/10 | 0/16 |
|  | Both: | 3/148 | 0/48 | 3/28 | 0/20 | 0/20 | 0/32 |

### Multilayer Results

#### Within-layer Global

**Figure 4** shows global network measures calculated for within-layer matrices. AUC values for each global measure can be found summarized in **Table S4**. Characteristic path length (*p* = 0.014) and diameter (*p* < 0.001) all increased from layer 1 to layer 5 (**Figure 4A–B**). Largest cluster size (*p* < 0.001), graph density (*p* < 0.001), average degree centrality (p < 0.001), average strength (*p* < 0.001), average eigenvector centrality (*p* < 0.001), and average participation coefficient (*p* = 0.0011) significantly decreased with cortical depth (layer 1 to 5) (**Figure 4C–H**). There were no significant differences for modularity, transitivity, global efficiency, radius, assortativity, average betweenness centrality, average clustering coefficient, and average local efficiency (**Figure S12–13**). In contrast to layer-by-layer results, graph density and average degree centrality were different across layers due to the within-layer matrix generation methodology allowing each individual layer to have a different graph density.

**Figure 4.**
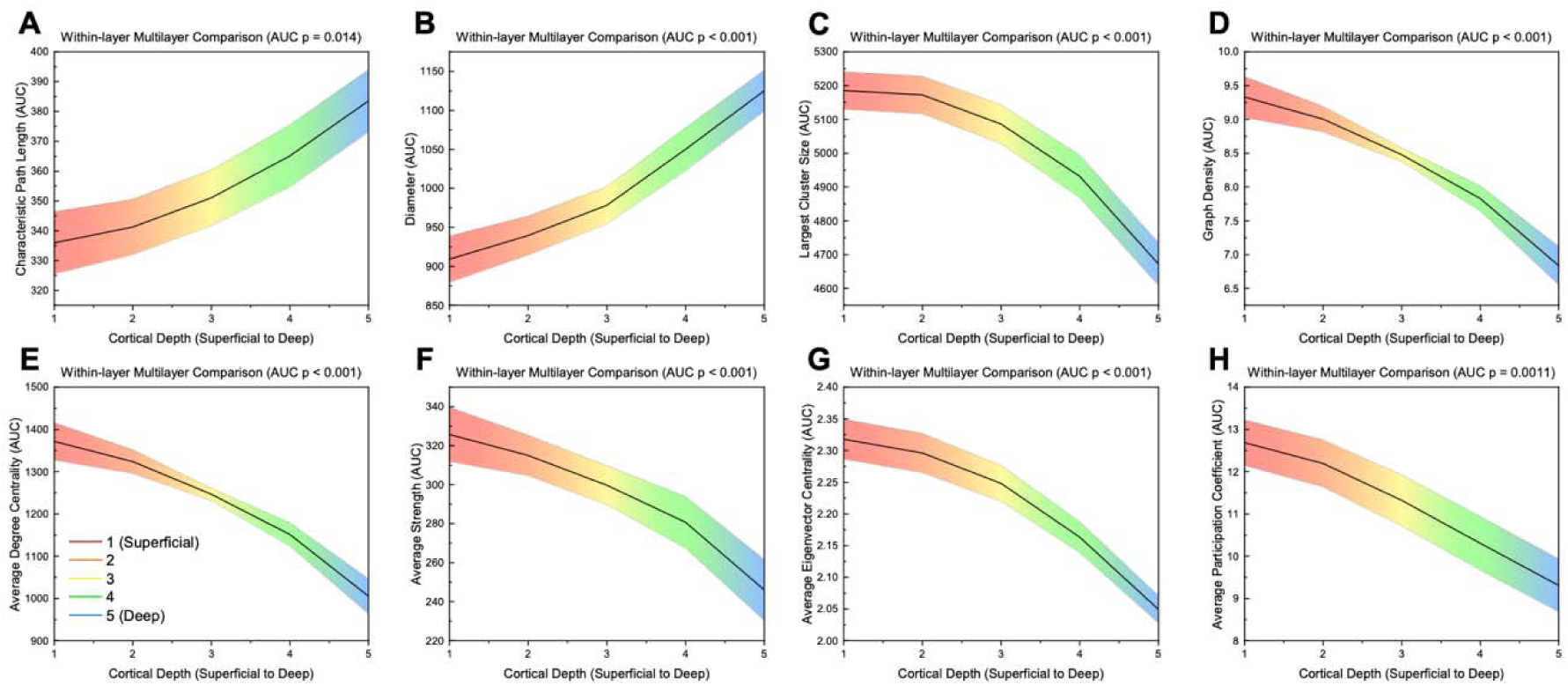
Area-under-the-curve (AUC) values across different layers for significant global measures (*p* ≤ 0.05) for within-layer analysis. Significance was calculated using a one-way ANOVA with an FDR correction (alpha = 0.05). Linear interpolation was used for visualization. The mean value across participants at each layer is plotted while the shaded region indicates the standard error. P values shown are FDR corrected (Benjamini-Hochberg method, alpha = 0.05).

#### Within-layer Local

The number of nodes in brain regions with significant differences using within-layer matrices can be found in **Table 4** (FDR correction with alpha = 0.01, see **Table S5** for specific values and regions). Degree centrality had the largest number of significant nodes (15/148) followed by strength (9/148), participation coefficient (9/148), and local efficiency (8/148) (**Table 4**). In all measures except participation coefficient, the limbic region had the most significant nodes; in participation coefficient, the temporal region had the most significant nodes (4/32). In all measures, the right hemisphere had more significant nodes than the left hemisphere (**Table 4**). For significant nodes, either layers 1 or 2 had the highest value (**Table 4**, **Table S5, Figure S14– 15).** Significant nodes were spread across different node thicknesses (**Figure S16–S17**).

**Table 4.** Number of significant nodes within each brain region for within-layer analysis. Significance was calculated from the area-under-the curve (AUC) values using a one-way ANOVA with an FDR correction (alpha = 0.01) to account for multiple comparisons (Groppe, 2023; Holm, 1979). Details of nodal mapping to each region can be found in Table S1. Orange: hemisphere with the highest number of nodes; Yellow: measure with the highest number of nodes; Green: region within each measure with the highest number of nodes.

| Network Measure | Hemisphere | All Regions | Frontal | Limbic | Occipital | Parietal | Temporal |
| --- | --- | --- | --- | --- | --- | --- | --- |
| Degree Centrality | Left: | 4/74 | 0/24 | 4/14 | 0/10 | 0/10 | 0/16 |
|  | Right: | 11/74 | 1/24 | 5/14 | 0/10 | 0/10 | 5/16 |
|  | Both: | 15/148 | 1/48 | 9/28 | 0/20 | 0/20 | 5/32 |
| Strength | Left: | 2/74 | 0/24 | 2/14 | 0/10 | 0/10 | 0/16 |
|  | Right: | 7/74 | 0/24 | 3/14 | 0/10 | 0/10 | 4/16 |
|  | Both: | 9/148 | 0/48 | 5/28 | 0/20 | 0/20 | 4/32 |
| Eigenvector | Left: | 1/74 | 0/24 | 1/14 | 0/10 | 0/10 | 0/16 |
| Centrality | Right: | 3/74 | 0/24 | 2/14 | 0/10 | 0/10 | 1/16 |
|  | Both: | 4/148 | 0/48 | 3/28 | 0/20 | 0/20 | 1/32 |
| Betweenness | Left: | 0/74 | 0/24 | 0/14 | 0/10 | 0/10 | 0/16 |
| Centrality | Right: | 3/74 | 1/24 | 2/14 | 0/10 | 0/10 | 0/16 |
|  | Both: | 3/148 | 1/48 | 2/28 | 0/20 | 0/20 | 0/32 |
| Clustering | Left: | 1/74 | 0/24 | 1/14 | 0/10 | 0/10 | 0/16 |
| Coefficient | Right: | 5/74 | 0/24 | 3/14 | 0/10 | 0/10 | 2/16 |
|  | Both: | 6/148 | 0/48 | 4/28 | 0/20 | 0/20 | 2/32 |
| Local | Left: | 1/74 | 0/24 | 1/14 | 0/10 | 0/10 | 0/16 |
| Efficiency | Right: | 7/74 | 0/24 | 4/14 | 0/10 | 0/10 | 3/16 |
|  | Both: | 8/148 | 0/48 | 5/28 | 0/20 | 0/20 | 3/32 |
| Participation | Left: | 2/74 | 0/24 | 1/14 | 0/10 | 0/10 | 1/16 |
| Coefficient | Right: | 7/74 | 1/24 | 2/14 | 1/10 | 0/10 | 3/16 |
|  | Both: | 9/148 | 1/48 | 3/28 | 1/20 | 0/20 | 4/32 |

### Multilayer Global

Alongside measures for individual layers, global measures were calculated for the supra-adjacency matrix created using the multilayer approach. While only global values derived from nodal averages were statistically compared between layers, AUC values for all global measures are summarized in **Table S6**. Average degree centrality (*p* < 0.001), average strength (*p* = 0.0019), and average eigenvector centrality (*p* < 0.001) decreased from layer 1 to layer 5 with a slight peak at layer 2 (**Figure 5A–C**). Average betweenness centrality (*p* < 0.001) and average participation coefficient (*p* = 0.019) also decreased from layer 1 to layer 5 (**Figure 5D–E**). In contrast, average clustering coefficient tended to increase from layer 1 to layer 5 (*p* = 0.080) (**Figure S18–19, Table S6**). Average local efficiency showed no significant difference between layers (**Figure S18–19, Table S6**). Additionally, layer-wise graph density, graph density derived from each individual layer within the multilayer network, decreased from layer 1 to layer 5 with a slight peak at layer 2 (**Figure S20**).

**Figure 5.**
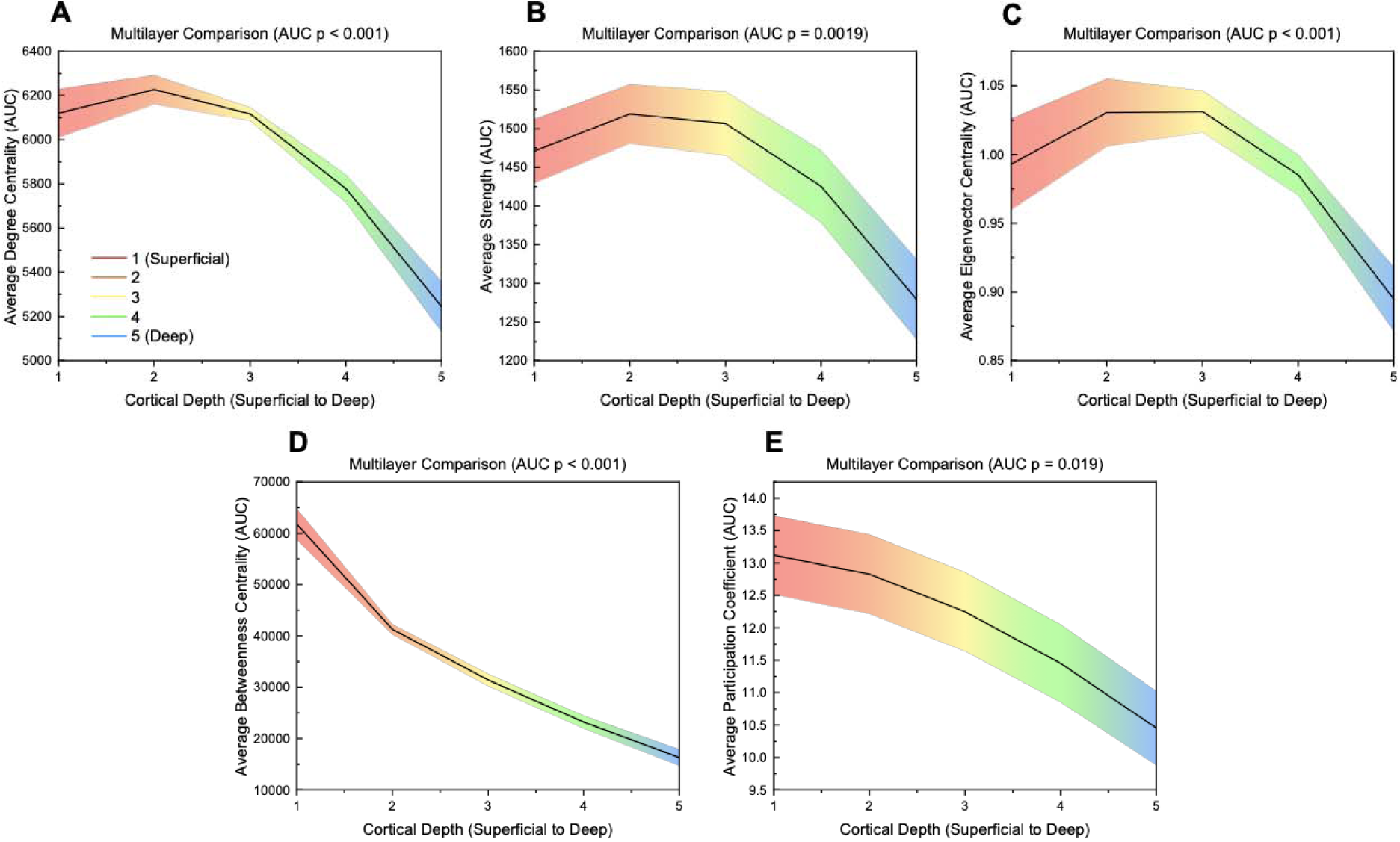
Area-under-the-curve (AUC) values across different layers for significant global measures (*p* ≤ 0.05) for multilayer analysis. Significance was calculated using a one-way ANOVA with an FDR correction (alpha = 0.05). Linear interpolation was used for visualization. The mean value across participants at each layer is plotted while the shaded region indicates the standard error. P values shown are FDR corrected (Benjamini-Hochberg method, alpha = 0.05).

### Multilayer Local

**Table 5** depicts the number of nodes in specific brain regions that were significantly different between layers for the multilayer analysis (FDR correction with alpha = 0.01) (See **Table S7** for specific values and regions). Betweenness centrality identified the most significant nodes (58/148) with 19/48 frontal nodes, 16/28 limbic nodes, 2/20 occipital nodes, 5/20 parietal nodes, and 16/32 temporal nodes (**Table 5**). The limbic region had highest percentage of nodes in all measures except clustering coefficient (4/32 in temporal) and local efficiency (no significant nodes found). Additionally, using betweenness centrality the frontal region had the highest number of significant nodes (19/48). Again, in all measures, the right hemisphere had more nodes with significant differences between layers (**Table 5**). While most significant regions across measures were highest in the superficial layers (layers 1,2), especially in betweenness centrality, the deepest layer (layer 5) had the highest values for nodes significant in clustering coefficient (**Table 5, Table S7, Figure S21**). The thickness of significant nodes was spread across the spectrum of thickness levels, with a preference towards thicker nodes for betweenness centrality (**Figure S22–S23**).

**Table 5.**
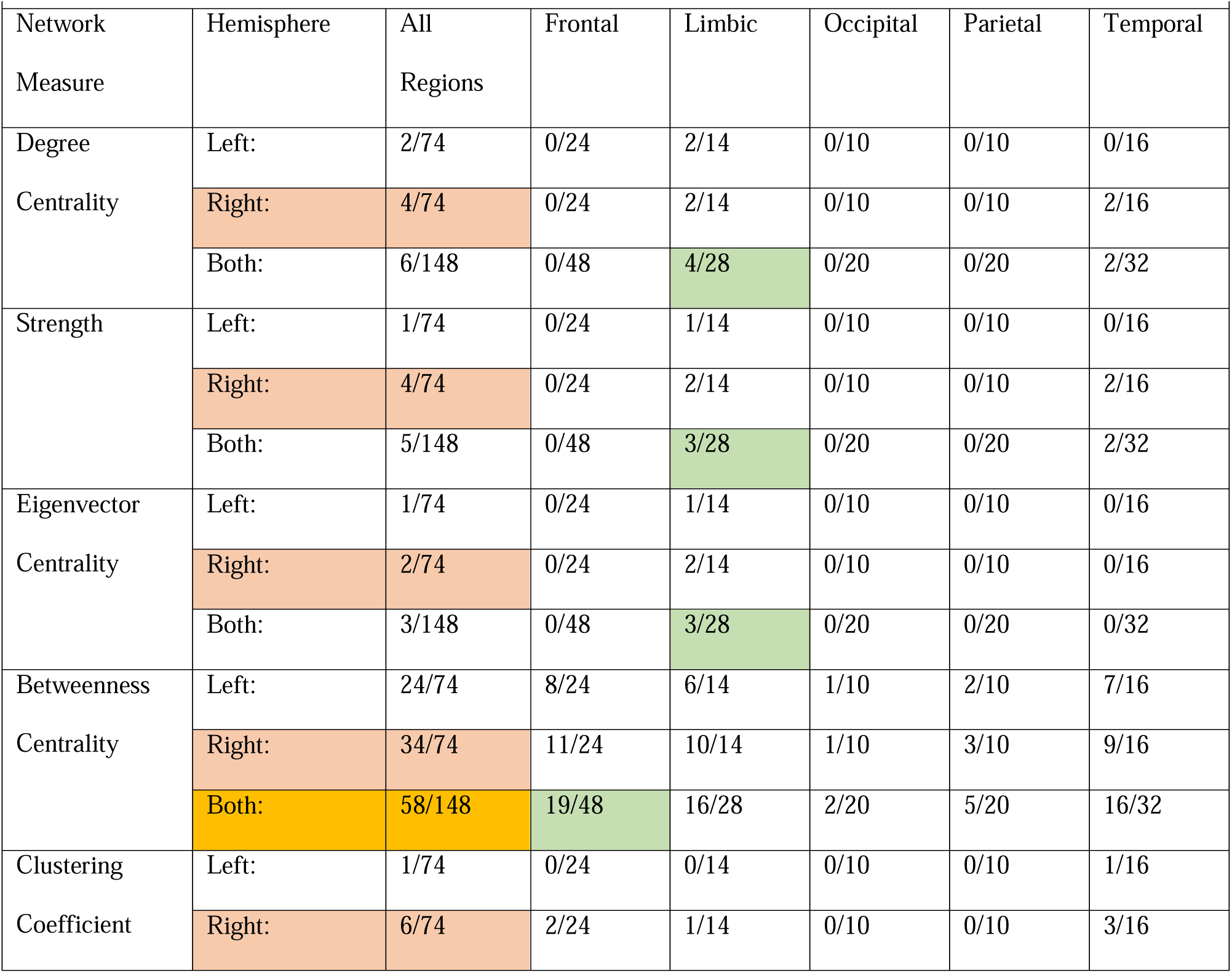
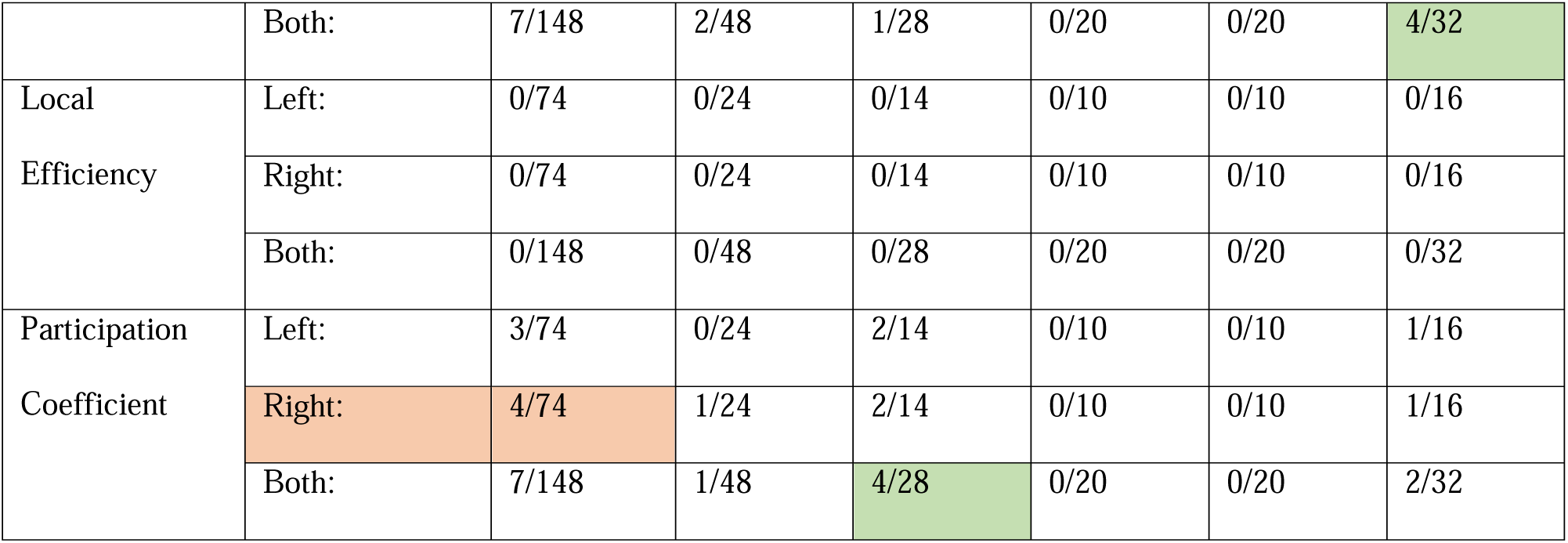
Number of significant nodes within each brain region for multilayer analysis. Significance was calculated from the area-under-the curve (AUC) values using a one-way ANOVA with an FDR correction (alpha = 0.01) to account for multiple comparisons (Groppe, 2023; Holm, 1979). Details of nodal mapping to each region can be found in Table S1. Orange: hemisphere with the highest number of nodes; Yellow: measure with the highest number of nodes; Green: region within each measure with the highest number of nodes.

| Network Measure | Hemisphere | All Regions | Frontal | Limbic | Occipital | Parietal | Temporal |
| --- | --- | --- | --- | --- | --- | --- | --- |
| Degree Centrality | Left: | 2/74 | 0/24 | 2/14 | 0/10 | 0/10 | 0/16 |
|  | Right: | 4/74 | 0/24 | 2/14 | 0/10 | 0/10 | 2/16 |
|  | Both: | 6/148 | 0/48 | 4/28 | 0/20 | 0/20 | 2/32 |
| Strength | Left: | 1/74 | 0/24 | 1/14 | 0/10 | 0/10 | 0/16 |
|  | Right: | 4/74 | 0/24 | 2/14 | 0/10 | 0/10 | 2/16 |
|  | Both: | 5/148 | 0/48 | 3/28 | 0/20 | 0/20 | 2/32 |
| Eigenvector Centrality | Left: | 1/74 | 0/24 | 1/14 | 0/10 | 0/10 | 0/16 |
|  | Right: | 2/74 | 0/24 | 2/14 | 0/10 | 0/10 | 0/16 |
|  | Both: | 3/148 | 0/48 | 3/28 | 0/20 | 0/20 | 0/32 |
| Betweenness Centrality | Left: | 24/74 | 8/24 | 6/14 | 1/10 | 2/10 | 7/16 |
|  | Right: | 34/74 | 11/24 | 10/14 | 1/10 | 3/10 | 9/16 |
|  | Both: | 58/148 | 19/48 | 16/28 | 2/20 | 5/20 | 16/32 |
| Clustering Coefficient | Left: | 1/74 | 0/24 | 0/14 | 0/10 | 0/10 | 1/16 |
|  | Right: | 6/74 | 2/24 | 1/14 | 0/10 | 0/10 | 3/16 |
|  | Both: | 7/148 | 2/48 | 1/28 | 0/20 | 0/20 | 4/32 |
| Local Efficiency | Left: | 0/74 | 0/24 | 0/14 | 0/10 | 0/10 | 0/16 |
|  | Right: | 0/74 | 0/24 | 0/14 | 0/10 | 0/10 | 0/16 |
|  | Both: | 0/148 | 0/48 | 0/28 | 0/20 | 0/20 | 0/32 |
| Participation Coefficient | Left: | 3/74 | 0/24 | 2/14 | 0/10 | 0/10 | 1/16 |
|  | Right: | 4/74 | 1/24 | 2/14 | 0/10 | 0/10 | 1/16 |
|  | Both: | 7/148 | 1/48 | 4/28 | 0/20 | 0/20 | 2/32 |

### Between-layer Global

Significant between-layer global measures can be found in **Figure 6**. AUC values for all global measures can be found in **Table S8**. Average degree centrality (*p* < 0.001), average strength (*p* = 0.0021), average eigenvector centrality (*p* < 0.001), average betweenness centrality (p < 0.001), and average participation coefficient (*p* = 0.032) decreased from superficial (layer 1) to deep (layer 5) with all except average participation coefficient showing a peak in layers 2 and 3 (**Figure 6A–D, F**). Average clustering coefficient (*p* = 0.0033) increased from layer 1 to layer 5 with a slight decrease from layer 1 to layer 2 (**Figure 6E**). Average local efficiency was the only measure that showed no significant difference between layers (**Figure S24–25, Table S8**).

**Figure 6.**
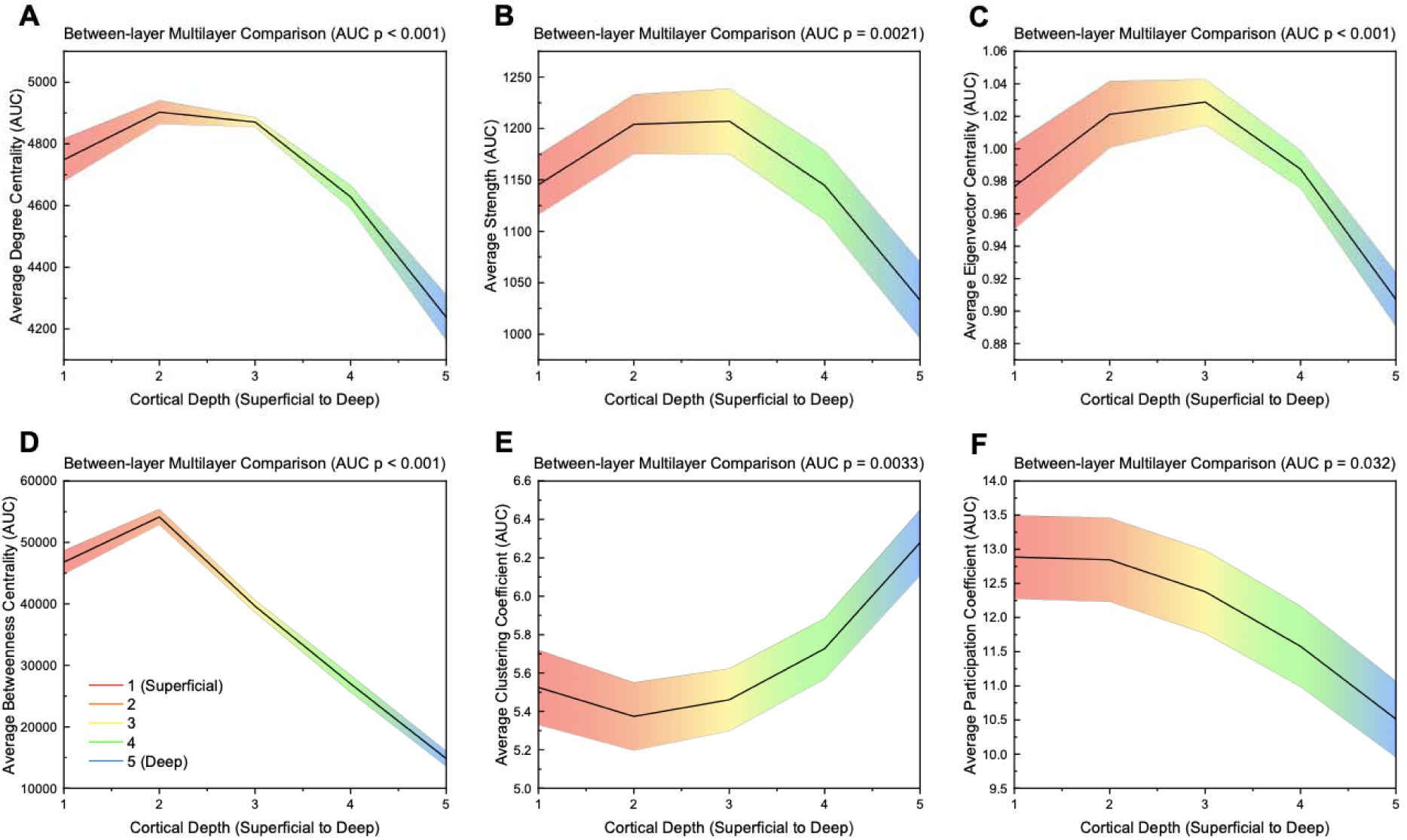
Area-under-the-curve (AUC) values across different layers for significant global measures (*p* ≤ 0.05) for between-layer analysis. Significance was calculated using a one-way ANOVA with an FDR correction (alpha = 0.05). Linear interpolation was used for visualization. The mean value across participants at each layer is plotted while the shaded region indicates the standard error. P values shown are FDR corrected (Benjamini-Hochberg method, alpha = 0.05).

### Between-layer Local

Brain regions with nodes that were significantly different using between-layer matrices can be found in **Table 6** (FDR correction with alpha = 0.01) (See **Table S9** for specific values and regions). Betweenness centrality had the largest number of significant nodes (57/148) followed by clustering coefficient (22/148) (**Table 6**). For betweenness centrality, most significant nodes had the highest values in the superficial layers (layers 1 and 2) while for clustering coefficient, all significant nodes were highest in layer 5 (**Figure S26**). In degree centrality, strength, eigenvector centrality, and betweenness centrality, the limbic region had the highest percentage of significant nodes within each region (**Table 6**). In betweenness centrality, frontal region nodes had the highest absolute number of significant nodes (20/48). In clustering coefficient (10/32) and participation coefficient (4/32), the temporal region had the most significant nodes (**Table 6**). Local efficiency had no significant nodes (**Table 6, Table S9, Figure S26**). The right hemisphere had more significant nodes than the left hemisphere for all measures (**Table 6**). Significant nodes were dispersed across different thickness levels, with betweenness centrality nodes leaning slightly more toward thicker regions (**Figure S27–S28**).

**Table 6.**
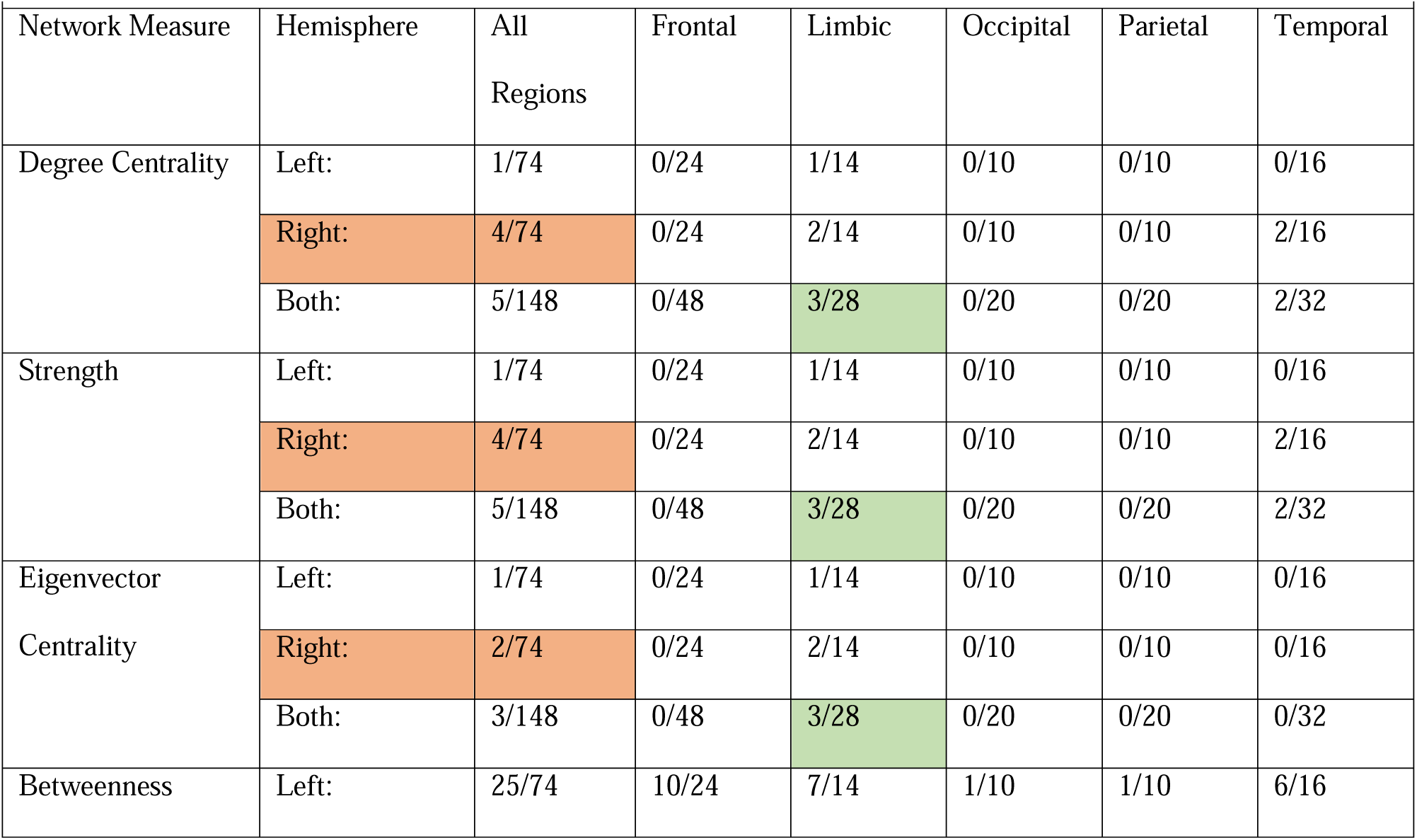
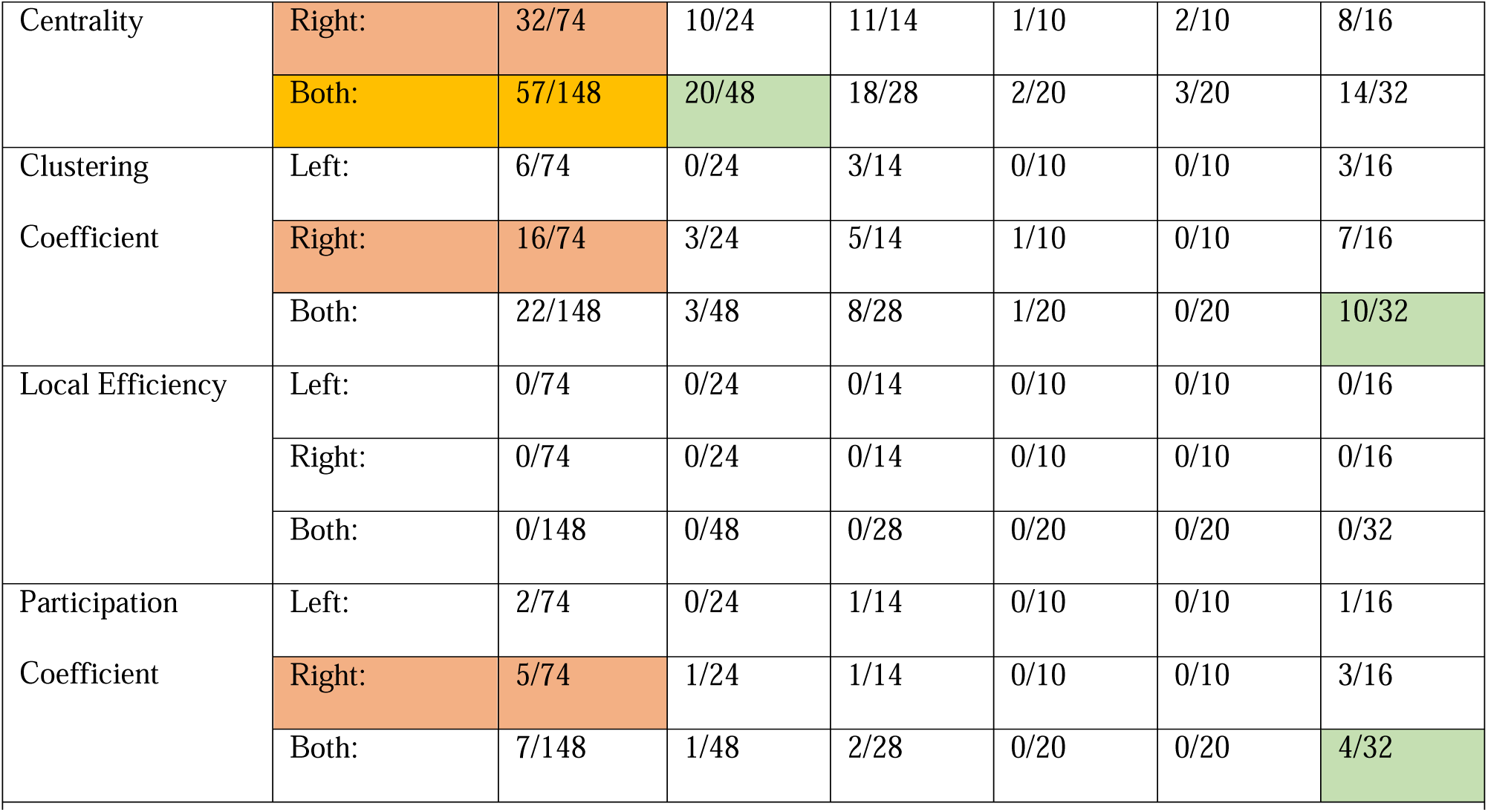
Number of significant nodes within each brain region for between analysis. Significance was calculated from the area-under-the curve (AUC) values using a one-way ANOVA with an FDR correction (alpha = 0.01) to account for multiple comparisons (Groppe, 2023; Holm, 1979). Details of nodal mapping to each region can be found in Table S1. Orange: hemisphere with the highest number of nodes; Yellow: measure with the highest number of nodes; Green: region within each measure with the highest number of nodes.

| Network Measure | Hemisphere | All Regions | Frontal | Limbic | Occipital | Parietal | Temporal |
| --- | --- | --- | --- | --- | --- | --- | --- |
| Degree Centrality | Left: | 1/74 | 0/24 | 1/14 | 0/10 | 0/10 | 0/16 |
|  | Right: | 4/74 | 0/24 | 2/14 | 0/10 | 0/10 | 2/16 |
|  | Both: | 5/148 | 0/48 | 3/28 | 0/20 | 0/20 | 2/32 |
| Strength | Left: | 1/74 | 0/24 | 1/14 | 0/10 | 0/10 | 0/16 |
|  | Right: | 4/74 | 0/24 | 2/14 | 0/10 | 0/10 | 2/16 |
|  | Both: | 5/148 | 0/48 | 3/28 | 0/20 | 0/20 | 2/32 |
| Eigenvector Centrality | Left: | 1/74 | 0/24 | 1/14 | 0/10 | 0/10 | 0/16 |
|  | Right: | 2/74 | 0/24 | 2/14 | 0/10 | 0/10 | 0/16 |
|  | Both: | 3/148 | 0/48 | 3/28 | 0/20 | 0/20 | 0/32 |
| Betweenness | Left: | 25/74 | 10/24 | 7/14 | 1/10 | 1/10 | 6/16 |
| Centrality | Right: | 32/74 | 10/24 | 11/14 | 1/10 | 2/10 | 8/16 |
|  | Both: | 57/148 | 20/48 | 18/28 | 2/20 | 3/20 | 14/32 |
| Clustering | Left: | 6/74 | 0/24 | 3/14 | 0/10 | 0/10 | 3/16 |
| Coefficient | Right: | 16/74 | 3/24 | 5/14 | 1/10 | 0/10 | 7/16 |
|  | Both: | 22/148 | 3/48 | 8/28 | 1/20 | 0/20 | 10/32 |
| Local Efficiency | Left: | 0/74 | 0/24 | 0/14 | 0/10 | 0/10 | 0/16 |
|  | Right: | 0/74 | 0/24 | 0/14 | 0/10 | 0/10 | 0/16 |
|  | Both: | 0/148 | 0/48 | 0/28 | 0/20 | 0/20 | 0/32 |
| Participation | Left: | 2/74 | 0/24 | 1/14 | 0/10 | 0/10 | 1/16 |
| Coefficient | Right: | 5/74 | 1/24 | 1/14 | 0/10 | 0/10 | 3/16 |
|  | Both: | 7/148 | 1/48 | 2/28 | 0/20 | 0/20 | 4/32 |

### Method Comparison

All four network measure methods (layer-by-layer, within-layer, multilayer, between-layer) identified global differences between layers (**Figure 7**). Layer-by-layer and within-layer methods showed an increase in an integration-based global measure (diameter) from layer 1 to layer 5; however, the within-layer approach identified an increase in characteristic path length as well. Similarly, layer-by-layer and within-layer approaches identified a decrease in largest cluster size from layer 1 to layer 5, with an additional decrease found in graph density using the within-layer method.

**Figure 7.**
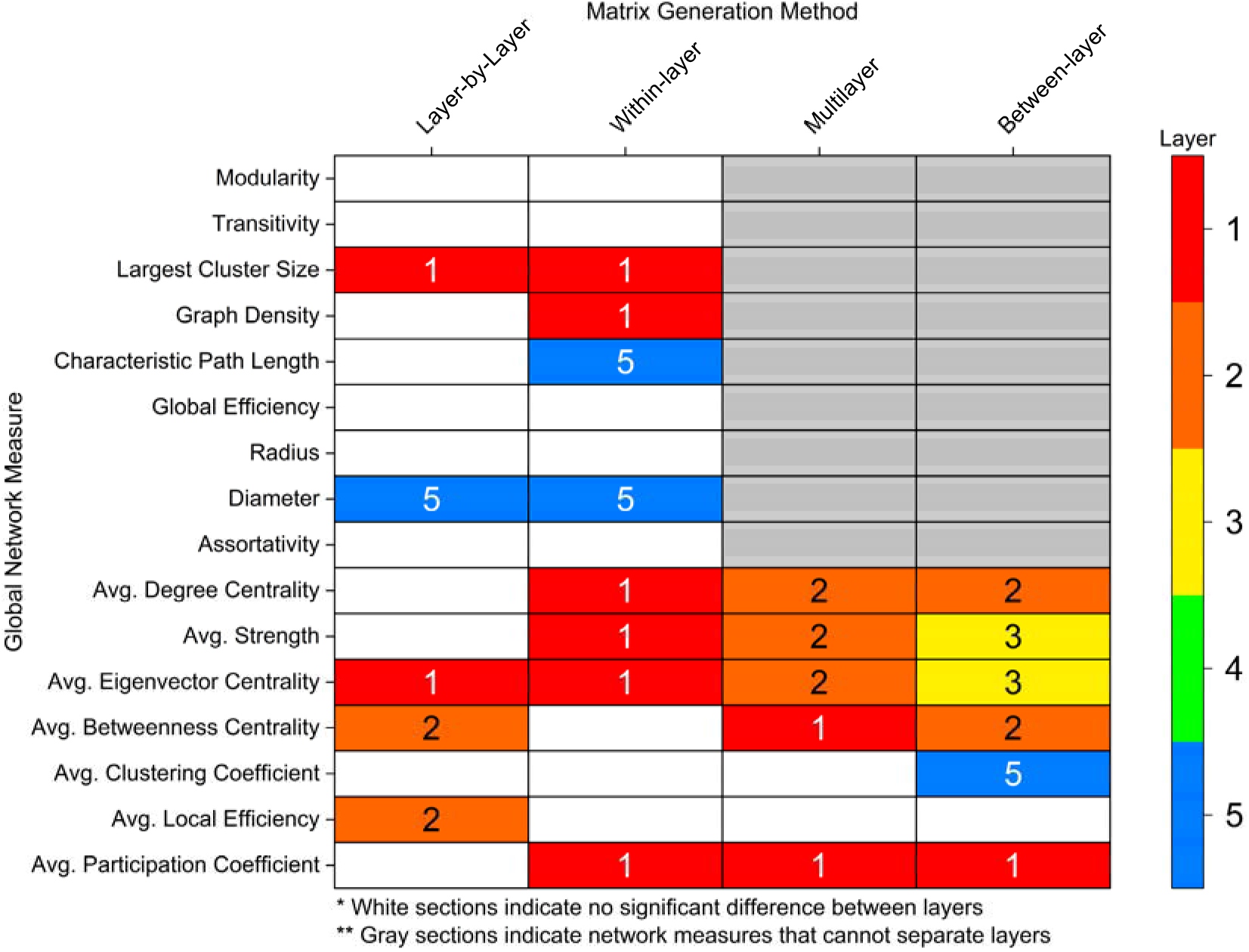
Layer with the highest area-under-the-curve (AUC) value across different global network measures methods (*p* ≤ 0.05). Significance was calculated using a one-way ANOVA with an FDR correction (alpha = 0.05). White sections indicate no significant difference between layers. Gray sections indicate network measures that cannot separate effects of different layers.

For nodal-averaged global measures, which can be applied to all four methods, measures generally decreased from superficial layers (1 and 2) to deeper layers (**Figure 7**). The layer-by-layer methodology identified significant differences in three measures (average eigenvector centrality, average betweenness centrality, and average local efficiency) while the within-layer approach found significant differences in four measures (average degree centrality, average strength, average eigenvector centrality, and average participation coefficient). The multilayer approach also found differences in five measures (average degree centrality, average strength, average eigenvector centrality, average betweenness centrality, and average participation coefficient). Interestingly, the between-layer method identified the most measures with significant differences between layers using six measures, with two measures (average strength and average eigenvector centrality) peaking in the middle layer. Additionally, the between-layer approach was the only method to identify a difference in average clustering coefficient which was the highest in layer 5.

Similar to global measures, the multilayer-based approaches (within-layer, multilayer, between-layer) identified more nodal differences between layers than the layer-by-layer approach (**Table 3–6**). For example, the layer-by-layer method identified a maximum of four nodes with significant differences per measures compared to fifteen for within-layer (degree centrality), fifty-eight for multilayer (betweenness centrality), and fifty-seven for between-layer (betweenness centrality). Despite this, in a majority of measures for all four methods, the limbic region had the greatest number of nodes with significant differences between layers. The right hemisphere also had more significant nodes across all methods and measures (**Table 3–6**). The right (2.4127 ± 0.0306 mm; AVG ± SE) and left (2.4069 ± 0.0303 mm) hemispheres had comparable cortical thicknesses overall and across brain regions (**Figure S29**). Additionally, while the thickness of significant nodes versus non-significant nodes was significantly larger in betweenness centrality metrics (multilayer, between-layer), the absolute difference between significant and non-significant nodes was typically less than 1 mm (**Figure S30**).

Two measures that showed a considerable benefit from the multilayer-based approach were betweenness centrality and clustering coefficient (**Figure 8**). For example, the multilayer and between-layer methods showed substantial increase in the number of nodes that had significant differences between layers. Similarly, the number of nodes with significant differences between layers in clustering coefficient increased using multilayer and between-layer methods. More importantly, however, is clustering coefficient in multilayer and between-layer approaches is the only measure to highlight the deepest layer as having the largest value. Likewise, the multilayer and between-layer methods are the only methods to include nodes that are the highest value in the middle layer (**Figure S21, S26**).

**Figure 8.**
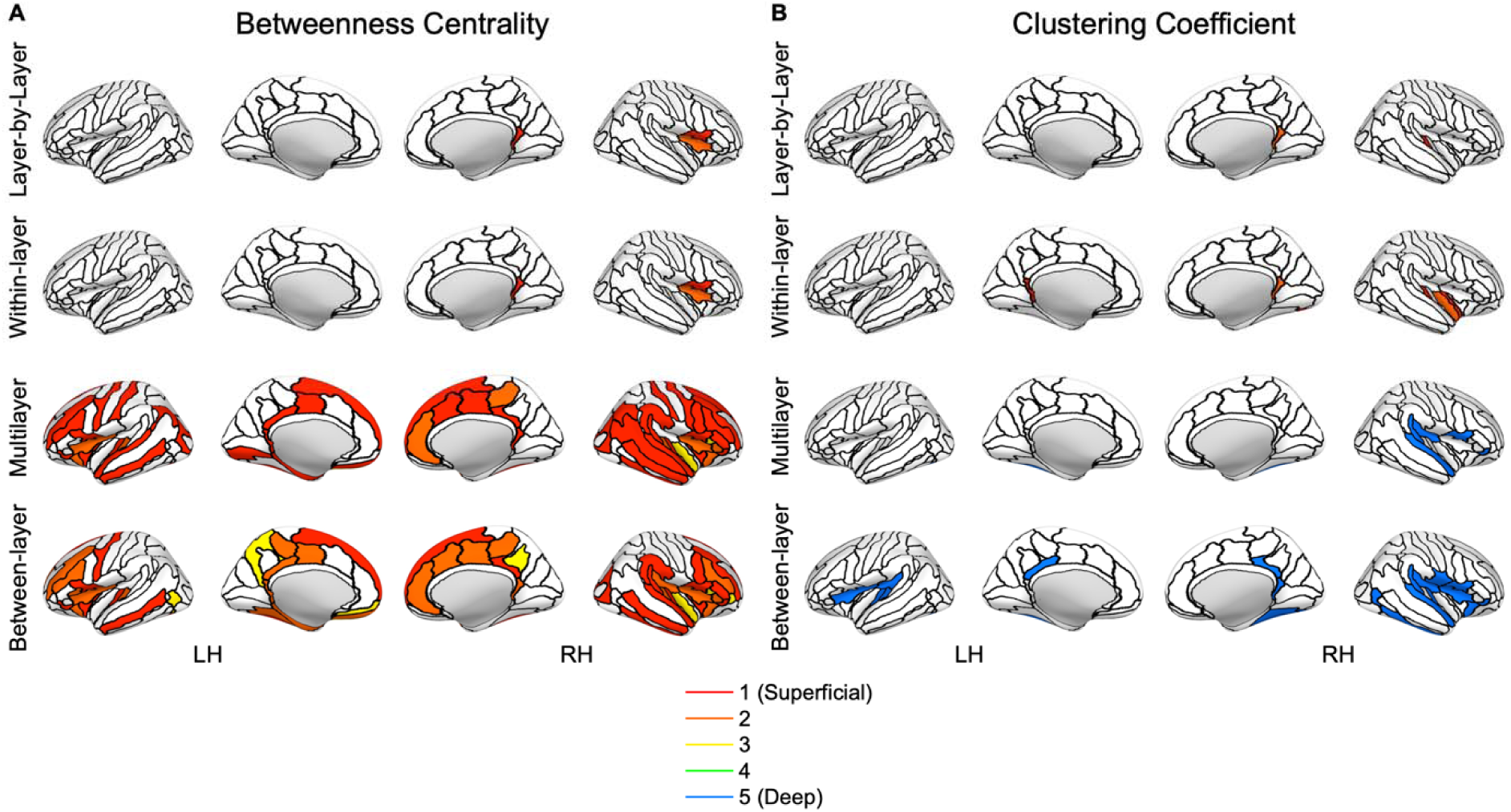
Nodes with significant differences between layers for each nodal measure pipeline (layer-by-layer, within-layer, multilayer, between-layer) for **(A)** betweenness centrality and **(B)** clustering coefficient. Significance was calculated from the area-under-the curve (AUC) value using a one-way ANOVA with an FDR correction (alpha = 0.01) to account for multiple comparisons (Groppe, 2023; Holm, 1979). The colored section represents the layer with the highest value for the node. The nodes are based on the Destrieux atlas in FreeSurfer (Destrieux et al., 2010; Fischl et al., 2004). LH: left hemisphere; RH: right hemisphere.

## Discussion

Graph theory analysis of high-resolution (7T) resting-state fMRI revealed global and nodal network differences between cortical depths. Global integration measures (diameter, characteristic path length) were higher in deeper layers while composition (largest cluster size, graph density, degree centrality, strength) and centrality (eigenvector centrality, betweenness centrality) measures were often higher in superficial layers (**Figure 7**).

Current literature exploring graph theory measures and overall laminar connectivity through networks in the human brain is very limited. Structural analysis of the human connectome using diffusion MRI combined with T_1_-weighted anatomical imaging found qualitative differences in degree, strength, and betweenness centrality nodal distributions across cortical depths (Shamir et al., 2022); however, network-wide global calculations were absent. In a functional approach, Deshpande et al. found no global differences between layers using mean blind deconvoluted Pearson correlations from resting-state fMRI (Deshpande et al., 2022). However, no threshold was used, enabling spurious correlations to impact the mean. Additionally, using the mean across the whole brain obfuscates any impact of a particular ROI. Our results, thus, significantly extend global and nodal network analysis of cortical architecture across the entire brain.

Our findings provide evidence of an advantage of applying of multilayer graph theory to connectomic analysis. While differences between layers were seen across all methodologies, the multilayer approach provided a greater identification of these differences through identifying more measures with larger significant differences (**Figure 7**). Previous connectomic studies have shown a benefit of applying a multilayer network framework (Betzel & Bassett, 2017; De Domenico, 2017; Vaiana & Muldoon, 2018). For example, multilayer connectomics enables the integration of complex neuroimaging data (cortical lamina, frequency bands, multi-modal neuroimaging) (Betzel & Bassett, 2017; Breedt et al., 2023; Buldu & Porter, 2018; Casas-Roma et al., 2022; De Domenico, 2017; Shamir & Assaf, 2023; Vaiana & Muldoon, 2018) and the creation of new network features. New network features can be used to explain neuroscientific findings, as in this work, or even enhance machine learning workflows to better discriminate between disease states (Zhu et al., 2022). Future connectomics studies with available data should therefore consider using a multilayer framework to augment brain network modeling and analysis.

One particular benefit of multilayer analysis in laminar connectomics is the ability to discriminate between and incorporate the impact of within- and between-layer connections. While this comparison was limited to nodal averaged global measures and nodal values, there was still a stark comparison between within- and between-layer connections. When exploring layers individually (within-layer connections), the most superficial layer had the highest activation and is densely connected to itself while the deepest layer was relatively sparsely connected and took longer network paths to transmit information to different brain regions (**Figure 4,7**). However, in between layer connections, layer 1 becomes less important and the superficial-middle layers (layers 2 and 3) become integral for cortical connectivity (**Figure 6–7**). The superficial-middle layers (layers 2 and 3) had the highest between-layer degree, strength, and eigenvector centrality, indicating both layers are densely connected to other layers. Additionally, layer 2’s significantly larger betweenness centrality demonstrates it is the most important layer for information flow between layers (**Figure 6–7**). Lastly, the deepest layer had the highest clustering coefficient meaning it had the highest likelihood of forming local hubs (triangles) with other layers. Thus, within- and between-layer analysis provides evidence of a highly active superficial layer that utilizes layers 2 and 3 to transmit information to other cortical layers.

One important note to contextualize the above findings is that while activity and hubs of information flow can be identified, the direction of information flow cannot be delineated. Cortical layer architecture can have diverse connectivity patterns across layers and hierarchal schemes (Felleman & Van Essen, 1991). Therefore, it is important to recognize the correlation nature of this work rather than infer causality. Furthermore, while our results primarily focus on layers with the highest measure/activity, this does not imply that other layers are inactive. This contextualization is notably important in the context of feedback/feedforward interpretations, a common framework for laminar analysis. In general, feedback is thought to target superficial and deep layers, and feedforward targets the middle layer (Barbas, 2015; Felleman & Van Essen, 1991; Rockland & Pandya, 1979). However, the fact that one area might also send information, makes the interpretation less straightforward. Thus, the information transfer and clustering processes might not directly reflect the feedback/feedforward processes, but complex interactions between them. This intricate feedback/feedforward interplay may also explain the connections between layer 1 and layer 5 (7.06%) being the most consistent between participants (**Figure 2A**) with both superficial and deep layers are activated on opposite sides of the feedback/feedforward circuit (Miyashita, 2022). However, this feedback/feedforward interaction would need to be further validated and studied before drawing concrete conclusions.

The significance of connectivity patterns and characteristics for cortical depths differed across different brain regions. We identified extensive differences between layers in the frontal, limbic, and temporal brain regions (**Figure 8**, **Table 3–6**). Interestingly, the limbic cortex, often with the most significant regions per measures, typically has less layers than other brain regions (Barbas, 2015). Thus, cellular architecture may play a role in the ability for laminar fMRI, and potentially multilayer connectomics, to detect differences between cortical layers. Cortical thickness may also play a role in detecting differences between layers (Fukutomi et al., 2018). The regions (frontal, limbic, and temporal) with the most differences were often the thickest regions (**Figure S29**), with our results overlapping with previous studies (Fukutomi et al., 2018). Additionally, other factors that may impact cortical function and detectability including the neurite density index, orientation dispersion index, and myelin (Fukutomi et al., 2018). However, Fukutomi’s et al. findings show a varied distribution across regions for these measures (Fukutomi et al., 2018). Despite this variation, hot-spots in these metrics near the posterior-ventral part of the cingulate gyrus and transverse temporal sulcus often overlap with significant nodes in our work. Therefore, our results indicate the need to contextualize layer fMRI results within cortical metrics while providing a framework for potential regions (frontal, limbic, temporal) that may be suited for whole-brain laminar analysis.

On a global network level, there were high levels of activation in superficial layers when compared to deeper layers, in line with previous resting-state fMRI analyses reporting increased activation patterns in superficial cortical depths (Guidi et al., 2020; L. Huber et al., 2021; L. R. Huber et al., 2021; Polimeni et al., 2010). This was also reflected in the in higher graph density in superficial layers (**Figure S20**), despite marginally higher tSNR in deeper layers (**Figure S3**). Higher composition and centrality measures indicate a more robustly connected network (**Figure 7**). Additionally, at least within the same network (layer-by-layer and within-layer), deeper layers had significantly longer paths to transmit information, shown by higher global integration measures. It is, however, important to consider the pial vein bias (Polimeni et al., 2010), which increases gradient-echo BOLD signals from the deep to the more superficial parts of cortex. Further studies with alternative contrast mechanisms, which are less affected by the draining vein effect, are thus needed to determine whether the superficial cortical depths play a more critical role in the brain at rest, compared to the deeper aspects of the cortex or if this result is a function of signal strength and variance increasing toward the superficial depth.

### Limitations

This study does have some limitations, both in terms of laminar analysis and connectomic analysis. Regarding our laminar analysis approach, the limitations of this study include biases associated with our fMRI pulse sequence, signal-to-noise ratio bias, the number of cortical depths chosen, the depth to cortical layer incongruence, and the impact of resting-state versus task-based paradigms. This work uses gradient-echo BOLD pulse sequences which may not be fully optimal for laminar analysis (Bandettini et al., 2021; L. Huber et al., 2021). Gradient-echo BOLD can be influenced largely by large draining vessels (Markuerkiaga et al., 2016a; Olman et al., 2007; Polimeni et al., 2010; Turner, 2002), while spin-echo BOLD (Duong et al., 2003; Uludag et al., 2009; Yacoub et al., 2003; Zhao et al., 2004) and VAscular Space Occupancy (VASO) (Chai et al., 2020; Chai et al., 2021) have been proposed as alternative fMRI contrasts for laminar analysis to address this large vein bias (Bandettini et al., 2021; L. Huber et al., 2021). However, VASO and spin-echo BOLD have lower sensitivity and several practical challenges (Moerel et al., 2021). Similarly, signal-to-noise ratio (SNR) can vary at different cortical depths. For example, depths within the middle of the cortex will contain less tissue boundary effects compared to the depths near the pial and white matter surfaces (Blazejewska et al., 2019). This difference may be further exacerbated since the thickness and functionality of cortical layers can change based on brain region (Barbas, 2015; Ding et al., 2009; Zachlod et al., 2020) and cortical curvature (Fatterpekar et al., 2003; Fischl & Dale, 2000; Hilgetag & Barbas, 2006; Van Essen & Maunsell, 1980). However, as shown above for nodal analysis, thickness varied across statistically significant nodes suggesting our results are not purely a function of cortical thickness since significant differences were identified in “thinner” nodes (**Figure S10–S11, S16–S17, S22–S23, S27–S30**). However, for multilayer (*p* < 0.001) and between-layer (*p* < 0.001) betweenness centrality (**Figure S30**), nodes with significant differences between layers had higher cortical thickness. This finding could either result from methodological constraints indicating an inability to detect differences at lower cortical thicknesses or a neurophysiological phenomenon of thicker nodes having larger functional differences between layers. Cortical curvature was not explored, and future laminar work should include the anatomical constraints of the cortex to address this. In addition to the location of the cortical depths chosen, the number of depths can affect the results. Other studies have used a smaller number of depths to ensure independence between depths (Sharoh et al., 2019), six depths to match the number of cortical layers (Pais-Roldan et al., 2023), or even a larger amount that showed an improved detection of cortical responses (Huber et al., 2017). The number of depths chosen should balance independence, cortical response detection, and computational demands from a higher depth count. The number of depths can also impact the role of partial volume effects due to voxel overlap. Furthermore, as mentioned above, the cortical depths do not directly equate to cytoarchitectural cortical layers. Lastly, this study used resting-state fMRI to study whole-brain connectivity; however, laminar resting-state fMRI activation patterns may be different than laminar task-based patterns (Pais-Roldan et al., 2023), limiting the broad applicability to task-based laminar paradigms. Despite potential activation pattern differences, the underlying anatomical basis of resting-state connections (Adachi et al., 2012; Honey et al., 2009; Turk et al., 2016; van den Heuvel et al., 2016) can still inform task-based paradigms. Ideally, a second dataset would be utilized to validate our results; however, few comparable datasets are available.

Regarding our connectomic analysis, limitations include the parcellation choice, network construction approach, thresholding methodology, and multilayer measure calculations. Parcellation choice can impact graph theory results (Albers et al., 2021; Arslan et al., 2018). This work used the Destrieux atlas in FreeSurfer (Destrieux et al., 2010; Fischl et al., 2004), which is based on anatomical nomenclature. However, an atlas derived from functional connectivity (Schaefer et al., 2018) or utilizing functional localizers for specific areas of interest (Nieto-Castanon & Fedorenko, 2012) may be more appropriate for a functional analysis study. Additionally, for laminar analysis, a custom atlas using laminar cytoarchitecture and cortical thickness may improve the accuracy of the results. Another impactful choice in connectomic methodology is how to construct the network from the fMRI time series. Pearson correlations perform better for network construction when using a large number of ROIs (Smith et al., 2011) or in noisier data (Liegeois et al., 2020; Matkovic et al., 2023). However, Pearson correlations also include indirect effects of ROIs which can alter analysis. A popular alternative is partial correlation which utilizes the inverse covariance matrix, and, thus, excludes the indirect network effects. However, partial correlations can also include spurious connections (Berkson’s paradox (Berkson, 1946)) and tend to increase network construction complexity since partial correlations require regularization that has varying optimization parameters (Kim et al., 2015; Pervaiz et al., 2020). Thus, partial correlations may have future utility in laminar multilayer analysis; however, limited ROI number in relation to fMRI time series data points and lack of optimized laminar multilayer regularization parameters led to Pearson correlation being used in this work. Also, as mentioned above, graph theory measures are directly impacted by thresholding the network (Osmanlioglu et al., 2020). AUC analysis attempts to correct for this thresholding bias but still may be inadequate for eliminating thresholding’s effect on network characteristics. Additionally, network measures may be impacted as a result of SNR and network layer normalization (Mandke et al., 2018). For example, increased noise will transition network structure from small-world to more random (Humphries & Gurney, 2008), which may occur as we measure deeper into the cortex. However, our results showed no significant differences in small-worldness between layers, indicating that this network structure change is not occurring in our work (**Figure S7**). Similarly, within multilayer approaches, normalization plays a key role since graph density can influence network properties. However, even with comparable tSNR (**Figure S3**), the graph density varied across cortical depths. While we believe this to be an intrinsic property of cortical connectivity having higher density in superficial cortical depths, as demonstrated by other studies (Logothetis et al., 2001), future work should explore different multilayer normalization schemes in laminar connectivity to more thoroughly parse through this effect (Mandke et al., 2018). Lastly, our statistical analysis of our measures may be limited be the use of ANOVA since it assumes normality and equal variance which are sometimes violated by network measures. Additionally, when selecting peak values for each measure, the highest value was selected instead of using a planned contrast ANOVA.

Increased BOLD signal in superficial vs. deeper layers may be due to vascular-related bias (Markuerkiaga et al., 2016a; Olman et al., 2007; Pais-Roldan et al., 2023; Polimeni et al., 2010; Turner, 2002). One might conclude that the present results reflect vascular biases. The most superficial depth was excluded in this work to reduce this bias; however, the other layers will still have some effect of vascular draining. Additionally, even with removal of the most superficial depth, the current most superficial depth may still be including superficial voxels that are sensitive to vascular-related bias. Despite this limitation, some composition and centrality measures peaked in layers 2 and 3, notably average strength, suggesting that some observed effects are not explainable by biases in superficial layers (**Figure 7, S10–S11, S16–S17, S22– S23, S27–S30**). Lastly, even the utility of the multilayer approach to find more significant effects may be a result of the multilayer model being more sensitive to draining/signal confounds.

## Conclusion

Our multilayer connectomics findings demonstrate global and nodal network differences between cortical depths that can be more aptly identified through the multilayer approach compared to traditional single layer connectomics. These results demonstrate the validity of the multilayer connectomic framework on laminar fMRI and provide a methodological foundation for future multilayer laminar studies. Future work should further explore the intersection of connectomics and laminar studies and address current methodological constraints.

## Supporting Information

## Author Contributions

**Parker Kotlarz:** Conceptualization; Methodology; Software; Validation; Formal analysis; Investigation; Data curation; Writing – original draft; Writing – review & editing; Visualization.

**Kaisu Lankinen:** Conceptualization; Methodology; Software; Investigation; Writing – review & editing; Visualization; Supervision.

**Maria Hakonen:** Methodology; Software; Investigation; Data curation; Writing – review & editing.

**Tori Turpin:** Investigation; Writing – review & editing.

**Jonathan R. Polimeni:** Methodology; Software; Writing – review & editing.

**Jyrki Ahveninen:** Conceptualization; Methodology; Investigation; Writing – review & editing; Supervision; Project administration; Funding acquisition.

## Funding Information

Supported by R01DC017991, R01DC016765, R01DC016915 and P41-EB030006 and was made possible by the resources provided by NIH Shared Instrumentation Grant S10-OD023637.

## Notes

### Competing Interest Statement

The authors have declared no competing interest.

### Summary of Updates

Updated to reflect current draft based on peer-review revisions.

