## Supplemental Data for "Multilayer Network Analysis across Cortical Depths in Resting-State 7T fMRI"

Supplement

| **Table S1.** List of node identifier numbers, region of interest (ROI) short and long names, and their associated brain region. The data in this table is adapted from *Destrieux et al.* (Destrieux et al., 2010; Fischl et al., 2004). | | | | |
| --- | --- | --- | --- | --- |
| L. No. | R. No. | Brain Region | ROI Short Name | ROI Long Name |
| 1 | 75 | Frontal | G_and_S_frontomargin | Fronto-marginal gyrus (of Wernicke) and sulcus |
| 2 | 76 | Occipital | G_and_S_occipital_inf | Inferior occipital gyrus (O3) and sulcus |
| 3 | 77 | Frontal | G_and_S_paracentral | Paracentral lobule and sulcus |
| 4 | 78 | Frontal | G_and_S_subcentral | Subcentral gyrus (central operculum) and sulci |
| 5 | 79 | Frontal | G_and_S_transv_frontopol | Transverse frontopolar gyri and sulci |
| 6 | 80 | Limbic | G_and_S_cingul-Ant | Anterior part of the cingulate gyrus and sulcus (ACC) |
| 7 | 81 | Limbic | G_and_S_cingul-Mid-Ant | Middle-anterior part of the cingulate gyrus and sulcus (aMCC) |
| 8 | 82 | Limbic | G_and_S_cingul-Mid-Post | Middle-posterior part of the cingulate gyrus and sulcus (pMCC) |
| 9 | 83 | Limbic | G_cingul-Post-dorsal | Posterior-dorsal part of the cingulate gyrus (dPCC) |
| 10 | 84 | Limbic | G_cingul-Post-ventral | Posterior-ventral part of the cingulate gyrus (vPCC, isthmus of the cingulate gyrus) |
| 11 | 85 | Occipital | G_cuneus | Cuneus (O6) |
| 12 | 86 | Frontal | G_front_inf-Opercular | Opercular part of the inferior frontal gyrus |
| 13 | 87 | Frontal | G_front_inf-Orbital | Orbital part of the inferior frontal gyrus |
| 14 | 88 | Frontal | G_front_inf-Triangul | Triangular part of the inferior frontal gyrus |
| 15 | 89 | Frontal | G_front_middle | Middle frontal gyrus (F2) |
| 16 | 90 | Frontal | G_front_sup | Superior frontal gyrus (F1) |
| 17 | 91 | Limbic | G_Ins_lg_and_S_cent_ins | Long insular gyrus and central sulcus of the insula |
| 18 | 92 | Limbic | G_insular_short | Short insular gyri |
| 19 | 93 | Occipital | G_occipital_middle | Middle occipital gyrus (O2, lateral occipital gyrus) |
| 20 | 94 | Occipital | G_occipital_sup | Superior occipital gyrus (O1) |
| 21 | 95 | Temporal | G_oc-temp_lat-fusifor | Lateral occipito-temporal gyrus (fusiform gyrus, O4-T4) |
| 22 | 96 | Temporal | G_oc-temp_med-Lingual | Lingual gyrus, ligual part of the medial occipito-temporal gyrus, (O5) |
| 23 | 97 | Temporal | G_oc-temp_med-Parahip | Parahippocampal gyrus, parahippocampal part of the medial occipito-temporal gyrus, (T5) |
| 24 | 98 | Frontal | G_orbital | Orbital gyri |
| 25 | 99 | Parietal | G_pariet_inf-Angular | Angular gyrus |
| 26 | 100 | Parietal | G_pariet_inf-Supramar | Supramarginal gyrus |
| 27 | 101 | Parietal | G_parietal_sup | Superior parietal lobule (lateral part of P1) |
| 28 | 102 | Parietal | G_postcentral | Postcentral gyrus |
| 29 | 103 | Frontal | G_precentral | Precentral gyrus |
| 30 | 104 | Parietal | G_precuneus | Precuneus (medial part of P1) |
| 31 | 105 | Frontal | G_rectus | Straight gyrus, Gyrus rectus |
| 32 | 106 | Limbic | G_subcallosal | Subcallosal area, subcallosal gyrus |
| 33 | 107 | Temporal | G_temp_sup-G_T_transv | Anterior transverse temporal gyrus (of Heschl) |
| 34 | 108 | Temporal | G_temp_sup-Lateral | Lateral aspect of the superior temporal gyrus |
| 35 | 109 | Temporal | G_temp_sup-Plan_polar | Planum polare of the superior temporal gyrus |
| 36 | 110 | Temporal | G_temp_sup-Plan_tempo | Planum temporale or temporal plane of the superior temporal gyrus |
| 37 | 111 | Temporal | G_temporal_inf | Inferior temporal gyrus (T3) |
| 38 | 112 | Temporal | G_temporal_middle | Middle temporal gyrus (T2) |
| 39 | 113 | Frontal | Lat_Fis-ant-Horizont | Horizontal ramus of the anterior segment of the lateral sulcus (or fissure) |
| 40 | 114 | Frontal | Lat_Fis-ant-Vertical | Vertical ramus of the anterior segment of the lateral sulcus (or fissure) |
| 41 | 115 | Limbic | Lat_Fis-post | Posterior ramus (or segment) of the lateral sulcus (or fissure) |
| 42 | 116 | Occipital | Pole_occipital | Occipital pole |
| 43 | 117 | Temporal | Pole_temporal | Temporal pole |
| 44 | 118 | Occipital | S_calcarine | Calcarine sulcus |
| 45 | 119 | Frontal | S_central | Central sulcus (Rolando’s fissure) |
| 46 | 120 | Limbic | S_cingul-Marginalis | Marginal branch (or part) of the cingulate sulcus |
| 47 | 121 | Limbic | S_circular_insula_ant | Anterior segment of the circular sulcus of the insula |
| 48 | 122 | Limbic | S_circular_insula_inf | Inferior segment of the circular sulcus of the insula |
| 49 | 123 | Limbic | S_circular_insula_sup | Superior segment of the circular sulcus of the insula |
| 50 | 124 | Temporal | S_collat_transv_ant | Anterior transverse collateral sulcus |
| 51 | 125 | Temporal | S_collat_transv_post | Posterior transverse collateral sulcus |
| 52 | 126 | Frontal | S_front_inf | Inferior frontal sulcus |
| 53 | 127 | Frontal | S_front_middle | Middle frontal sulcus |
| 54 | 128 | Frontal | S_front_sup | Superior frontal sulcus |
| 55 | 129 | Parietal | S_interm_prim-Jensen | Sulcus intermedius primus (of Jensen) |
| 56 | 130 | Parietal | S_intrapariet_and_P_trans | Intraparietal sulcus (interparietal sulcus) and transverse parietal sulci |
| 57 | 131 | Occipital | S_oc_middle_and_Lunatus | Middle occipital sulcus and lunatus sulcus |
| 58 | 132 | Occipital | S_oc_sup_and_transversal | Superior occipital sulcus and transverse occipital sulcus |
| 59 | 133 | Occipital | S_occipital_ant | Anterior occipital sulcus and preoccipital notch (temporo-occipital incisure) |
| 60 | 134 | Occipital | S_oc-temp_lat | Lateral occipito-temporal sulcus |
| 61 | 135 | Temporal | S_oc-temp_med_and_Lingual | Medial occipito-temporal sulcus (collateral sulcus) and lingual sulcus |
| 62 | 136 | Frontal | S_orbital_lateral | Lateral orbital sulcus |
| 63 | 137 | Frontal | S_orbital_med-olfact | Medial orbital sulcus (olfactory sulcus) |
| 64 | 138 | Frontal | S_orbital-H_Shaped | Orbital sulci (H-shaped sulci) |
| 65 | 139 | Parietal | S_parieto_occipital | Parieto-occipital sulcus (or fissure) |
| 66 | 140 | Limbic | S_pericallosal | Pericallosal sulcus (S of corpus callosum) |
| 67 | 141 | Parietal | S_postcentral | Postcentral sulcus |
| 68 | 142 | Frontal | S_precentral-inf-part | Inferior part of the precentral sulcus |
| 69 | 143 | Frontal | S_precentral-sup-part | Superior part of the precentral sulcus |
| 70 | 144 | Frontal | S_suborbital | Suborbital sulcus (sulcus rostrales, supraorbital sulcus) |
| 71 | 145 | Parietal | S_subparietal | Subparietal sulcus |
| 72 | 146 | Temporal | S_temporal_inf | Inferior temporal sulcus |
| 73 | 147 | Temporal | S_temporal_sup | Superior temporal sulcus (parallel sulcus) |
| 74 | 148 | Temporal | S_temporal_transverse | Transverse temporal sulcus |

| **Table S2.** Area-under-the-curve (AUC) values for each global measure for layer-by-layer analysis. Data are shown as mean ± standard error of mean (standard deviation). One-way ANOVA was used to compare layers with an FDR correction (alpha = 0.05). Graph density and average degree centrality do not have standard error and standard deviation since they are consistent across all participants due to thresholding. Shaded rows have significant differences between layers. P values shown are false discovery rate corrected (FDR, Benjamini-Hochberg method, alpha = 0.05).  * p < 0.05, ** p < 0.01. | | | | | | |
| --- | --- | --- | --- | --- | --- | --- |
| Measures (AUC) | Layer 1 (Superficial) | Layer 2 | Layer 3 | Layer 4 | Layer 5 (Deep) | p value |
| Modularity | 13.33 + 0.52 (2.83) | 13.24 + 0.56 (3.08) | 12.90 + 0.63 (3.43) | 12.27 + 0.68 (3.71) | 11.48 + 0.68 (3.71) | 0.32 |
| Transitivity | 10.81 + 0.32 (1.78) | 11.03 + 0.35 (1.90) | 11.29 + 0.40 (2.17) | 11.47 + 0.44 (2.43) | 11.52 + 0.46 (2.51) | 0.73 |
| Largest Cluster  Size | 5167.97 + 36.53 (200.09) | 5151.87 + 41.83 (229.11) | 5059.57 + 58.99 (323.10) | 4928.27 + 78.80 (431.60) | 4802.27 + 86.16 (471.93) | 0.0024** |
| Graph Density | 7.98 | 7.98 | 7.98 | 7.98 | 7.98 | NaN |
| Characteristic  Path Length | 204.42 + 5.30 (29.06) | 205.75 + 4.96 (27.19) | 207.48 + 4.77 (26.11) | 210.86 + 5.40 (29.58) | 214.96 + 5.85 (32.02) | 0.73 |
| Global Efficiency | 8.97 + 0.21  (1.17) | 8.99 + 0.21  (1.13) | 9.05 + 0.23  (1.23) | 9.09 + 0.29  (1.58) | 9.07 + 0.31  (1.69) | 0.99 |
| Radius | 228.37 + 11.40 (62.41) | 244.91 + 9.13 (50.00) | 249.15 + 9.67 (52.94) | 258.64 + 10.85 (59.42) | 271.26 + 9.97 (54.62) | 0.12 |
| Diameter | 556.97 + 17.06 (93.47) | 568.48 + 14.95 (81.89) | 582.71 + 14.71 (80.57) | 608.22 + 17.49 (95.78) | 629.05 + 18.98 (103.97) | 0.049* |
| Assortativity | 9.19 + 0.47  (2.55) | 9.13 + 0.50  (2.72) | 8.58 + 0.54  (2.96) | 8.11 + 0.56  (3.04) | 7.62 + 0.59  (3.21) | 0.32 |
| Avg. Degree Centrality | 1173.07 | 1173.07 | 1173.07 | 1173.07 | 1173.07 | NaN |
| Avg. Strength | 483.02 + 11.27 (61.70) | 485.82 + 10.73 (58.77) | 484.59 + 10.61 (58.09) | 477.20 + 10.95 (59.98) | 465.23 + 10.05 (55.07) | 0.73 |
| Avg. Eigenvector  Centrality | 2.26 + 0.02  (0.10) | 2.25 + 0.02  (0.13) | 2.22 + 0.03  (0.14) | 2.18 + 0.03  (0.14) | 2.14 + 0.03  (0.15) | 0.028* |
| Avg. Betweenness  Centrality | 6000.07 + 138.76 (760.03) | 6089.39 + 150.75 (825.71) | 5904.46 + 155.92 (853.98) | 5596.98 + 194.75 (1066.66) | 5270.58 + 207.93 (1138.90) | 0.028* |
| Avg. Clustering  Coefficient | 9.68 + 0.22  (1.23) | 9.73 + 0.22  (1.20) | 9.74 + 0.23  (1.23) | 9.56 + 0.23  (1.25) | 9.27 + 0.21  (1.13) | 0.73 |
| Avg. Local  Efficiency | 11.86 + 0.25 (1.39) | 11.88 + 0.24 (1.30) | 11.75 + 0.23 (1.29) | 11.38 + 0.22 (1.23) | 10.91 + 0.19 (1.02) | 0.046* |
| Avg. Participation  Coefficient | 11.82 + 0.47 (2.55) | 11.64 + 0.49 (2.69) | 11.05 + 0.58 (3.17) | 10.46 + 0.67 (3.70) | 10.16 + 0.74 (4.04) | 0.34 |

| **Table S3.** Area-under-the-curve (AUC) values for each significant nodal measure for layer-by-layer analysis. Data are shown as mean ± standard error (standard deviation). One-way ANOVA was used to compare layers. Node long names can be found in Table S1. P values shown are false discovery rate corrected (FDR, Bonferroni-Holm method, alpha = 0.01).  * p < 0.05, ** p < 0.01. | | | | | | |
| --- | --- | --- | --- | --- | --- | --- |
| Node | Layer 1 (Superficial) | Layer 2 | Layer 3 | Layer 4 | Layer 5 (Deep) | P value |
| Degree centrality | | | | | | |
| Left G_cingul-Post-ventral (10) | 982.70 + 80.68 (441.88) | 933.40 + 89.14 (488.26) | 729.17 + 80.08 (438.60) | 534.27 + 74.18 (406.29) | 381.17 + 64.72 (354.47) | < 0.001** |
| Right G_cingul-Post-ventral (84) | 1018.53 + 99.39 (544.36) | 956.00 + 88.54 (484.95) | 717.40 + 84.96 (465.34) | 509.10 + 78.99 (432.63) | 352.03 + 68.67 (376.14) | < 0.001** |
| Right S_circular_insula_sup (123) | 1253.10 + 75.06 (411.11) | 1370.90 + 83.90 (459.53) | 1280.87 + 85.07 (465.93) | 903.67 + 94.45 (517.35) | 683.17 + 93.22 (510.59) | < 0.001** |
| Right S_temporal_transverse (148) | 916.00 + 100.47 (550.31) | 858.67 + 107.08 (586.50) | 625.30 + 88.76 (486.14) | 451.30 + 86.03 (471.18) | 355.03 + 76.07 (416.68) | 0.0046** |
| Strength | | | | | | |
| Left G_cingul-Post-ventral (10) | 360.76 + 32.89 (180.14) | 341.90 + 36.03 (197.36) | 256.32 + 31.64 (173.28) | 174.36 + 26.57 (145.55) | 114.12 + 20.26 (110.96) | < 0.001** |
| Right G_cingul-Post-ventral (84) | 382.55 + 41.50 (227.32) | 352.85 + 37.36 (204.62) | 254.66 + 34.26 (187.63) | 168.14 + 29.52 (161.68) | 107.18 + 23.87 (130.72) | < 0.001** |
| Right S_circular_insula_sup (123) | 490.27 + 34.36 (188.18) | 537.60 + 36.96 (202.46) | 485.49 + 35.65 (195.25) | 325.06 + 39.46 (216.13) | 234.57 + 36.81 (201.64) | < 0.001** |
| Right S_temporal_transverse (148) | 344.05 + 43.08 (235.98) | 320.15 + 44.47 (243.59) | 221.59 + 36.52 (200.02) | 151.01 + 32.12 (175.90) | 114.20 + 26.17 (143.35) | 0.0026** |
| Eigenvector Centrality | | | | | | |
| Left G_cingul-Post-ventral (10) | 1.66 + 0.23 (1.24) | 1.61 + 0.22 (1.20) | 1.19 + 0.18 (0.98) | 0.78 + 0.14 (0.77) | 0.51 + 0.11 (0.60) | 0.0016** |
| Right G_cingul-Post-ventral (84) | 1.79 + 0.23 (1.28) | 1.66 + 0.20 (1.11) | 1.18 + 0.17 (0.96) | 0.76 + 0.14 (0.79) | 0.48 + 0.11 (0.62) | < 0.001** |
| Right S_circular_insula_sup (123) | 2.52 + 0.26 (1.41) | 2.74 + 0.28 (1.55) | 2.40 + 0.25 (1.39) | 1.59 + 0.24 (1.32) | 1.15 + 0.22 (1.20) | 0.0035** |
| Right S_temporal_transverse (148) | 1.72 + 0.27 (1.50) | 1.50 + 0.26 (1.43) | 0.92 + 0.19 (1.06) | 0.57 + 0.16 (0.89) | 0.43 + 0.15 (0.83) | 0.0052** |
| Betweenness Centrality | | | | | | |
| Right G_cingul-Post-ventral (84) | 1842.87 + 492.14 (2695.58) | 1292.67 + 366.50 (2007.43) | 420.93 + 233.78 (1280.47) | 142.27 + 65.30 (357.66) | 60.87 + 48.72 (266.83) | 0.0054** |
| Right G_front_inf-Opercular (86) | 8238.80 + 1148.64 (6291.38) | 5349.93 + 730.46 (4000.89) | 3726.60 + 582.99 (3193.16) | 2879.87 + 514.49 (2817.98) | 2465.00 + 567.53 (3108.50) | < 0.001** |
| Right S_circular_insula_sup (123) | 5156.67 + 879.22 (4815.66) | 6725.47 + 1273.00 (6972.51) | 5025.60 + 796.93 (4364.98) | 1866.33 + 463.74 (2539.99) | 1106.67 + 304.51 (1667.88) | < 0.001** |
| Clustering Coefficient | | | | | | |
| Right G_cingul-Post-ventral (84) | 9.96 + 0.44 (2.41) | 10.02 + 0.45 (2.45) | 9.04 + 0.62 (3.38) | 7.23 + 0.70 (3.82) | 5.53 + 0.70 (3.81) | < 0.001** |
| Right S_temporal_transverse (148) | 11.02 + 0.51 (2.80) | 9.94 + 0.49 (2.71) | 8.63 + 0.61 (3.33) | 6.75 + 0.81 (4.41) | 6.22 + 0.80 (4.39) | < 0.001** |
| Local Efficiency | | | | | | |
| Left G_cingul-Post-ventral (10) | 10.98 + 0.57 (3.14) | 10.60 + 0.63 (3.45) | 9.36 + 0.74 (4.07) | 8.03 + 0.74 (4.07) | 6.19 + 0.79 (4.31) | 0.0013** |
| Right G_cingul-Post-ventral (84) | 11.56 + 0.51 (2.77) | 11.42 + 0.53 (2.88) | 9.95 + 0.71 (3.88) | 7.74 + 0.77 (4.22) | 5.80 + 0.76 (4.16) | < 0.001** |
| Right S_circular_insula_inf (122) | 11.64 + 0.44 (2.41) | 11.79 + 0.51 (2.80) | 10.28 + 0.78 (4.26) | 8.54 + 0.86 (4.73) | 7.47 + 0.86 (4.72) | 0.0043** |
| Right S_temporal_transverse (148) | 12.33 + 0.50 (2.72) | 11.14 + 0.52 (2.82) | 9.47 + 0.65 (3.55) | 7.27 + 0.84 (4.62) | 6.61 + 0.84 (4.62) | < 0.001** |
| Participation Coefficient | | | | | | |
| Left G_cingul-Post-ventral (10) | 13.99 + 1.07 (5.84) | 13.47 + 0.99 (5.44) | 11.33 + 1.13 (6.18) | 8.96 + 1.14 (6.24) | 6.49 + 1.03 (5.65) | < 0.001** |
| Right G_cingul-Post-ventral (84) | 13.21 + 1.22 (6.68) | 13.12 + 1.13 (6.20) | 10.92 + 1.25 (6.83) | 7.99 + 1.27 (6.95) | 5.57 + 1.04 (5.71) | 0.0013** |
| Right S_circular_insula_sup (123) | 13.12 + 0.81 (4.45) | 13.94 + 0.91 (4.98) | 12.17 + 1.06 (5.80) | 8.24 + 1.18 (6.49) | 6.84 + 1.16 (6.37) | < 0.001** |

| **Table S4.** Area-under-the-curve (AUC) values for each global measure for within-layer analysis. Data are shown as mean ± standard error (standard deviation). One-way ANOVA was used to compare layers with an FDR correction (alpha = 0.05). Shaded rows have significant differences between layers. P values shown are false discovery rate corrected (FDR, Benjamini-Hochberg method, alpha = 0.05).  * p < 0.05, ** p < 0.01. | | | | | | |
| --- | --- | --- | --- | --- | --- | --- |
| Measures (AUC) | Layer 1 (Superficial) | Layer 2 | Layer 3 | Layer 4 | Layer 5 (Deep) | p value |
| Modularity | 12.46 + 0.57 (3.12) | 12.57 + 0.59 (3.20) | 12.62 + 0.63 (3.47) | 12.54 + 0.69 (3.79) | 12.53 + 0.72 (3.92) | 0.99 |
| Transitivity | 6.40 + 0.20  (1.12) | 6.48 + 0.23  (1.25) | 6.65 + 0.28  (1.52) | 6.84 + 0.32  (1.76) | 6.93 + 0.35  (1.94) | 0.85 |
| Largest Cluster  Size | 5185.47 + 55.46 (303.78) | 5172.47 + 55.98 (306.64) | 5085.87 + 57.87 (316.96) | 4931.50 + 62.69 (343.35) | 4672.17 + 63.20 (346.18) | < 0.001** |
| Graph Density | 9.33 + 0.30  (1.66) | 9.00 + 0.19  (1.05) | 8.48 + 0.10  (0.55) | 7.83 + 0.19  (1.03) | 6.84 + 0.28  (1.55) | < 0.001** |
| Characteristic  Path Length | 335.95 + 10.47 (57.35) | 341.29 + 9.33 (51.09) | 351.09 + 9.41 (51.55) | 365.12 + 10.38 (56.83) | 383.55 + 10.51 (57.55) | 0.014* |
| Global Efficiency | 5.47 + 0.15  (0.80) | 5.41 + 0.14  (0.77) | 5.36 + 0.16  (0.88) | 5.32 + 0.20  (1.09) | 5.22 + 0.23  (1.23) | 0.99 |
| Radius | 380.51 + 17.46 (95.62) | 418.33 + 15.68 (85.86) | 426.48 + 17.47 (95.68) | 441.42 + 19.34 (105.93) | 458.49 + 19.17 (104.99) | 0.055 |
| Diameter | 909.17 + 29.68 (162.57) | 939.53 + 25.07 (137.33) | 978.45 + 24.02 (131.56) | 1050.08 + 27.60 (151.16) | 1125.11 + 26.34 (144.29) | < 0.001** |
| Assortativity | 8.25 + 0.54  (2.94) | 8.45 + 0.53  (2.90) | 8.37 + 0.55  (3.04) | 8.46 + 0.57  (3.12) | 8.67 + 0.59  (3.21) | 0.99 |
| Avg. Degree Centrality | 1371.76 + 44.52 (243.84) | 1323.73 + 28.13 (154.07) | 1246.24 + 14.78 (80.96) | 1151.24 + 27.63 (151.35) | 1005.13 + 41.71 (228.45) | < 0.001** |
| Avg. Strength | 325.74 + 13.93 (76.29) | 315.00 + 10.40 (56.94) | 299.71 + 10.04 (54.97) | 280.68 + 13.20 (72.31) | 246.04 + 15.64 (85.67) | < 0.001** |
| Avg. Eigenvector  Centrality | 2.32 + 0.03  (0.17) | 2.30 + 0.03  (0.17) | 2.25 + 0.03  (0.16) | 2.16 + 0.02  (0.13) | 2.05 + 0.02  (0.12) | < 0.001** |
| Avg. Betweenness  Centrality | 5600.25 + 190.38 (1042.78) | 5704.47 + 160.09 (876.86) | 5747.65 + 155.90 (853.90) | 5727.50 + 191.92 (1051.21) | 5540.19 + 194.97 (1067.90) | 0.99 |
| Avg. Clustering  Coefficient | 5.77 + 0.15  (0.80) | 5.75 + 0.14  (0.77) | 5.74 + 0.16  (0.90) | 5.66 + 0.19  (1.06) | 5.38 + 0.22  (1.20) | 0.72 |
| Avg. Local  Efficiency | 7.01 + 0.16  (0.89) | 6.98 + 0.15  (0.84) | 6.90 + 0.17  (0.91) | 6.72 + 0.19  (1.01) | 6.32 + 0.20  (1.10) | 0.055 |
| Avg. Participation  Coefficient | 12.69 + 0.54 (2.96) | 12.19 + 0.56 (3.06) | 11.32 + 0.60 (3.31) | 10.31 + 0.64 (3.48) | 9.31 + 0.63  (3.43) | 0.0011** |

| **Table S5.** Area-under-the-curve (AUC) values for each significant nodal measure for within-layer analysis. Data are shown as mean ± standard error (standard deviation). One-way ANOVA was used to compare layers. Node long names can be found in Table S1. P values shown are false discovery rate corrected (FDR, Bonferroni-Holm method, alpha = 0.01).  * p < 0.05, ** p < 0.01. | | | | | | |
| --- | --- | --- | --- | --- | --- | --- |
| Node | Layer 1 (Superficial) | Layer 2 | Layer 3 | Layer 4 | Layer 5 (Deep) | P value |
| Degree centrality | | | | | | |
| Left G_and_S_cingul-Mid-Post (8) | 2453.70 + 102.00 (558.67) | 2369.50 + 83.28 (456.16) | 2181.00 + 78.71 (431.10) | 1948.07 + 92.23 (505.15) | 1658.67 + 103.09 (564.65) | < 0.001** |
| G_cingul-Post-ventral (10) | 1235.83 + 105.41 (577.37) | 1119.87 + 102.18 (559.66) | 802.23 + 83.68 (458.31) | 505.50 + 69.92 (382.98) | 281.77 + 53.47 (292.89) | < 0.001** |
| Left Lat_Fis-post (41) | 1940.63 + 129.80 (710.95) | 1876.60 + 121.41 (665.00) | 1655.17 + 125.67 (688.31) | 1368.37 + 137.19 (751.44) | 1073.03 + 131.74 (721.57) | 0.0012** |
| Left S_circular_insula_sup (49) | 1753.07 + 113.61 (622.26) | 1805.33 + 103.36 (566.13) | 1701.57 + 90.66 (496.56) | 1408.00 + 97.00 (531.31) | 1014.70 + 105.06 (575.45) | < 0.001** |
| Right G_and_S_subcentral (78) | 1617.93 + 104.91 (574.59) | 1464.47 + 102.18 (559.67) | 1301.80 + 98.82 (541.24) | 1053.93 + 103.29 (565.77) | 820.33 + 118.63 (649.79) | < 0.001** |
| Right G_and_S_cingul-Mid-Post (82) | 2416.90 + 102.21 (559.83) | 2355.47 + 94.21 (516.03) | 2198.57 + 93.25 (510.73) | 2015.23 + 100.85 (552.35) | 1734.57 + 104.51 (572.42) | 0.0014** |
| Right G_cingul-Post-ventral (84) | 1289.37 + 134.01 (734.02) | 1163.60 + 111.91 (612.97) | 795.47 + 94.86 (519.57) | 472.33 + 73.18 (400.84) | 223.30 + 44.61 (244.34) | < 0.001** |
| Right G_oc-temp_lat-fusifor (95) | 1703.73 + 137.81 (754.82) | 1564.90 + 128.28 (702.64) | 1361.43 + 109.30 (598.69) | 1085.97 + 101.49 (555.90) | 783.80 + 109.15 (597.82) | < 0.001** |
| Right G_temp_sup-G_T_transv (107) | 1137.67 + 110.64 (605.98) | 1008.40 + 111.03 (608.14) | 827.27 + 93.86 (514.10) | 637.50 + 87.57 (479.64) | 448.97 + 88.05 (482.26) | 0.0012** |
| Right G_temp_sup-Lateral (108) | 1631.83 + 108.81 (595.98) | 1409.97 + 96.23 (527.08) | 1180.43 + 98.23 (538.00) | 1022.60 + 109.24 (598.31) | 866.70 + 115.06 (630.22) | < 0.001** |
| Right G_temp_sup-Plan_tempo (110) | 1525.93 + 123.89 (678.59) | 1423.33 + 120.28 (658.81) | 1231.83 + 108.30 (593.17) | 949.63 + 101.54 (556.14) | 722.20 + 109.61 (600.35) | < 0.001** |
| Right Lat_Fis-post (115) | 2042.73 + 143.53 (786.16) | 2004.73 + 134.18 (734.91) | 1783.97 + 116.03 (635.50) | 1426.77 + 112.17 (614.40) | 996.20 + 108.13 (592.24) | < 0.001** |
| Right S_circular_insula_inf (122) | 1259.37 + 130.51 (714.81) | 1198.43 + 120.04 (657.48) | 931.47 + 105.74 (579.14) | 646.30 + 102.02 (558.78) | 505.97 + 102.45 (561.13) | < 0.001** |
| Right S_circular_insula_sup (123) | 1481.80 + 109.33 (598.84) | 1550.13 + 104.59 (572.85) | 1360.07 + 88.90 (486.94) | 878.80 + 100.41 (549.96) | 562.57 + 101.58 (556.38) | < 0.001** |
| Right S_temporal_transverse (148) | 1143.17 + 130.21 (713.17) | 1019.17 + 127.48 (698.26) | 672.10 + 90.28 (494.50) | 425.37 + 82.21 (450.30) | 265.77 + 74.80 (409.69) | < 0.001** |
| Strength | | | | | | |
| G_cingul-Post-ventral (10) | 264.18 + 25.45 (139.42) | 236.06 + 24.36 (133.40) | 162.14 + 19.10 (104.60) | 95.76 + 14.31 (78.40) | 48.61 + 9.10 (49.86) | < 0.001** |
| Left S_circular_insula_sup (49) | 418.19 + 32.81 (179.69) | 429.25 + 31.70 (173.63) | 398.91 + 30.56 (167.39) | 325.88 + 31.59 (173.00) | 229.18 + 29.67 (162.50) | 0.0040** |
| Right G_cingul-Post-ventral (84) | 283.14 + 32.94 (180.44) | 248.07 + 27.59 (151.14) | 163.77 + 23.03 (126.16) | 92.58 + 16.54 (90.61) | 40.48 + 9.01 (49.33) | < 0.001** |
| Right G_oc-temp_lat-fusifor (95) | 421.04 + 39.58 (216.80) | 382.91 + 36.23 (198.45) | 328.21 + 31.17 (170.73) | 257.55 + 29.28 (160.38) | 184.46 + 30.90 (169.24) | 0.0010** |
| Right G_temp_sup-G_T_transv (107) | 250.02 + 28.13 (154.05) | 217.63 + 27.03 (148.04) | 172.72 + 22.65 (124.04) | 130.90 + 20.46 (112.07) | 91.83 + 20.57 (112.68) | 0.0044** |
| Right G_temp_sup-Plan_tempo (110) | 361.66 + 33.53 (183.64) | 333.42 + 31.29 (171.36) | 285.03 + 28.22 (154.56) | 220.65 + 28.59 (156.59) | 170.30 + 31.81 (174.22) | 0.0093** |
| Right Lat_Fis-post (115) | 511.70 + 42.28 (231.60) | 497.40 + 39.74 (217.66) | 429.71 + 32.93 (180.37) | 330.49 + 29.25 (160.23) | 220.23 + 26.59 (145.65) | < 0.001** |
| Right S_circular_insula_sup (123) | 340.46 + 30.37 (166.34) | 353.82 + 27.97 (153.18) | 299.25 + 22.66 (124.13) | 186.79 + 25.47 (139.50) | 116.80 + 25.01 (137.01) | < 0.001** |
| Right S_temporal_transverse (148) | 247.39 + 31.99 (175.20) | 217.66 + 30.16 (165.19) | 135.79 + 20.74 (113.61) | 81.99 + 17.37 (95.16) | 49.13 + 13.78 (75.47) | < 0.001** |
| Eigenvector centrality | | | | | | |
| G_cingul-Post-ventral (10) | 1.84 + 0.23 (1.26) | 1.73 + 0.22 (1.19) | 1.24 + 0.18 (0.97) | 0.76 + 0.14 (0.75) | 0.41 + 0.10 (0.52) | < 0.001** |
| Right G_cingul-Post-ventral (84) | 1.96 + 0.24 (1.33) | 1.79 + 0.21 (1.14) | 1.24 + 0.18 (0.99) | 0.73 + 0.14 (0.75) | 0.35 + 0.09 (0.49) | < 0.001** |
| Right S_circular_insula_sup (123) | 2.57 + 0.25 (1.39) | 2.78 + 0.28 (1.51) | 2.43 + 0.25 (1.38) | 1.57 + 0.25 (1.35) | 1.03 + 0.23 (1.24) | < 0.001** |
| Right S_temporal_transverse (148) | 1.84 + 0.27 (1.49) | 1.60 + 0.26 (1.45) | 0.95 + 0.19 (1.06) | 0.53 + 0.16 (0.86) | 0.33 + 0.15 (0.80) | < 0.001** |
| Betweenness centrality | | | | | | |
| Right G_cingul-Post-ventral (84) | 1809.80 + 482.95 (2645.25) | 1206.27 + 343.75 (1882.77) | 410.07 + 222.18 (1216.94) | 138.33 + 64.88 (355.38) | 12.67 + 10.27 (56.26) | 0.0033** |
| Right G_front_inf-Opercular (86) | 7844.80 + 1089.17 (5965.60) | 5032.47 + 689.14 (3774.59) | 3672.47 + 574.83 (3148.48) | 2797.33 + 513.13 (2810.54) | 2104.87 + 581.72 (3186.20) | < 0.001** |
| Right S_circular_insula_sup (123) | 4576.87 + 797.64 (4368.84) | 6260.67 + 1189.89 (6517.27) | 4952.67 + 849.55 (4653.15) | 1833.93 + 452.36 (2477.70) | 1200.27 + 317.18 (1737.29) | 0.0022** |
| Clustering coefficient | | | | | | |
| G_cingul-Post-ventral (10) | 5.54 + 0.30 (1.67) | 5.35 + 0.32 (1.78) | 4.85 + 0.37 (2.04) | 4.32 + 0.38 (2.10) | 3.00 + 0.43 (2.37) | 0.0017** |
| Right G_cingul-Post-ventral (84) | 5.83 + 0.31 (1.67) | 5.90 + 0.29 (1.61) | 5.34 + 0.34 (1.88) | 4.22 + 0.38 (2.08) | 2.68 + 0.39 (2.13) | < 0.001** |
| Right G_Ins_lg_and_S_cent_ins (91) | 6.45 + 0.22 (1.20) | 6.17 + 0.32 (1.74) | 6.10 + 0.31 (1.72) | 5.40 + 0.34 (1.85) | 4.27 + 0.46 (2.51) | 0.0078** |
| Right S_circular_insula_inf (122) | 6.01 + 0.22 (1.21) | 6.04 + 0.24 (1.33) | 5.42 + 0.39 (2.16) | 4.39 + 0.50 (2.72) | 3.61 + 0.56 (3.07) | 0.0063** |
| Right S_collat_transv_post (125) | 7.74 + 0.36 (1.98) | 7.53 + 0.36 (2.00) | 7.17 + 0.36 (1.97) | 6.43 + 0.41 (2.26) | 5.20 + 0.49 (2.67) | 0.0092** |
| Right S_temporal_transverse (148) | 6.39 + 0.34 (1.88) | 5.95 + 0.32 (1.78) | 5.16 + 0.35 (1.91) | 3.85 + 0.47 (2.55) | 3.06 + 0.49 (2.67) | < 0.001** |
| Local efficiency | | | | | | |
| G_cingul-Post-ventral (10) | 6.56 + 0.34 (1.87) | 6.26 + 0.36 (1.98) | 5.49 + 0.42 (2.29) | 4.69 + 0.43 (2.35) | 3.16 + 0.46 (2.51) | < 0.001** |
| Right G_cingul-Post-ventral (84) | 6.74 + 0.34 (1.88) | 6.70 + 0.34 (1.85) | 5.87 + 0.40 (2.17) | 4.51 + 0.43 (2.34) | 2.81 + 0.42 (2.32) | < 0.001** |
| Right G_Ins_lg_and_S_cent_ins (91) | 7.11 + 0.25 (1.40) | 6.85 + 0.35 (1.89) | 6.70 + 0.33 (1.83) | 5.91 + 0.37 (2.00) | 4.67 + 0.48 (2.64) | 0.0029** |
| Right G_temp_sup-G_T_transv (107) | 7.18 + 0.31 (1.71) | 6.62 + 0.34 (1.85) | 5.79 + 0.42 (2.29) | 5.15 + 0.48 (2.62) | 4.19 + 0.55 (3.00) | 0.0019** |
| Right S_circular_insula_inf (122) | 6.88 + 0.26 (1.41) | 6.89 + 0.28 (1.55) | 6.14 + 0.43 (2.33) | 4.97 + 0.52 (2.86) | 4.03 + 0.59 (3.24) | < 0.001** |
| Right S_circular_insula_sup (123) | 7.24 + 0.28 (1.51) | 7.09 + 0.27 (1.49) | 6.90 + 0.29 (1.61) | 6.15 + 0.36 (1.97) | 4.73 + 0.52 (2.86) | < 0.001** |
| Right S_collat_transv_post (125) | 8.53 + 0.36 (1.95) | 8.33 + 0.36 (1.99) | 7.97 + 0.37 (2.05) | 7.16 + 0.44 (2.43) | 5.86 + 0.53 (2.91) | 0.0068** |
| Right S_temporal_transverse (148) | 7.13 + 0.34 (1.85) | 6.64 + 0.33 (1.83) | 5.64 + 0.36 (1.98) | 4.15 + 0.48 (2.65) | 3.24 + 0.51 (2.77) | < 0.001** |
| Participation coefficient | | | | | | |
| G_cingul-Post-ventral (10) | 14.89 + 1.12 (6.15) | 13.98 + 1.04 (5.71) | 11.66 + 1.12 (6.15) | 8.95 + 1.10 (6.00) | 4.91 + 0.93 (5.10) | < 0.001** |
| Left G_oc-temp_med-Parahip (23) | 13.20 + 1.09 (5.99) | 12.26 + 1.14 (6.26) | 9.73 + 1.29 (7.06) | 6.82 + 1.23 (6.75) | 3.98 + 1.09 (6.00) | < 0.001** |
| Right G_cingul-Post-ventral (84) | 14.06 + 1.30 (7.10) | 13.67 + 1.16 (6.34) | 11.11 + 1.27 (6.95) | 7.85 + 1.25 (6.84) | 4.21 + 0.92 (5.04) | < 0.001** |
| Right G_oc-temp_med-Parahip (97) | 9.93 + 1.19 (6.52) | 9.37 + 1.23 (6.73) | 6.70 + 1.17 (6.41) | 4.54 + 1.16 (6.35) | 2.77 + 0.90 (4.95) | 0.0032** |
| Right G_temp_sup-G_T_transv (107) | 11.29 + 1.01 (5.55) | 10.65 + 1.10 (6.00) | 9.43 + 1.18 (6.45) | 6.59 + 1.08 (5.93) | 3.56 + 0.89 (4.86) | < 0.001** |
| Right Lat_Fis-ant-Vertical (114) | 8.14 + 1.04 (5.67) | 8.62 + 1.03 (5.64) | 6.72 + 1.03 (5.67) | 3.72 + 0.75 (4.09) | 1.79 + 0.63 (3.47) | < 0.001** |
| Right S_circular_insula_sup (123) | 14.17 + 0.80 (4.39) | 14.52 + 0.92 (5.06) | 12.45 + 1.07 (5.84) | 8.30 + 1.24 (6.79) | 5.94 + 1.09 (5.97) | < 0.001** |
| Right S_oc-temp_lat (134) | 11.08 + 0.98 (5.38) | 9.72 + 1.12 (6.15) | 7.70 + 1.12 (6.11) | 5.45 + 0.91 (4.99) | 4.87 + 0.82 (4.51) | 0.0038** |
| Right S_temporal_transverse (148) | 10.84 + 1.21 (6.64) | 10.67 + 1.18 (6.48) | 8.30 + 1.14 (6.24) | 5.03 + 0.98 (5.35) | 3.31 + 0.80 (4.36) | < 0.001** |

| **Table S6.** Area-under-the-curve (AUC) values for each global measure for the multilayer analysis. NaN values are due to global s that cannot be derived for specific layers. Data are shown as mean ± standard error (standard deviation). One-way ANOVA was used to compare layers with an FDR correction (alpha = 0.05). Graph density and average degree centrality do not have standard error and standard deviation since they are consistent across all participants due to thresholding. Shaded rows have significant differences between layers. P values shown are false discovery rate corrected (FDR, Benjamini-Hochberg method, alpha = 0.05).  * p < 0.05, ** p < 0.01. | | | | | | | |
| --- | --- | --- | --- | --- | --- | --- | --- |
| Measures (AUC) | Layer 1 (Superficial) | Layer 2 | Layer 3 | Layer 4 | Layer 5 (Deep) | Multilayer | p value |
| Modularity | NaN | NaN | NaN | NaN | NaN | 13.44 + 0.58 (3.20) | NaN |
| Transitivity | NaN | NaN | NaN | NaN | NaN | 6.88 + 0.26 (1.42) | NaN |
| Largest Cluster  Size | NaN | NaN | NaN | NaN | NaN | 26720.50 + 198.88 (1089.31) | NaN |
| Graph Density | NaN | NaN | NaN | NaN | NaN | 7.98 | NaN |
| Characteristic  Path Length | NaN | NaN | NaN | NaN | NaN | 356.98 + 8.08 (44.27) | NaN |
| Global Efficiency | NaN | NaN | NaN | NaN | NaN | 5.20 + 0.12 (0.68) | NaN |
| Radius | NaN | NaN | NaN | NaN | NaN | 282.04 + 20.57 (112.65) | NaN |
| Diameter | NaN | NaN | NaN | NaN | NaN | 1276.96 + 32.24 (176.58) | NaN |
| Assortativity | NaN | NaN | NaN | NaN | NaN | 10.03 + 0.50 (2.74) | NaN |
| Avg. Degree Centrality | 6120.06 + 109.88 (601.86) | 6227.01 + 66.25 (362.87) | 6117.00 + 29.75 (162.96) | 5778.98 + 63.54 (348.03) | 5243.04 + 114.25 (625.76) | 5897.22 | < 0.001** |
| Avg. Strength | 1471.07 + 41.23 (225.80) | 1519.12 + 38.35 (210.05) | 1506.81 + 41.33 (226.37) | 1425.36 + 46.45 (254.42) | 1279.02 + 52.03 (285.00) | 1440.27 + 36.56 (200.23) | 0.0019** |
| Avg. Eigenvector  Centrality | 0.99 + 0.03 (0.18) | 1.03 + 0.02 (0.13) | 1.03 + 0.02 (0.08) | 0.99 + 0.01 (0.08) | 0.89 + 0.02 (0.13) | 0.99 + 0.01 (0.06) | < 0.001** |
| Avg. Betweenness  Centrality | 61782.76 + 3026.44 (16576.48) | 41292.92 + 1015.60 (5562.69) | 31429.48 + 1187.24 (6502.81) | 23221.57 + 1255.80 (6878.27) | 16319.54 + 1597.85 (8751.78) | 34809.25 + 634.48 (3475.21) | < 0.001** |
| Avg. Clustering  Coefficient | 6.92 + 0.22 (1.19) | 6.94 + 0.21 (1.14) | 7.05 + 0.20 (1.11) | 7.26 + 0.20 (1.11) | 7.67 + 0.21 (1.16) | 7.17 + 0.21 (1.13) | 0.080 |
| Avg. Local  Efficiency | 8.39 + 0.21 (1.14) | 8.47 + 0.21 (1.14) | 8.53 + 0.21 (1.13) | 8.58 + 0.21 (1.13) | 8.74 + 0.21 (1.15) | 8.54 + 0.21 (1.13) | 0.81 |
| Avg. Participation  Coefficient | 13.12 + 0.61 (3.33) | 12.83 + 0.61 (3.35) | 12.25 + 0.61 (3.34) | 11.45 + 0.60 (3.28) | 10.45 + 0.57 (3.13) | 12.02 + 0.59 (3.22) | 0.019* |

| **Table S7.** Area-under-the-curve (AUC) values for each significant nodal measure for multilayer analysis. Data are shown as mean ± standard error (standard deviation). One-way ANOVA was used to compare layers. Node long names can be found in Table S1. P values shown are false discovery rate corrected (FDR, Bonferroni-Holm method, alpha = 0.01).  * p < 0.05, ** p < 0.01. | | | | | | |
| --- | --- | --- | --- | --- | --- | --- |
| Node | Layer 1 (Superficial) | Layer 2 | Layer 3 | Layer 4 | Layer 5 (Deep) | P value |
| Degree centrality | | | | | | |
| Left G_cingul-Post-ventral (10) | 5076.87 + 431.15 (2361.53) | 4969.63 + 452.10 (2476.26) | 3985.50 + 405.98 (2223.62) | 2812.13 + 373.43 (2045.38) | 1630.10 + 287.36 (1573.94) | < 0.001** |
| Left S_circular_insula_sup (49) | 7193.63 + 486.46 (2664.46) | 8094.03 + 482.03 (2640.19) | 8278.37 + 439.28 (2406.03) | 7029.77 + 462.79 (2534.79) | 5169.37 + 485.06 (2656.79) | 0.0062** |
| Right G_cingul-Post-ventral (84) | 5147.20 + 540.39 (2959.82) | 5051.87 + 480.47 (2631.63) | 3938.47 + 445.94 (2442.51) | 2751.13 + 394.04 (2158.26) | 1539.73 + 280.11 (1534.20) | < 0.001** |
| Right G_temp_sup-G_T_transv (107) | 5177.73 + 519.32 (2844.45) | 4754.93 + 518.62 (2840.60) | 4123.13 + 457.67 (2506.78) | 3238.60 + 420.49 (2303.14) | 2119.83 + 366.05 (2004.92) | 0.0043** |
| Right S_circular_insula_sup (123) | 5919.10 + 492.79 (2699.12) | 6862.40 + 475.47 (2604.27) | 6705.53 + 425.99 (2333.27) | 4324.00 + 461.73 (2528.98) | 2521.00 + 415.35 (2274.98) | < 0.001** |
| Right S_temporal_transverse (148) | 4721.87 + 572.15 (3133.78) | 4574.57 + 570.52 (3124.88) | 3376.47 + 433.26 (2373.07) | 2286.10 + 407.43 (2231.56) | 1349.23 + 317.51 (1739.06) | < 0.001** |
| Strength | | | | | | |
| Left G_cingul-Post-ventral (10) | 1076.90 + 99.01 (542.29) | 1057.98 + 106.84 (585.16) | 826.63 + 93.02 (509.48) | 557.68 + 78.19 (428.29) | 298.81 + 52.85 (289.47) | < 0.001** |
| Right G_cingul-Post-ventral (84) | 1122.08 + 128.86 (705.81) | 1087.16 + 117.50 (643.58) | 830.00 + 107.62 (589.48) | 564.44 + 90.62 (496.36) | 302.89 + 62.34 (341.47) | < 0.001** |
| Right G_temp_sup-G_T_transv (107) | 1137.14 + 124.93 (684.29) | 1043.00 + 123.51 (676.49) | 889.30 + 111.09 (608.45) | 684.54 + 96.70 (529.64) | 431.61 + 81.54 (446.62) | 0.0068** |
| Right S_circular_insula_sup (123) | 1322.47 + 126.86 (694.86) | 1550.80 + 120.67 (660.95) | 1498.64 + 105.59 (578.36) | 938.46 + 115.63 (633.34) | 529.45 + 101.39 (555.31) | < 0.001** |
| Right S_temporal_transverse (148) | 1003.81 + 132.47 (725.58) | 981.60 + 131.61 (720.83) | 704.36 + 99.97 (547.55) | 458.87 + 84.51 (462.87) | 250.96 + 55.15 (302.06) | < 0.001** |
| Eigenvector centrality | | | | | | |
| Left G_cingul-Post-ventral (10) | 0.72 + 0.09 (0.47) | 0.71 + 0.09 (0.48) | 0.55 + 0.08 (0.42) | 0.38 + 0.07 (0.36) | 0.21 + 0.05 (0.25) | < 0.001** |
| Right G_cingul-Post-ventral (84) | 0.79 + 0.11 (0.59) | 0.75 + 0.10 (0.52) | 0.56 + 0.08 (0.45) | 0.38 + 0.07 (0.37) | 0.20 + 0.04 (0.24) | < 0.001** |
| Right S_circular_insula_sup (123) | 0.95 + 0.12 (0.66) | 1.13 + 0.12 (0.68) | 1.09 + 0.11 (0.62) | 0.67 + 0.10 (0.56) | 0.37 + 0.08 (0.45) | < 0.001** |
| Betweenness centrality | | | | | | |
| Left G_and_S_cingul-Mid-Post (8) | 105111.27 + 21687.37 (118786.64) | 88184.53 + 13701.49 (75046.13) | 53839.20 + 10915.52 (59786.76) | 25821.60 + 6124.16 (33543.42) | 11866.60 + 3565.61 (19529.64) | < 0.001** |
| Left G_cingul-Post-dorsal (9) | 52676.60 + 12036.96 (65929.12) | 33667.53 + 9945.04 (54471.23) | 8431.53 + 2396.37 (13125.47) | 6178.80 + 1168.20 (6398.47) | 4070.73 + 3076.98 (16853.32) | < 0.001** |
| Left G_cingul-Post-ventral (10) | 41191.27 + 8343.09 (45696.99) | 34999.13 + 7045.00 (38587.07) | 19735.00 + 3091.28 (16931.64) | 16520.60 + 2274.77 (12459.45) | 118.60 + 99.52 (545.10) | < 0.001** |
| Left G_front_inf-Opercular (12) | 92545.73 + 13077.13 (71626.40) | 33567.13 + 7110.37 (38945.10) | 19672.27 + 4132.23 (22633.16) | 11064.33 + 3243.25 (17763.99) | 3848.73 + 1381.32 (7565.78) | < 0.001** |
| Left G_front_inf-Orbital (13) | 49827.53 + 8908.29 (48792.71) | 16566.47 + 3174.81 (17389.12) | 9858.60 + 1910.07 (10461.87) | 9051.00 + 1659.84 (9091.32) | 4139.80 + 2511.86 (13758.02) | < 0.001** |
| Left G_front_middle (15) | 197365.53 + 31853.65 (174469.61) | 95720.40 + 13721.65 (75156.58) | 75488.80 + 11051.19 (60529.87) | 66872.73 + 11962.40 (65520.76) | 38275.20 + 6217.33 (34053.73) | < 0.001** |
| Left G_front_sup (16) | 313885.67 + 40031.09 (219259.30) | 142010.27 + 18217.86 (99783.30) | 80349.13 + 12807.12 (70147.47) | 65373.80 + 8248.62 (45179.57) | 88886.93 + 15911.67 (87151.82) | < 0.001** |
| Left G_insular_short (18) | 31900.33 + 6397.00 (35037.80) | 17459.27 + 3608.71 (19765.71) | 9014.27 + 1829.48 (10020.46) | 8794.20 + 1569.99 (8599.19) | 2107.13 + 1122.81 (6149.87) | < 0.001** |
| Left G_occipital_middle (19) | 237544.73 + 48014.28 (262985.06) | 140370.93 + 33163.87 (181645.97) | 104380.53 + 16827.03 (92165.41) | 82597.40 + 15025.27 (82296.81) | 41015.93 + 11499.84 (62987.20) | 0.0043** |
| Left G_oc-temp_lat-fusifor (21) | 96157.40 + 19979.53 (109432.37) | 39225.13 + 15223.08 (83380.22) | 20072.27 + 4128.17 (22610.92) | 14707.20 + 4454.70 (24399.39) | 4135.80 + 2410.01 (13200.16) | < 0.001** |
| Left G_oc-temp_med-Lingual (22) | 69466.07 + 12899.34 (70652.59) | 31268.93 + 7764.06 (42525.49) | 29618.33 + 7506.47 (41114.63) | 17770.00 + 4370.58 (23938.66) | 13969.33 + 5456.86 (29888.45) | 0.0023** |
| Left G_oc-temp_med-Parahip (23) | 41862.33 + 9796.64 (53658.40) | 41384.53 + 8338.12 (45669.78) | 28466.33 + 4438.51 (24310.71) | 19811.60 + 3127.25 (17128.67) | 1506.27 + 982.02 (5378.73) | 0.0025** |
| Left G_pariet_inf-Angular (25) | 98068.00 + 22425.47 (122829.36) | 27804.47 + 4884.88 (26755.60) | 16650.27 + 3040.58 (16653.94) | 19055.67 + 4087.76 (22389.59) | 43204.93 + 10614.68 (58138.97) | < 0.001** |
| Left G_postcentral (28) | 116097.13 + 27145.36 (148681.25) | 35613.33 + 9633.32 (52763.85) | 24237.13 + 7107.45 (38929.11) | 25213.00 + 7376.22 (40401.23) | 41871.00 + 11103.41 (60815.87) | 0.0035** |
| Left G_precentral (29) | 170934.87 + 26909.80 (147391.07) | 59417.27 + 11538.89 (63201.10) | 35258.87 + 5729.28 (31380.55) | 32005.93 + 5076.26 (27803.84) | 24004.80 + 6956.28 (38101.14) | < 0.001** |
| Left G_rectus (31) | 57752.80 + 11331.03 (62062.61) | 44832.00 + 9037.83 (49502.24) | 28576.60 + 7376.67 (40403.71) | 15513.20 + 6767.90 (37069.33) | 5153.53 + 1704.78 (9337.49) | 0.0024** |
| Left G_temp_sup-G_T_transv (33) | 51038.60 + 9510.61 (52091.75) | 19368.47 + 2719.68 (14896.29) | 17839.13 + 3444.43 (18865.91) | 9286.87 + 1348.75 (7387.42) | 331.33 + 136.49 (747.59) | < 0.001** |
| Left G_temp_sup-Lateral (34) | 83950.93 + 11879.40 (65066.14) | 39945.93 + 7011.60 (38404.09) | 41298.47 + 16601.94 (90932.58) | 12945.47 + 2762.52 (15130.93) | 23871.73 + 6904.94 (37819.94) | 0.0034** |
| Left G_temporal_middle (38) | 187415.87 + 28244.91 (154703.74) | 82059.53 + 17024.94 (93249.45) | 43896.00 + 7876.16 (43139.48) | 31837.60 + 6662.65 (36492.85) | 24252.00 + 6890.19 (37739.15) | < 0.001** |
| Left S_circular_insula_ant (47) | 24766.73 + 5261.14 (28816.47) | 36282.00 + 5509.23 (30175.28) | 25445.67 + 4485.30 (24567.01) | 13956.87 + 3600.78 (19722.27) | 3111.53 + 1396.26 (7647.64) | < 0.001** |
| Left S_circular_insula_sup (49) | 48917.93 + 12875.21 (70520.42) | 106401.80 + 16440.63 (90049.06) | 92955.93 + 16929.26 (92725.38) | 35489.13 + 6671.83 (36543.14) | 6466.47 + 3290.52 (18022.93) | < 0.001** |
| Left S_front_inf (52) | 62831.73 + 10360.31 (56745.74) | 59448.40 + 6516.83 (35694.13) | 48399.93 + 6938.10 (38001.54) | 30227.87 + 4508.65 (24694.92) | 16950.27 + 4099.20 (22452.26) | < 0.001** |
| Left S_orbital_med-olfact (63) | 596.67 + 421.62 (2309.28) | 14329.20 + 1963.18 (10752.78) | 18730.47 + 3237.31 (17731.47) | 13094.47 + 2811.08 (15396.94) | 2033.27 + 1853.08 (10149.73) | < 0.001** |
| Left S_temporal_transverse (74) | 14236.27 + 3659.12 (20041.80) | 22790.07 + 4035.57 (22103.73) | 20026.27 + 3043.27 (16668.68) | 12175.60 + 2697.16 (14772.95) | 2630.53 + 1356.82 (7431.60) | 0.0084** |
| Right G_and_S_cingul-Ant (80) | 64216.40 + 11745.09 (64330.52) | 67585.53 + 9000.28 (49296.54) | 41575.13 + 7966.16 (43632.48) | 16670.13 + 3050.32 (16707.30) | 11471.93 + 2845.97 (15588.01) | < 0.001** |
| Right G_and_S_cingul-Mid-Ant (81) | 78713.80 + 10416.55 (57053.79) | 64613.47 + 10780.29 (59046.05) | 31945.73 + 6193.80 (33924.82) | 21900.40 + 3748.59 (20531.87) | 10724.93 + 3149.80 (17252.16) | < 0.001** |
| Right G_and_S_cingul-Mid-Post (82) | 126618.73 + 28915.78 (158378.24) | 93623.27 + 16875.42 (92430.48) | 57820.60 + 13101.10 (71757.68) | 27995.00 + 7217.24 (39530.48) | 15588.13 + 3819.77 (20921.73) | 0.0011** |
| Right G_cingul-Post-dorsal (83) | 35260.33 + 5941.80 (32544.58) | 16464.00 + 3574.12 (19576.29) | 12573.53 + 3781.53 (20712.29) | 7328.67 + 2412.00 (13211.09) | 3385.47 + 2361.34 (12933.58) | < 0.001** |
| Right G_cingul-Post-ventral (84) | 38386.87 + 9970.24 (54609.28) | 28352.87 + 5042.84 (27620.76) | 19435.40 + 2500.57 (13696.21) | 13525.60 + 1753.23 (9602.85) | 7.33 + 5.36 (29.34) | < 0.001** |
| Right G_front_inf-Opercular (86) | 152062.27 + 19806.48 (108484.57) | 28949.33 + 4864.10 (26641.77) | 20385.87 + 4250.10 (23278.77) | 10186.73 + 2443.80 (13385.25) | 3269.87 + 1214.77 (6653.59) | < 0.001** |
| Right G_front_inf-Orbital (87) | 43334.73 + 8432.02 (46184.09) | 23913.67 + 3371.55 (18466.74) | 9534.67 + 1398.99 (7662.56) | 7453.20 + 1589.05 (8703.59) | 937.53 + 373.98 (2048.37) | < 0.001** |
| Right G_front_inf-Triangul (88) | 52534.00 + 7659.68 (41953.79) | 23017.73 + 5431.49 (29749.51) | 21416.40 + 5105.64 (27964.73) | 11666.53 + 1624.36 (8896.98) | 8153.33 + 2783.86 (15247.80) | < 0.001** |
| Right G_front_middle (89) | 214042.27 + 20934.55 (114663.24) | 98732.93 + 13682.04 (74939.64) | 69602.60 + 10903.46 (59720.70) | 50030.07 + 6732.17 (36873.60) | 27503.33 + 4564.23 (24999.32) | < 0.001** |
| Right G_front_sup (90) | 345289.40 + 51019.32 (279444.30) | 148996.13 + 32052.34 (175557.88) | 93208.67 + 14036.72 (76882.28) | 66308.47 + 10686.41 (58531.86) | 81284.80 + 23421.57 (128285.23) | < 0.001** |
| Right G_insular_short (92) | 37965.13 + 10619.85 (58167.32) | 14086.80 + 2894.45 (15853.55) | 9767.13 + 1585.03 (8681.58) | 6942.67 + 1383.97 (7580.31) | 811.33 + 503.80 (2759.43) | < 0.001** |
| Right G_occipital_middle (93) | 159380.87 + 28486.50 (156026.98) | 76258.13 + 8756.12 (47959.26) | 53203.20 + 9637.67 (52787.72) | 37151.67 + 7178.59 (39318.77) | 19012.33 + 4927.31 (26988.00) | < 0.001** |
| Right G_oc-temp_lat-fusifor (95) | 110338.87 + 25381.37 (139019.46) | 52452.33 + 12697.63 (69547.78) | 33206.60 + 10774.91 (59016.59) | 13479.07 + 2749.87 (15061.67) | 1109.27 + 352.26 (1929.42) | < 0.001** |
| Right G_pariet_inf-Angular (99) | 120177.87 + 24578.46 (134621.77) | 50247.67 + 11493.28 (62951.31) | 26490.67 + 4538.14 (24856.40) | 37551.47 + 11515.87 (63075.04) | 30817.40 + 10703.71 (58626.65) | 0.0019** |
| Right G_pariet_inf-Supramar (100) | 109984.67 + 18074.86 (99000.11) | 45252.60 + 8881.63 (48646.71) | 27305.13 + 5030.82 (27554.96) | 23405.27 + 4688.14 (25678.02) | 14616.80 + 4831.58 (26463.64) | < 0.001** |
| Right G_postcentral (102) | 84326.40 + 22112.45 (121114.87) | 17218.27 + 3998.58 (21901.14) | 15242.80 + 3379.61 (18510.90) | 15186.33 + 3873.90 (21218.24) | 13728.80 + 3577.11 (19592.66) | < 0.001** |
| Right G_precentral (103) | 190926.87 + 37351.28 (204581.40) | 66875.33 + 15460.99 (84683.34) | 45879.67 + 12249.57 (67093.64) | 51201.80 + 12121.71 (66393.36) | 34691.80 + 8540.63 (46778.94) | < 0.001** |
| Right G_temp_sup-G_T_transv (107) | 36130.53 + 6608.81 (36197.94) | 16648.60 + 2333.59 (12781.57) | 16394.67 + 2539.55 (13909.67) | 14561.47 + 2598.72 (14233.77) | 765.13 + 424.82 (2326.84) | < 0.001** |
| Right G_temp_sup-Lateral (108) | 121087.73 + 17298.97 (94750.34) | 35942.07 + 6505.30 (35630.99) | 25542.07 + 6751.03 (36976.92) | 10383.07 + 2624.35 (14374.16) | 4302.53 + 1699.32 (9307.54) | < 0.001** |
| Right G_temp_sup-Plan_polar (109) | 24289.13 + 6501.03 (35607.61) | 42593.20 + 8815.28 (48283.29) | 40997.87 + 7500.53 (41082.10) | 22264.40 + 4818.90 (26394.22) | 1445.00 + 777.33 (4257.63) | 0.0035** |
| Right G_temp_sup-Plan_tempo (110) | 42973.00 + 7071.08 (38729.90) | 23677.33 + 3275.76 (17942.08) | 15663.87 + 2486.45 (13618.85) | 8484.20 + 1639.26 (8978.59) | 2561.20 + 790.48 (4329.65) | < 0.001** |
| Right G_temporal_inf (111) | 79664.73 + 11389.51 (62382.92) | 28290.27 + 4688.69 (25680.99) | 18152.93 + 2752.38 (15075.39) | 11442.00 + 2032.76 (11133.87) | 10119.40 + 3380.23 (18514.27) | < 0.001** |
| Right G_temporal_middle (112) | 173375.93 + 26035.41 (142601.79) | 54950.33 + 9840.76 (53900.07) | 33286.60 + 7269.20 (39815.04) | 25858.80 + 6018.67 (32965.62) | 9680.13 + 3832.14 (20989.49) | < 0.001** |
| Right Lat_Fis-ant-Horizont (113) | 18865.07 + 4435.12 (24292.13) | 33603.73 + 4600.21 (25196.39) | 25345.60 + 3046.15 (16684.45) | 16119.67 + 2265.73 (12409.91) | 1435.87 + 582.78 (3192.02) | < 0.001** |
| Right Lat_Fis-ant-Vertical (114) | 15767.00 + 4736.48 (25942.75) | 24265.33 + 4096.57 (22437.84) | 29988.27 + 3615.07 (19800.57) | 16808.27 + 2649.16 (14510.03) | 684.27 + 404.03 (2212.94) | < 0.001** |
| Right S_cingul-Marginalis (120) | 52174.07 + 9268.89 (50767.80) | 72520.80 + 13369.77 (73229.26) | 54375.73 + 9053.30 (49586.99) | 28907.67 + 5690.32 (31167.17) | 12463.80 + 5053.25 (27677.81) | 0.0043** |
| Right S_circular_insula_ant (121) | 19154.87 + 4957.66 (27154.23) | 30905.27 + 5141.69 (28162.20) | 21499.13 + 3704.73 (20291.63) | 11061.87 + 1578.91 (8648.05) | 2136.67 + 824.68 (4516.98) | < 0.001** |
| Right S_circular_insula_inf (122) | 14624.40 + 3654.62 (20017.19) | 29294.47 + 4666.89 (25561.62) | 40747.33 + 6191.92 (33914.53) | 23195.40 + 4683.69 (25653.62) | 7728.40 + 3794.70 (20784.43) | 0.0017** |
| Right S_circular_insula_sup (123) | 46867.93 + 16203.43 (88749.86) | 79768.47 + 19275.33 (105575.31) | 51013.67 + 5742.32 (31451.99) | 18050.33 + 2963.28 (16230.55) | 661.13 + 332.72 (1822.38) | 0.0035** |
| Right S_front_inf (126) | 107645.47 + 17722.77 (97071.63) | 74946.60 + 11637.13 (63739.16) | 45043.93 + 7252.59 (39724.07) | 27807.07 + 4862.20 (26631.35) | 26214.60 + 5548.54 (30390.60) | < 0.001** |
| Right S_front_sup (128) | 79632.47 + 13830.82 (75754.54) | 36168.53 + 7678.46 (42056.68) | 29531.80 + 5614.97 (30754.47) | 26537.53 + 4818.88 (26394.08) | 29881.60 + 9005.54 (49325.38) | 0.0083** |
| Right S_precentral-inf-part (142) | 57704.07 + 9423.79 (51616.23) | 30839.20 + 5082.89 (27840.12) | 17550.73 + 4074.14 (22315.01) | 14715.93 + 2876.74 (15756.57) | 11496.40 + 2844.94 (15582.39) | < 0.001** |
| Right S_temporal_sup (147) | 174853.93 + 26896.66 (147319.07) | 118394.60 + 18811.82 (103036.58) | 75028.93 + 18497.30 (101313.89) | 49068.60 + 8470.57 (46395.23) | 66687.87 + 18400.09 (100781.46) | 0.0035** |
| Right S_temporal_transverse (148) | 30180.20 + 6426.10 (35197.21) | 34518.00 + 7055.62 (38645.19) | 26316.60 + 4549.83 (24920.47) | 15185.87 + 2159.35 (11827.24) | 669.53 + 412.91 (2261.62) | < 0.001** |
| Clustering coefficient | | | | | | |
| Left G_oc-temp_lat-fusifor (21) | 6.92 + 0.29 (1.57) | 7.23 + 0.28 (1.53) | 7.52 + 0.26 (1.40) | 7.96 + 0.24 (1.33) | 8.71 + 0.26 (1.45) | 0.0064** |
| Right G_front_inf-Opercular (86) | 5.84 + 0.30 (1.66) | 6.51 + 0.30 (1.64) | 7.11 + 0.31 (1.67) | 7.80 + 0.34 (1.88) | 8.67 + 0.42 (2.29) | < 0.001** |
| Right G_front_inf-Orbital (87) | 6.35 + 0.29 (1.56) | 6.70 + 0.25 (1.38) | 7.40 + 0.29 (1.56) | 7.84 + 0.36 (1.94) | 8.56 + 0.39 (2.12) | 0.0016** |
| Right G_oc-temp_lat-fusifor (95) | 6.62 + 0.22 (1.23) | 6.77 + 0.21 (1.13) | 6.98 + 0.20 (1.10) | 7.45 + 0.20 (1.07) | 8.33 + 0.24 (1.29) | < 0.001** |
| Right G_temp_sup-Lateral (108) | 6.26 + 0.30 (1.63) | 6.69 + 0.30 (1.66) | 7.20 + 0.32 (1.75) | 7.64 + 0.31 (1.70) | 8.28 + 0.31 (1.69) | 0.0096** |
| Right G_temp_sup-Plan_tempo (110) | 6.82 + 0.24 (1.29) | 6.90 + 0.24 (1.34) | 7.21 + 0.26 (1.44) | 7.83 + 0.28 (1.51) | 8.67 + 0.32 (1.76) | < 0.001** |
| Right Lat_Fis-post (115) | 6.30 + 0.22 (1.21) | 6.09 + 0.20 (1.08) | 6.20 + 0.20 (1.07) | 6.78 + 0.23 (1.25) | 7.63 + 0.30 (1.63) | 0.0025** |
| Participation coefficient | | | | | | |
| Left G_cingul-Post-ventral (10) | 15.53 + 1.03 (5.66) | 15.15 + 0.99 (5.44) | 13.13 + 1.06 (5.80) | 10.37 + 1.14 (6.24) | 7.41 + 1.12 (6.16) | < 0.001** |
| Left G_oc-temp_med-Parahip (23) | 14.71 + 1.22 (6.69) | 13.78 + 1.17 (6.43) | 11.82 + 1.29 (7.08) | 8.88 + 1.20 (6.58) | 5.26 + 0.97 (5.33) | < 0.001** |
| Left S_circular_insula_sup (49) | 16.00 + 1.02 (5.56) | 16.91 + 0.95 (5.18) | 16.50 + 0.95 (5.23) | 13.75 + 1.09 (5.95) | 10.60 + 1.10 (6.00) | 0.0094** |
| Right G_cingul-Post-ventral (84) | 14.61 + 1.28 (7.04) | 14.62 + 1.19 (6.52) | 13.14 + 1.22 (6.68) | 10.81 + 1.20 (6.56) | 7.28 + 1.07 (5.83) | 0.0072** |
| Right G_oc-temp_med-Parahip (97) | 11.51 + 1.21 (6.65) | 10.33 + 1.26 (6.91) | 8.30 + 1.15 (6.32) | 6.83 + 1.15 (6.30) | 3.91 + 0.95 (5.18) | 0.0074** |
| Right Lat_Fis-ant-Vertical (114) | 9.76 + 1.04 (5.67) | 9.88 + 1.10 (6.05) | 8.38 + 1.07 (5.89) | 5.57 + 0.98 (5.34) | 3.62 + 0.90 (4.91) | 0.0032** |
| Right S_circular_insula_sup (123) | 15.04 + 0.95 (5.21) | 15.28 + 1.03 (5.66) | 13.92 + 1.15 (6.31) | 10.59 + 1.23 (6.74) | 7.60 + 1.15 (6.31) | < 0.001** |

| **Table S8.** Area-under-the-curve (AUC) values for each global measure for the between-layer analysis. NaN values are due to global s that cannot be derived for specific layers. Data are shown as mean ± standard error (standard deviation). One-way ANOVA was used to compare layers with an FDR correction (alpha = 0.05). Graph density and average degree centrality do not have standard error and standard deviation since they are consistent across all participants due to thresholding. Shaded rows have significant differences between layers. P values shown are false discovery rate corrected (FDR, Benjamini-Hochberg method, alpha = 0.05).  * p < 0.05, ** p < 0.01. | | | | | | | |
| --- | --- | --- | --- | --- | --- | --- | --- |
| Measures (AUC) | Layer 1 (Superficial) | Layer 2 | Layer 3 | Layer 4 | Layer 5 (Deep) | Multilayer | p value |
| Modularity | NaN | NaN | NaN | NaN | NaN | 13.74 + 0.57 (3.14) | NaN |
| Transitivity | NaN | NaN | NaN | NaN | NaN | 5.16 + 0.19 (1.03) | NaN |
| Largest Cluster  Size | NaN | NaN | NaN | NaN | NaN | 26641.40 + 202.95 (1111.58) | NaN |
| Graph Density | NaN | NaN | NaN | NaN | NaN | 6.33 + 0.01 (0.05) | NaN |
| Characteristic  Path Length | NaN | NaN | NaN | NaN | NaN | 363.49 + 8.04 (44.03) | NaN |
| Global Efficiency | NaN | NaN | NaN | NaN | NaN | 5.06 + 0.12 (0.64) | NaN |
| Radius | NaN | NaN | NaN | NaN | NaN | 277.38 + 20.37 (111.56) | NaN |
| Diameter | NaN | NaN | NaN | NaN | NaN | 1283.42 + 31.67 (173.45) | NaN |
| Assortativity | NaN | NaN | NaN | NaN | NaN | 10.34 + 0.49 (2.69) | NaN |
| Avg. Degree Centrality | 4748.31 + 69.28 (379.49) | 4903.28 + 38.52 (210.96) | 4870.76 + 15.08 (82.60) | 4627.74 + 37.58 (205.85) | 4237.92 + 73.16 (400.72) | 4677.60 + 6.68 (36.56) | < 0.001** |
| Avg. Strength | 1145.33 + 29.00 (158.84) | 1204.12 + 28.97 (158.68) | 1207.09 + 31.67 (173.45) | 1144.67 + 33.91 (185.71) | 1032.98 + 37.04 (202.88) | 1146.84 + 28.45 (155.81) | 0.0021** |
| Avg. Eigenvector  Centrality | 0.98 + 0.03 (0.14) | 1.02 + 0.02 (0.11) | 1.03 + 0.01 (0.08) | 0.99 + 0.01 (0.06) | 0.91 + 0.02 (0.09) | 0.98 + 0.01 (0.06) | < 0.001** |
| Avg. Betweenness  Centrality | 46777.98 + 1942.31 (10638.44) | 54155.59 + 1315.20 (7203.67) | 39616.03 + 1012.14 (5543.72) | 27019.47 + 1407.76 (7710.59) | 14833.07 + 1259.23 (6897.06) | 36480.43 + 572.22 (3134.15) | < 0.001** |
| Avg. Clustering  Coefficient | 5.53 + 0.20 (1.07) | 5.37 + 0.18 (0.97) | 5.46 + 0.16 (0.89) | 5.73 + 0.16 (0.87) | 6.28 + 0.17 (0.94) | 5.67 + 0.17 (0.92) | 0.0033** |
| Avg. Local  Efficiency | 7.78 + 0.20 (1.07) | 7.74 + 0.19 (1.04) | 7.85 + 0.19 (1.02) | 7.94 + 0.18 (1.00) | 8.18 + 0.19 (1.04) | 7.90 + 0.19 (1.02) | 0.50 |
| Avg. Participation  Coefficient | 12.89 + 0.61 (3.35) | 12.85 + 0.62 (3.37) | 12.38 + 0.61 (3.34) | 11.58 + 0.59 (3.24) | 10.51 + 0.56 (3.06) | 12.04 + 0.59 (3.21) | 0.032* |

| **Table S9.** Area-under-the-curve (AUC) values for each significant nodal measure for between-layer analysis. Data are shown as mean ± standard error (standard deviation). One-way ANOVA was used to compare layers. Node long names can be found in Table S1. P values shown are false discovery rate corrected (FDR, Bonferroni-Holm method, alpha = 0.01).  * p < 0.05, ** p < 0.01. | | | | | | |
| --- | --- | --- | --- | --- | --- | --- |
| Node | Layer 1 (Superficial) | Layer 2 | Layer 3 | Layer 4 | Layer 5 (Deep) | P value |
| Degree centrality | | | | | | |
| Left G_cingul-Post-ventral (10) | 3841.03 + 329.56 (1805.05) | 3849.77 + 350.81 (1921.47) | 3183.27 + 322.75 (1767.77) | 2306.63 + 304.62 (1668.47) | 1348.33 + 237.56 (1301.17) | < 0.001** |
| Right G_cingul-Post-ventral (84) | 3857.83 + 409.31 (2241.89) | 3888.27 + 369.22 (2022.28) | 3143.00 + 352.19 (1929.02) | 2278.80 + 322.49 (1766.33) | 1316.43 + 238.51 (1306.39) | < 0.001** |
| Right G_temp_sup-G_T_transv (107) | 4040.07 + 417.16 (2284.87) | 3746.53 + 410.20 (2246.77) | 3295.87 + 363.96 (1993.52) | 2601.10 + 334.77 (1833.60) | 1670.87 + 280.79 (1537.97) | 0.0068** |
| Right S_circular_insula_sup (123) | 4437.30 + 392.73 (2151.06) | 5312.27 + 374.22 (2049.67) | 5345.47 + 338.21 (1852.44) | 3445.20 + 363.29 (1989.82) | 1958.43 + 316.42 (1733.13) | < 0.001** |
| Right S_temporal_transverse (148) | 3578.70 + 447.70 (2452.15) | 3555.40 + 445.31 (2439.09) | 2704.37 + 343.23 (1879.94) | 1860.73 + 326.43 (1787.91) | 1083.47 + 246.27 (1348.90) | < 0.001** |
| Strength | | | | | | |
| Left G_cingul-Post-ventral (10) | 812.72 + 74.37 (407.33) | 821.92 + 82.64 (452.65) | 664.49 + 74.04 (405.52) | 461.91 + 64.09 (351.02) | 250.19 + 44.38 (243.06) | < 0.001** |
| Right G_cingul-Post-ventral (84) | 838.94 + 96.61 (529.15) | 839.09 + 90.04 (493.18) | 666.23 + 84.90 (464.99) | 471.86 + 74.48 (407.95) | 262.40 + 53.86 (295.02) | < 0.001** |
| Right G_temp_sup-G_T_transv (107) | 887.12 + 98.99 (542.19) | 825.37 + 97.21 (532.42) | 716.58 + 88.47 (484.59) | 553.64 + 76.77 (420.48) | 339.79 + 61.55 (337.10) | 0.0086** |
| Right S_circular_insula_sup (123) | 982.01 + 98.63 (540.19) | 1196.99 + 93.44 (511.79) | 1199.38 + 83.28 (456.17) | 751.67 + 90.64 (496.48) | 412.65 + 76.81 (420.68) | < 0.001** |
| Right S_temporal_transverse (148) | 756.41 + 101.77 (557.41) | 763.94 + 101.97 (558.52) | 568.57 + 79.29 (434.27) | 376.88 + 67.49 (369.67) | 201.84 + 42.29 (231.66) | < 0.001** |
| Eigenvector centrality | | | | | | |
| Left G_cingul-Post-ventral (10) | 0.69 + 0.08 (0.46) | 0.69 + 0.09 (0.47) | 0.55 + 0.08 (0.42) | 0.38 + 0.07 (0.37) | 0.22 + 0.05 (0.26) | 0.0018** |
| Right G_cingul-Post-ventral (84) | 0.75 + 0.10 (0.56) | 0.72 + 0.09 (0.50) | 0.56 + 0.08 (0.44) | 0.39 + 0.07 (0.38) | 0.21 + 0.05 (0.27) | 0.0015** |
| Right S_circular_insula_sup (123) | 0.89 + 0.12 (0.63) | 1.08 + 0.12 (0.66) | 1.08 + 0.11 (0.61) | 0.67 + 0.10 (0.55) | 0.36 + 0.08 (0.42) | < 0.001** |
| Betweenness centrality | | | | | | |
| Left G_and_S_cingul-Mid-Post (8) | 75702.67 + 15316.89 (83894.08) | 116668.93 + 17455.29 (95606.56) | 64943.20 + 11393.84 (62406.64) | 30863.40 + 6692.49 (36656.26) | 12208.07 + 3419.26 (18728.07) | < 0.001** |
| Left G_cingul-Post-dorsal (9) | 38577.27 + 9521.32 (52150.44) | 43946.80 + 10765.32 (58964.11) | 14194.20 + 3648.27 (19982.39) | 8228.47 + 1894.50 (10376.59) | 3158.20 + 2251.49 (12331.90) | 0.0018** |
| Left G_cingul-Post-ventral (10) | 32904.73 + 6443.48 (35292.41) | 38474.00 + 6997.18 (38325.12) | 20040.73 + 3049.83 (16704.58) | 15662.33 + 2105.77 (11533.80) | 143.00 + 118.72 (650.24) | < 0.001** |
| Left G_front_inf-Opercular (12) | 73589.33 + 11669.80 (63918.15) | 43739.53 + 7197.94 (39424.74) | 25929.73 + 4865.49 (26649.37) | 12373.00 + 2701.49 (14796.70) | 3848.67 + 1458.55 (7988.83) | < 0.001** |
| Left G_front_inf-Orbital (13) | 39247.60 + 7283.20 (39891.76) | 23116.00 + 3341.05 (18299.68) | 11865.87 + 1862.56 (10201.66) | 9916.87 + 1461.56 (8005.31) | 4184.53 + 2431.42 (13317.41) | < 0.001** |
| Left G_front_middle (15) | 138701.53 + 23136.59 (126724.33) | 139929.07 + 15424.83 (84485.27) | 103788.73 + 12776.50 (69979.75) | 77956.67 + 11602.30 (63548.42) | 34907.53 + 5922.10 (32436.69) | < 0.001** |
| Left G_front_sup (16) | 225676.67 + 29970.60 (164155.76) | 213360.13 + 22275.09 (122005.67) | 118380.20 + 14580.33 (79859.74) | 83235.80 + 9088.32 (49778.78) | 76423.13 + 14209.82 (77830.39) | < 0.001** |
| Left G_insular_short (18) | 29407.80 + 6153.58 (33704.53) | 18947.80 + 3552.32 (19456.83) | 9035.47 + 1675.70 (9178.17) | 9099.00 + 1548.31 (8480.43) | 1930.47 + 1058.46 (5797.44) | < 0.001** |
| Left G_oc-temp_lat-fusifor (21) | 68827.13 + 13646.22 (74743.40) | 55690.80 + 14512.29 (79487.06) | 28297.33 + 5224.97 (28618.34) | 15889.47 + 4201.53 (23012.74) | 4454.60 + 1810.77 (9918.00) | < 0.001** |
| Left G_oc-temp_med-Parahip (23) | 37669.47 + 8039.60 (44034.70) | 41599.93 + 7574.38 (41486.57) | 29273.73 + 4361.26 (23887.61) | 20158.07 + 2951.38 (16165.35) | 1459.40 + 1005.31 (5506.32) | < 0.001** |
| Left G_precentral (29) | 125074.00 + 20356.32 (111496.18) | 87534.07 + 14575.36 (79832.53) | 51056.73 + 6922.60 (37916.65) | 41199.53 + 6284.88 (34423.68) | 20960.47 + 5480.23 (30016.44) | < 0.001** |
| Left G_precuneus (30) | 121827.80 + 21297.97 (116653.78) | 148253.67 + 14325.09 (78461.74) | 155588.07 + 19908.51 (109043.41) | 115986.67 + 17019.13 (93217.61) | 45811.13 + 10390.61 (56911.72) | 0.0072** |
| Left G_rectus (31) | 49758.40 + 8762.13 (47992.16) | 50778.00 + 8413.47 (46082.47) | 33679.60 + 7209.15 (39486.14) | 16681.53 + 6334.71 (34696.63) | 4908.93 + 1725.57 (9451.33) | < 0.001** |
| Left G_temp_sup-G_T_transv (33) | 44577.47 + 7192.04 (39392.42) | 22894.47 + 2812.57 (15405.09) | 18342.73 + 3178.58 (17409.82) | 9582.40 + 1155.84 (6330.82) | 979.20 + 410.38 (2247.76) | < 0.001** |
| Left G_temporal_middle (38) | 146013.60 + 23001.56 (125984.76) | 110814.07 + 18063.21 (98936.26) | 58867.87 + 9263.71 (50739.42) | 40733.87 + 7749.29 (42444.63) | 19833.53 + 4697.76 (25730.68) | < 0.001** |
| Left Lat_Fis-ant-Vertical (40) | 15182.00 + 4762.28 (26084.08) | 26746.73 + 5200.90 (28486.48) | 17831.87 + 3012.40 (16499.58) | 12839.00 + 1971.15 (10796.44) | 1397.93 + 621.08 (3401.82) | 0.0059** |
| Left S_cingul-Marginalis (46) | 35560.93 + 7747.29 (42433.66) | 87018.40 + 16690.23 (91416.14) | 62532.53 + 7693.19 (42137.34) | 31877.67 + 6749.48 (36968.45) | 5717.47 + 1707.92 (9354.67) | < 0.001** |
| Left S_circular_insula_ant (47) | 22985.20 + 4640.74 (25418.36) | 35558.53 + 5096.53 (27914.86) | 25872.33 + 4116.32 (22546.02) | 14273.87 + 3854.08 (21109.66) | 3291.47 + 1474.28 (8074.96) | < 0.001** |
| Left S_circular_insula_sup (49) | 45641.20 + 13876.54 (76004.93) | 114592.33 + 17161.34 (93996.54) | 100037.20 + 15239.67 (83471.11) | 39018.87 + 7610.79 (41685.99) | 7777.07 + 3736.63 (20466.35) | < 0.001** |
| Left S_front_inf (52) | 46978.00 + 7706.07 (42207.87) | 73656.53 + 7682.63 (42079.50) | 56670.33 + 7440.35 (40752.50) | 36598.60 + 4754.43 (26041.10) | 15051.80 + 3372.90 (18474.13) | < 0.001** |
| Left S_occipital_ant (59) | 16028.07 + 4141.87 (22685.95) | 28534.40 + 5201.06 (28487.36) | 29355.47 + 4815.49 (26375.50) | 12761.00 + 1635.56 (8958.32) | 6905.60 + 2521.90 (13813.04) | 0.0083** |
| Left S_oc-temp_med_and_Lingual (61) | 54830.93 + 11201.79 (61354.72) | 93648.20 + 16667.08 (91289.37) | 64551.60 + 15222.25 (83375.67) | 28342.80 + 6541.65 (35830.07) | 11213.13 + 3685.45 (20186.04) | 0.0018** |
| Left S_orbital_med-olfact (63) | 672.00 + 443.23 (2427.68) | 14931.47 + 2050.08 (11228.73) | 19011.33 + 2960.52 (16215.42) | 13536.13 + 2785.66 (15257.66) | 1973.20 + 1862.66 (10202.20) | < 0.001** |
| Left S_suborbital (70) | 12237.33 + 3363.73 (18423.92) | 26963.20 + 4342.88 (23786.93) | 34860.93 + 5771.77 (31613.27) | 25693.13 + 4452.08 (24385.07) | 8704.27 + 2521.95 (13813.26) | 0.0060** |
| Left S_temporal_transverse (74) | 11475.07 + 3275.49 (17940.62) | 24588.33 + 3801.26 (20820.34) | 21991.53 + 3033.11 (16613.03) | 12302.00 + 2367.54 (12967.55) | 2215.53 + 1078.86 (5909.19) | < 0.001** |
| Right G_and_S_cingul-Ant (80) | 55586.93 + 9315.19 (51021.40) | 74826.00 + 9236.73 (50591.64) | 47476.27 + 8035.36 (44011.45) | 18669.27 + 2788.06 (15270.83) | 9350.33 + 2283.61 (12507.86) | < 0.001** |
| Right G_and_S_cingul-Mid-Ant (81) | 65371.73 + 8939.97 (48966.23) | 74850.20 + 10919.41 (59808.07) | 39805.93 + 6423.38 (35182.32) | 25395.40 + 4188.51 (22941.44) | 10791.80 + 2974.46 (16291.80) | < 0.001** |
| Right G_and_S_cingul-Mid-Post (82) | 113381.33 + 29726.58 (162819.16) | 122612.87 + 20248.73 (110906.88) | 74585.27 + 13976.73 (76553.69) | 34606.20 + 7473.26 (40932.72) | 17887.00 + 4195.16 (22977.81) | 0.0041** |
| Right G_cingul-Post-dorsal (83) | 23924.87 + 4250.26 (23279.61) | 23338.00 + 4047.51 (22169.10) | 16097.13 + 3797.57 (20800.12) | 9110.20 + 3240.15 (17747.05) | 3210.33 + 2138.55 (11713.33) | 0.0092** |
| Right G_cingul-Post-ventral (84) | 28772.07 + 6950.96 (38071.97) | 32604.80 + 5563.76 (30473.95) | 19917.33 + 2536.21 (13891.39) | 13543.93 + 1749.60 (9582.93) | 6.13 + 4.67 (25.57) | < 0.001** |
| Right G_front_inf-Opercular (86) | 114503.87 + 14029.92 (76845.05) | 50750.13 + 7017.15 (38434.53) | 23137.67 + 3752.09 (20551.06) | 12887.47 + 3035.17 (16624.30) | 2561.27 + 960.13 (5258.86) | < 0.001** |
| Right G_front_inf-Orbital (87) | 32034.60 + 6652.63 (36437.97) | 29211.07 + 4774.17 (26149.18) | 11475.40 + 1574.31 (8622.87) | 8079.73 + 1524.26 (8348.73) | 815.07 + 311.56 (1706.51) | < 0.001** |
| Right G_front_inf-Triangul (88) | 40368.67 + 6081.42 (33309.30) | 31831.73 + 5742.61 (31453.58) | 23557.00 + 4908.63 (26885.65) | 14391.67 + 1873.30 (10260.50) | 8488.27 + 2697.02 (14772.17) | < 0.001** |
| Right G_front_middle (89) | 148545.87 + 14683.05 (80422.35) | 146963.40 + 14169.56 (77609.90) | 98000.87 + 11164.55 (61150.78) | 61967.33 + 7004.93 (38367.58) | 24693.67 + 3439.13 (18836.87) | < 0.001** |
| Right G_front_sup (90) | 247404.40 + 37106.55 (203240.96) | 218354.00 + 36669.06 (200844.69) | 134093.00 + 17285.60 (94677.11) | 89596.73 + 13359.44 (73172.65) | 68420.93 + 18488.22 (101264.14) | < 0.001** |
| Right G_insular_short (92) | 34237.87 + 9422.57 (51609.53) | 17261.93 + 3217.12 (17620.89) | 10250.00 + 1653.84 (9058.43) | 6844.20 + 1426.84 (7815.10) | 847.47 + 520.46 (2850.69) | < 0.001** |
| Right G_occipital_middle (93) | 124539.47 + 23477.59 (128592.04) | 106470.73 + 11727.07 (64231.80) | 73541.20 + 10317.94 (56513.66) | 45579.60 + 7462.79 (40875.40) | 17482.53 + 4823.35 (26418.56) | < 0.001** |
| Right G_oc-temp_lat-fusifor (95) | 75604.20 + 16044.66 (87880.21) | 66715.80 + 11888.84 (65117.87) | 49830.67 + 13981.05 (76577.37) | 15404.20 + 3733.42 (20448.81) | 1206.93 + 366.23 (2005.94) | < 0.001** |
| Right G_pariet_inf-Supramar (100) | 79013.33 + 13253.20 (72590.78) | 73896.33 + 10674.04 (58464.15) | 41266.40 + 6132.62 (33589.76) | 28127.73 + 5217.09 (28575.20) | 13023.40 + 3732.64 (20444.50) | < 0.001** |
| Right G_temp_sup-G_T_transv (107) | 28688.27 + 4889.35 (26780.07) | 20297.13 + 2232.21 (12226.34) | 17430.13 + 2489.94 (13637.99) | 14177.80 + 2140.06 (11721.61) | 504.13 + 352.50 (1930.71) | < 0.001** |
| Right G_temp_sup-Lateral (108) | 95378.27 + 12848.63 (70374.86) | 53867.53 + 8279.99 (45351.38) | 31076.40 + 7279.92 (39873.75) | 12418.07 + 2668.68 (14616.96) | 4733.20 + 1718.23 (9411.12) | < 0.001** |
| Right G_temp_sup-Plan_polar (109) | 21923.07 + 6344.25 (34748.89) | 45282.73 + 8844.36 (48442.56) | 40286.27 + 7278.42 (39865.53) | 22238.73 + 4242.92 (23239.45) | 1997.93 + 975.54 (5343.23) | 0.0011** |
| Right G_temp_sup-Plan_tempo (110) | 33089.20 + 5319.40 (29135.57) | 31309.07 + 3734.48 (20454.60) | 19680.73 + 2898.01 (15873.04) | 10149.47 + 1575.88 (8631.48) | 2004.87 + 653.60 (3579.92) | < 0.001** |
| Right G_temporal_inf (111) | 69143.53 + 9399.90 (51485.38) | 42634.33 + 6911.29 (37854.70) | 24553.87 + 4122.87 (22581.91) | 16396.87 + 2903.24 (15901.67) | 7131.87 + 2271.50 (12441.52) | < 0.001** |
| Right G_temporal_middle (112) | 131183.80 + 19611.96 (107419.13) | 84549.53 + 11877.18 (65053.99) | 43504.47 + 8081.22 (44262.65) | 30459.13 + 6207.62 (34000.54) | 8839.93 + 3362.81 (18418.85) | < 0.001** |
| Right Lat_Fis-ant-Horizont (113) | 17184.80 + 4099.31 (22452.86) | 37715.67 + 5027.95 (27539.24) | 27527.27 + 3130.95 (17148.95) | 16467.27 + 2225.48 (12189.48) | 1730.87 + 776.08 (4250.75) | < 0.001** |
| Right Lat_Fis-ant-Vertical (114) | 13647.87 + 3235.10 (17719.40) | 28609.60 + 4052.91 (22198.72) | 32318.67 + 3634.36 (19906.22) | 16874.87 + 2592.08 (14197.38) | 720.73 + 400.14 (2191.63) | < 0.001** |
| Right Lat_Fis-post (115) | 83925.53 + 15214.60 (83333.80) | 168775.27 + 27786.47 (152192.79) | 128476.47 + 24449.97 (133917.97) | 49067.87 + 16505.06 (90401.95) | 19625.33 + 9825.30 (53815.39) | < 0.001** |
| Right S_cingul-Marginalis (120) | 38226.27 + 6712.50 (36765.89) | 84931.47 + 11904.38 (65203.00) | 70800.67 + 9453.11 (51776.84) | 33281.40 + 6280.68 (34400.70) | 12601.73 + 4438.10 (24308.46) | < 0.001** |
| Right S_circular_insula_ant (121) | 16236.87 + 3566.65 (19535.36) | 32662.53 + 4426.03 (24242.34) | 24257.20 + 3856.58 (21123.38) | 11537.07 + 1578.84 (8647.64) | 2259.47 + 839.51 (4598.21) | < 0.001** |
| Right S_circular_insula_inf (122) | 10513.67 + 3048.31 (16696.26) | 31816.20 + 4983.51 (27295.79) | 39497.73 + 6151.45 (33692.88) | 20505.60 + 3337.07 (18277.89) | 8525.53 + 4071.07 (22298.14) | < 0.001** |
| Right S_circular_insula_sup (123) | 42860.60 + 16598.59 (90914.22) | 88481.13 + 17289.14 (94696.50) | 57945.80 + 6283.53 (34416.30) | 20047.27 + 3232.24 (17703.72) | 463.67 + 203.18 (1112.89) | < 0.001** |
| Right S_front_inf (126) | 75484.53 + 11028.44 (60405.23) | 102940.47 + 13079.21 (71637.78) | 58452.93 + 8653.86 (47399.15) | 33091.60 + 5159.09 (28257.49) | 24088.87 + 5169.06 (28312.09) | < 0.001** |
| Right S_orbital_lateral (136) | 2703.67 + 915.52 (5014.51) | 13029.40 + 2699.89 (14787.92) | 14988.67 + 2018.57 (11056.15) | 13860.60 + 1890.52 (10354.82) | 5205.60 + 2013.91 (11030.65) | < 0.001** |
| Right S_precentral-inf-part (142) | 43195.27 + 7178.76 (39319.68) | 42715.53 + 6119.43 (33517.52) | 21427.00 + 4194.98 (22976.84) | 17019.00 + 3368.91 (18452.30) | 10655.47 + 2894.40 (15853.27) | < 0.001** |
| Right S_subparietal (145) | 17806.80 + 3214.60 (17607.08) | 48180.80 + 6639.12 (36363.95) | 60487.53 + 9260.85 (50723.74) | 41369.80 + 10265.39 (56225.88) | 14559.07 + 4233.04 (23185.34) | < 0.001** |
| Right S_temporal_transverse (148) | 22436.53 + 4766.62 (26107.85) | 39922.20 + 7980.10 (43708.80) | 28496.27 + 4692.51 (25701.92) | 15329.33 + 2070.92 (11342.90) | 553.60 + 281.58 (1542.26) | < 0.001** |
| Clustering coefficient | | | | | | |
| Left G_cingul-Post-dorsal (9) | 5.11 + 0.21 (1.13) | 5.09 + 0.21 (1.14) | 5.30 + 0.18 (1.01) | 5.61 + 0.20 (1.07) | 6.40 + 0.24 (1.30) | 0.0052** |
| Left G_oc-temp_lat-fusifor (21) | 5.33 + 0.24 (1.34) | 5.47 + 0.23 (1.28) | 5.70 + 0.20 (1.09) | 6.21 + 0.18 (1.01) | 7.17 + 0.26 (1.40) | < 0.001** |
| Left G_temp_sup-G_T_transv (33) | 4.94 + 0.24 (1.32) | 5.35 + 0.23 (1.26) | 5.61 + 0.26 (1.40) | 6.28 + 0.30 (1.65) | 7.55 + 0.46 (2.50) | < 0.001** |
| Left Lat_Fis-post (41) | 5.04 + 0.25 (1.37) | 4.81 + 0.21 (1.15) | 5.09 + 0.23 (1.28) | 5.80 + 0.31 (1.70) | 6.92 + 0.45 (2.47) | < 0.001** |
| Left S_circular_insula_sup (49) | 5.00 + 0.24 (1.33) | 4.52 + 0.20 (1.10) | 4.62 + 0.19 (1.02) | 5.14 + 0.19 (1.04) | 6.11 + 0.24 (1.32) | < 0.001** |
| Left S_temporal_transverse (74) | 5.41 + 0.29 (1.60) | 5.14 + 0.25 (1.38) | 5.40 + 0.27 (1.48) | 6.17 + 0.35 (1.92) | 7.36 + 0.47 (2.60) | 0.0034** |
| Right G_and_S_subcentral (78) | 5.47 + 0.22 (1.22) | 5.52 + 0.22 (1.23) | 5.88 + 0.28 (1.54) | 6.56 + 0.36 (1.98) | 7.65 + 0.48 (2.65) | 0.0017** |
| Right G_cingul-Post-dorsal (83) | 5.28 + 0.20 (1.08) | 5.26 + 0.19 (1.03) | 5.38 + 0.17 (0.91) | 5.73 + 0.17 (0.93) | 6.43 + 0.20 (1.10) | 0.0040** |
| Right G_cingul-Post-ventral (84) | 5.26 + 0.20 (1.09) | 5.14 + 0.18 (1.01) | 5.33 + 0.18 (0.98) | 5.62 + 0.27 (1.50) | 6.76 + 0.37 (2.05) | 0.0058** |
| Right G_front_inf-Opercular (86) | 4.66 + 0.25 (1.40) | 5.10 + 0.25 (1.36) | 5.69 + 0.28 (1.51) | 6.41 + 0.36 (1.95) | 7.39 + 0.45 (2.45) | < 0.001** |
| Right G_front_inf-Orbital (87) | 5.23 + 0.22 (1.22) | 5.41 + 0.22 (1.21) | 6.06 + 0.28 (1.54) | 6.59 + 0.37 (2.02) | 7.52 + 0.40 (2.17) | < 0.001** |
| Right G_oc-temp_lat-fusifor (95) | 5.11 + 0.21 (1.14) | 5.08 + 0.19 (1.02) | 5.26 + 0.16 (0.90) | 5.81 + 0.15 (0.80) | 6.94 + 0.21 (1.16) | < 0.001** |
| Right G_temp_sup-G_T_transv (107) | 4.94 + 0.21 (1.17) | 4.98 + 0.23 (1.25) | 5.37 + 0.20 (1.12) | 5.79 + 0.25 (1.38) | 6.96 + 0.39 (2.13) | < 0.001** |
| Right G_temp_sup-Lateral (108) | 4.88 + 0.24 (1.32) | 5.11 + 0.24 (1.31) | 5.55 + 0.25 (1.39) | 6.02 + 0.25 (1.37) | 6.77 + 0.29 (1.60) | < 0.001** |
| Right G_temp_sup-Plan_tempo (110) | 5.28 + 0.24 (1.29) | 5.18 + 0.22 (1.20) | 5.51 + 0.23 (1.25) | 6.23 + 0.26 (1.41) | 7.39 + 0.36 (1.95) | < 0.001** |
| Right G_temporal_inf (111) | 5.39 + 0.24 (1.31) | 5.39 + 0.22 (1.23) | 5.67 + 0.21 (1.15) | 6.17 + 0.23 (1.25) | 6.90 + 0.25 (1.35) | 0.0015** |
| Right Lat_Fis-post (115) | 4.83 + 0.23 (1.25) | 4.44 + 0.18 (0.96) | 4.60 + 0.15 (0.84) | 5.25 + 0.20 (1.12) | 6.27 + 0.29 (1.61) | < 0.001** |
| Right S_circular_insula_ant (121) | 5.94 + 0.28 (1.55) | 5.23 + 0.21 (1.16) | 5.28 + 0.20 (1.10) | 5.89 + 0.23 (1.24) | 6.92 + 0.30 (1.64) | 0.0018** |
| Right S_circular_insula_sup (123) | 5.02 + 0.22 (1.23) | 4.53 + 0.15 (0.80) | 4.75 + 0.13 (0.73) | 5.62 + 0.22 (1.21) | 6.85 + 0.41 (2.24) | < 0.001** |
| Right S_collat_transv_post (125) | 6.02 + 0.21 (1.13) | 6.00 + 0.20 (1.08) | 6.26 + 0.17 (0.93) | 6.68 + 0.20 (1.07) | 7.37 + 0.28 (1.53) | 0.0030** |
| Right S_occipital_ant (133) | 5.64 + 0.18 (1.00) | 5.48 + 0.15 (0.81) | 5.62 + 0.17 (0.95) | 6.08 + 0.20 (1.08) | 6.94 + 0.26 (1.43) | < 0.001** |
| Right S_oc-temp_med_and_Lingual (135) | 5.28 + 0.18 (0.98) | 5.12 + 0.18 (0.97) | 5.25 + 0.17 (0.93) | 5.64 + 0.19 (1.01) | 6.37 + 0.25 (1.38) | 0.0068** |
| Participation coefficient | | | | | | |
| Left G_cingul-Post-ventral (10) | 15.25 + 1.03 (5.65) | 15.03 + 1.01 (5.52) | 13.11 + 1.07 (5.84) | 10.41 + 1.12 (6.15) | 7.15 + 1.11 (6.06) | < 0.001** |
| Left G_oc-temp_med-Parahip (23) | 14.70 + 1.23 (6.73) | 13.68 + 1.20 (6.57) | 11.76 + 1.29 (7.04) | 8.90 + 1.20 (6.56) | 5.39 + 0.96 (5.27) | < 0.001** |
| Right G_oc-temp_med-Parahip (97) | 11.30 + 1.23 (6.74) | 10.19 + 1.23 (6.76) | 8.30 + 1.15 (6.31) | 6.60 + 1.14 (6.26) | 3.82 + 0.95 (5.22) | 0.0081** |
| Right G_temp_sup-G_T_transv (107) | 12.65 + 1.16 (6.34) | 12.04 + 1.19 (6.53) | 10.76 + 1.22 (6.68) | 8.90 + 1.15 (6.33) | 5.24 + 1.03 (5.62) | 0.0079** |
| Right Lat_Fis-ant-Vertical (114) | 9.08 + 1.05 (5.75) | 9.67 + 1.08 (5.91) | 8.60 + 1.04 (5.68) | 5.70 + 0.95 (5.21) | 3.36 + 0.87 (4.78) | 0.0042** |
| Right S_circular_insula_sup (123) | 15.02 + 0.98 (5.36) | 15.67 + 1.00 (5.50) | 14.40 + 1.12 (6.14) | 10.91 + 1.22 (6.68) | 7.66 + 1.16 (6.37) | < 0.001** |
| Right S_temporal_transverse (148) | 12.21 + 1.26 (6.88) | 12.20 + 1.25 (6.85) | 10.39 + 1.18 (6.47) | 7.92 + 1.08 (5.94) | 5.09 + 1.02 (5.59) | 0.0059** |

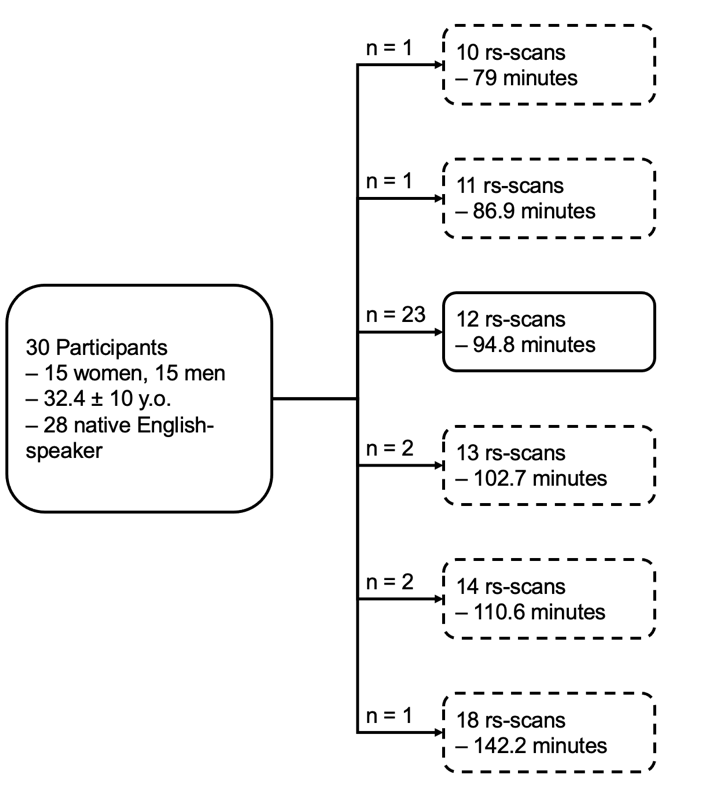

**Figure S1.** Flowchart of participant scanning organization. The originally designed protocol included 12 resting-state scans (indicated by the solid box).

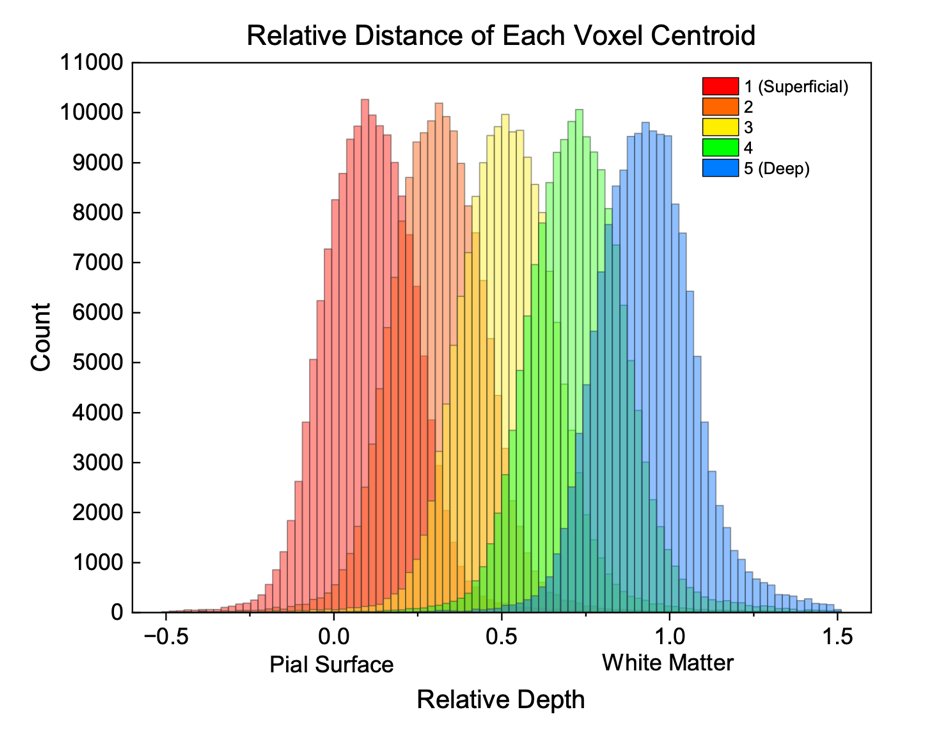

**Figure S2.** Relative distance of each voxel centroid organized by layer. The distance from each voxel centroid (within cortical volume) to the cortical surface (white matter/pial surface) was calculated. The relative distance was defined so that the depth at pial surface was zero, and one at the white matter border. Next, voxels intersecting each layer were picked and plotted with respect to their relative distances in a histogram.

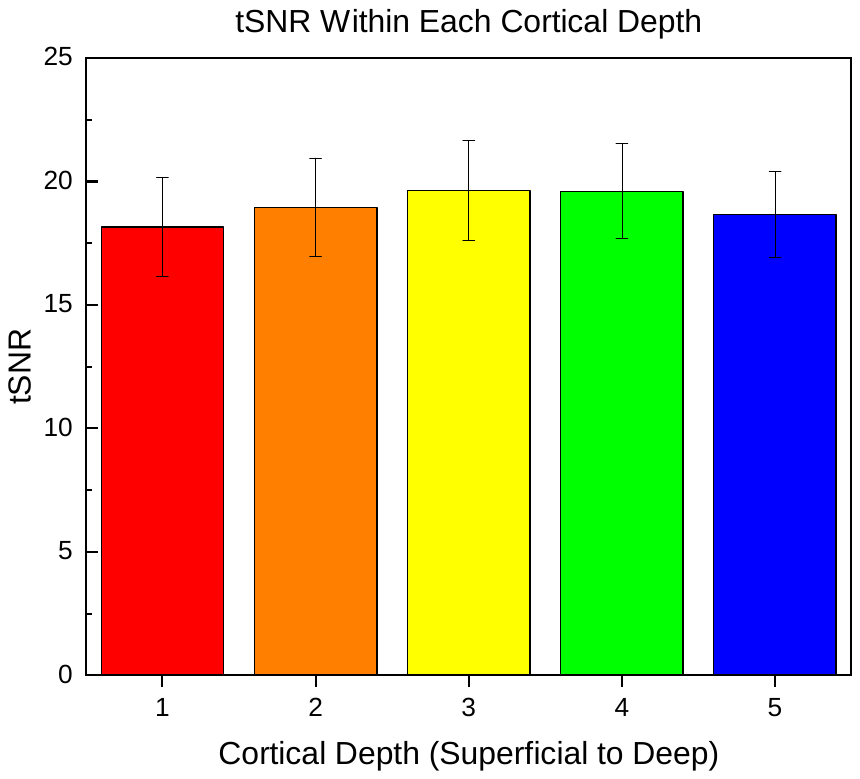

**Figure S3.** Temporal Signal-to-Noise Ratio (tSNR) across different cortical depths. Error bars indicate standard deviation.

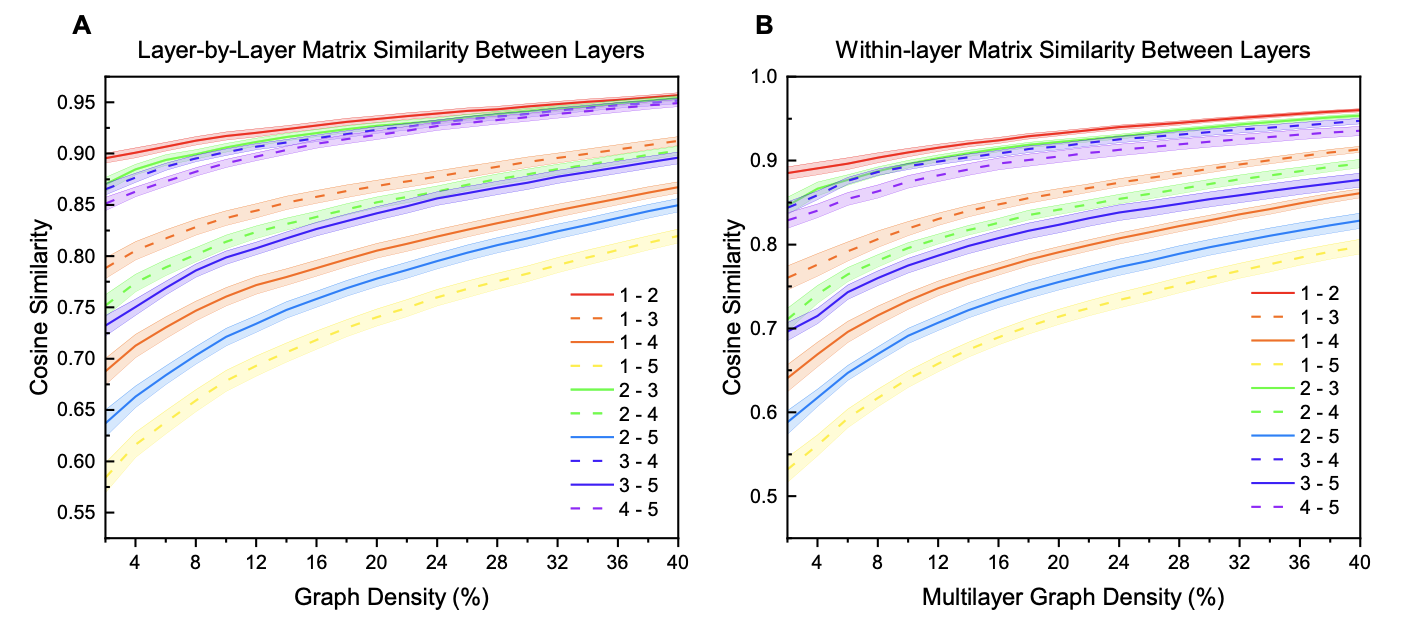

**Figure S4. (A)** Cosine similarity between each layer using layer-by-layer connectivity matrices. Within participant, each layer’s matrix was compared using cosine similarity across a range of thresholds. Cosine similarity values range from -1 (maximal dissimilarity) to +1 (maximal similarity). The mean value at each threshold is plotted while the shaded region indicates the standard error. **(B)** Cosine similarity between each layer using within-layer matrices. With participant, each layer’s matrix was compared using cosine similarity across a range of thresholds. The mean value at each threshold is plotted while the shaded region indicates the standard error.

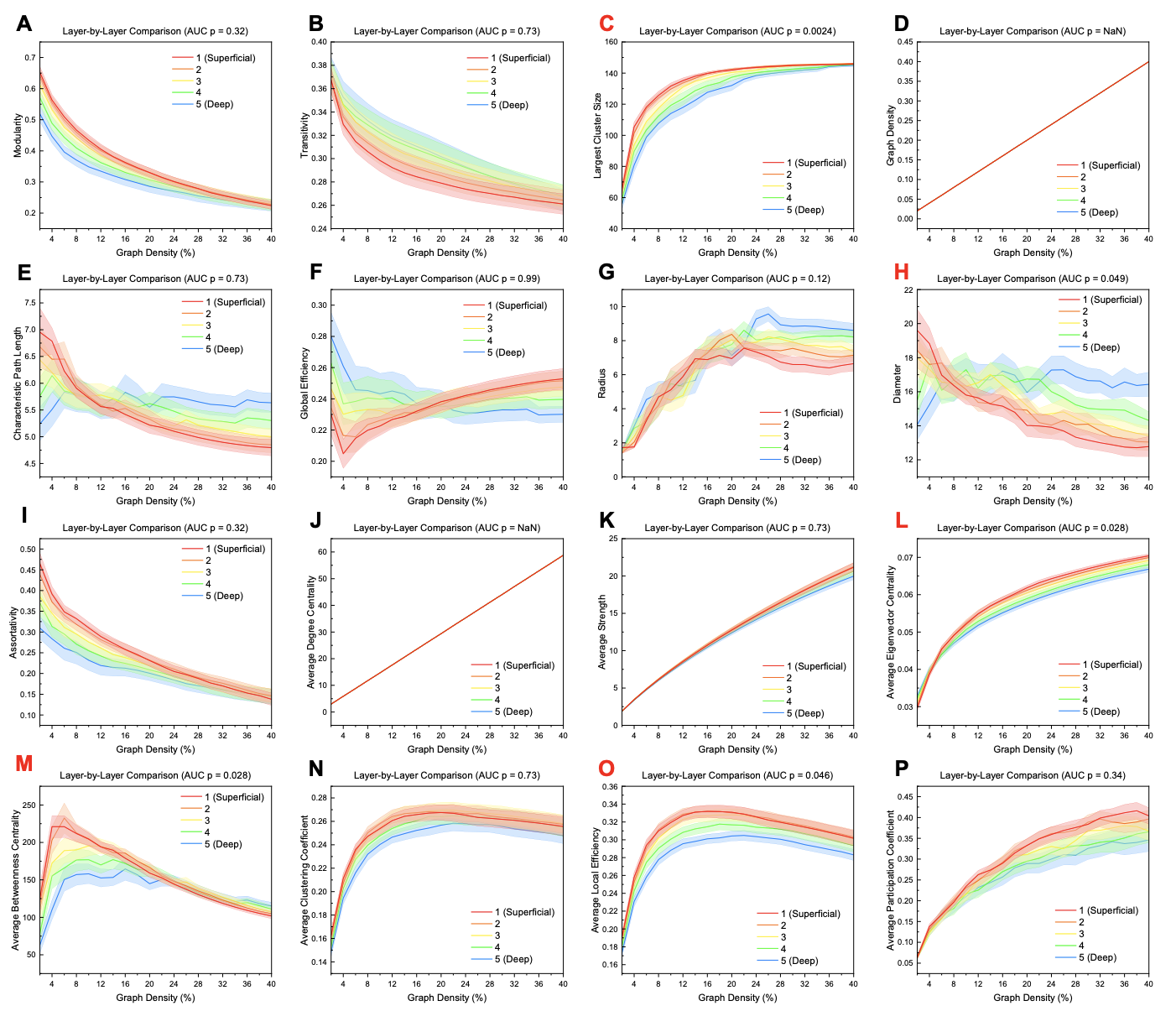

**Figure S5.** Global measures across different thresholds for layer-by-layer analysis. Significance for area-under-the-curve (AUC) values was calculated using a one-way ANOVA with an FDR correction (alpha = 0.05). Graph density and average degree centrality were constant across layers due to both s being a direct function of thresholding. Red letters indicate a measure that is significantly different between layers.

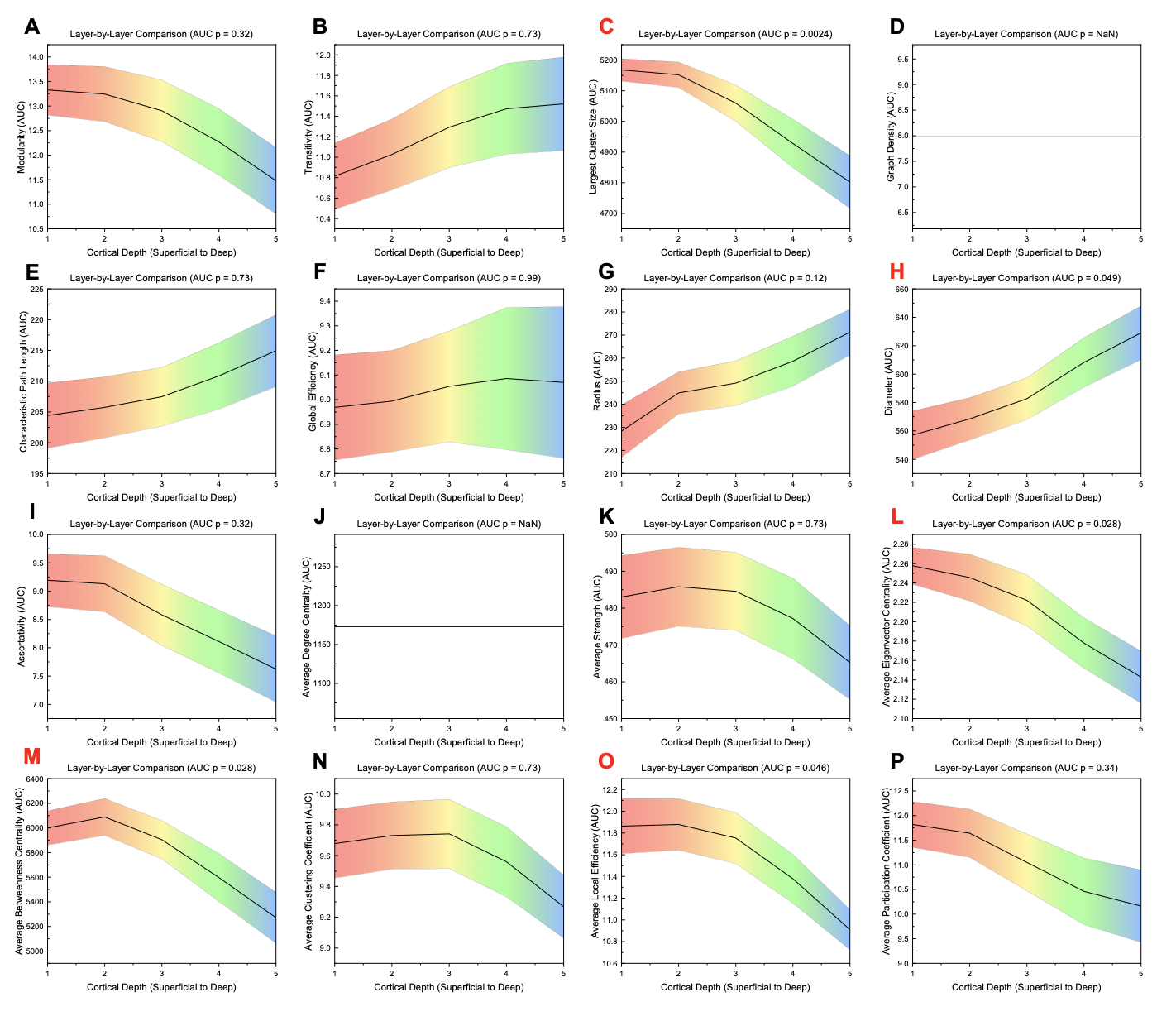

**Figure S6.** Area-under-the-curve (AUC) values across different layers for global measures for layer-by-layer analysis. Significance was calculated using a one-way ANOVA with an FDR correction (alpha = 0.05). Graph density and average degree centrality were constant across layers due to both s being a direct function of thresholding. Red letters indicate a measure that is significantly different between layers.

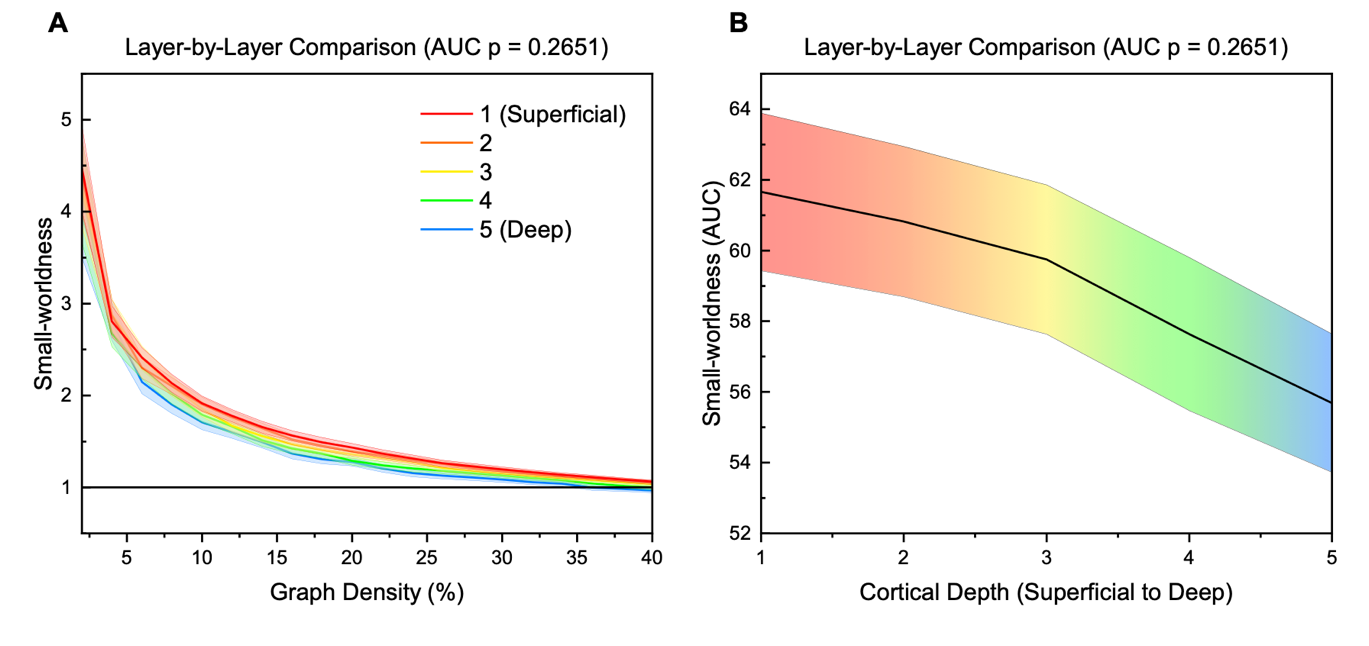

**Figure S7.** **(A)** Small-worldness across different thresholds for layer-by-layer analysis. The horizontal black line indicates the small-world threshold of one. **(B)** Area-under-the-curve (AUC) values across different layers for small-worldness for layer-by-layer analysis. Significance for area-under-the-curve (AUC) values was calculated using a one-way ANOVA. The mean value across participants at each layer is plotted while the shaded region indicates the standard error.

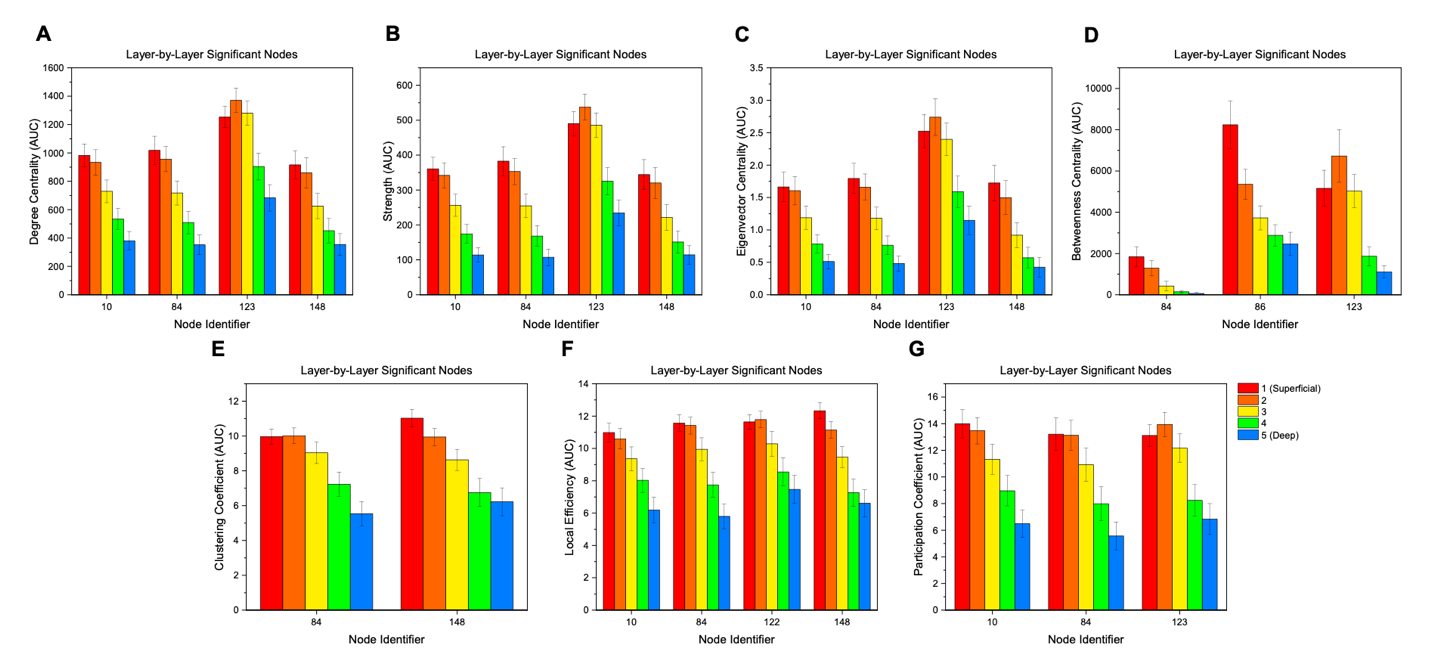

**Figure S8.** Area-under-the-curve (AUC) values for each significant nodal measure for layer-by-layer analysis: **(A)** degree centrality, **(B)** strength, **(C)** eigenvector centrality, **(D)** betweenness centrality, **(E)** clustering coefficient, **(F)** local efficiency, and **(G)** participation coefficient. Significance was calculated using a one-way ANOVA with an FDR correction (alpha = 0.01) to account for multiple comparisons (Groppe, 2023; Holm, 1979). Error bars display standard error. Node identifier numbers can be found in Table S1.

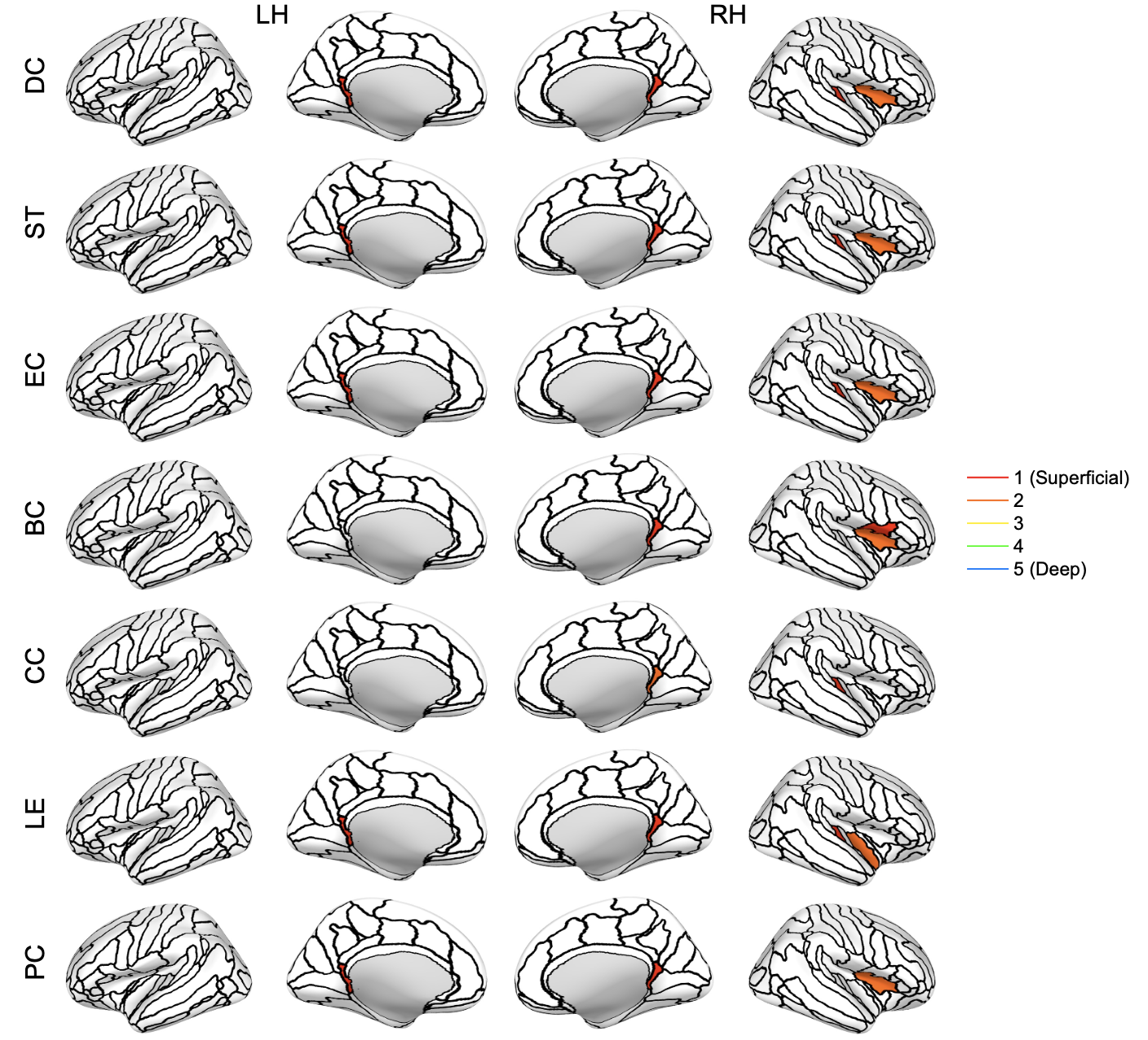

**Figure S9.** Nodes with significant differences between layers for the layer-by-layer analysis (inflated cortical surface for the corresponding hemisphere). Significance was calculated from the area-under-the curve (AUC) values using a one-way ANOVA with an FDR correction (alpha = 0.01) to account for multiple comparisons (Groppe, 2023; Holm, 1979). The colored section represents the layer with the highest value for the node. The nodes are based on the Destrieux atlas in FreeSurfer (Destrieux et al., 2010; Fischl et al., 2004). LH: left hemisphere; RH: right hemisphere; DC: degree centrality; ST: strength; EC: eigenvector centrality; BC: betweenness centrality; CC: clustering coefficient; LE: local efficiency; PC: participation coefficient.

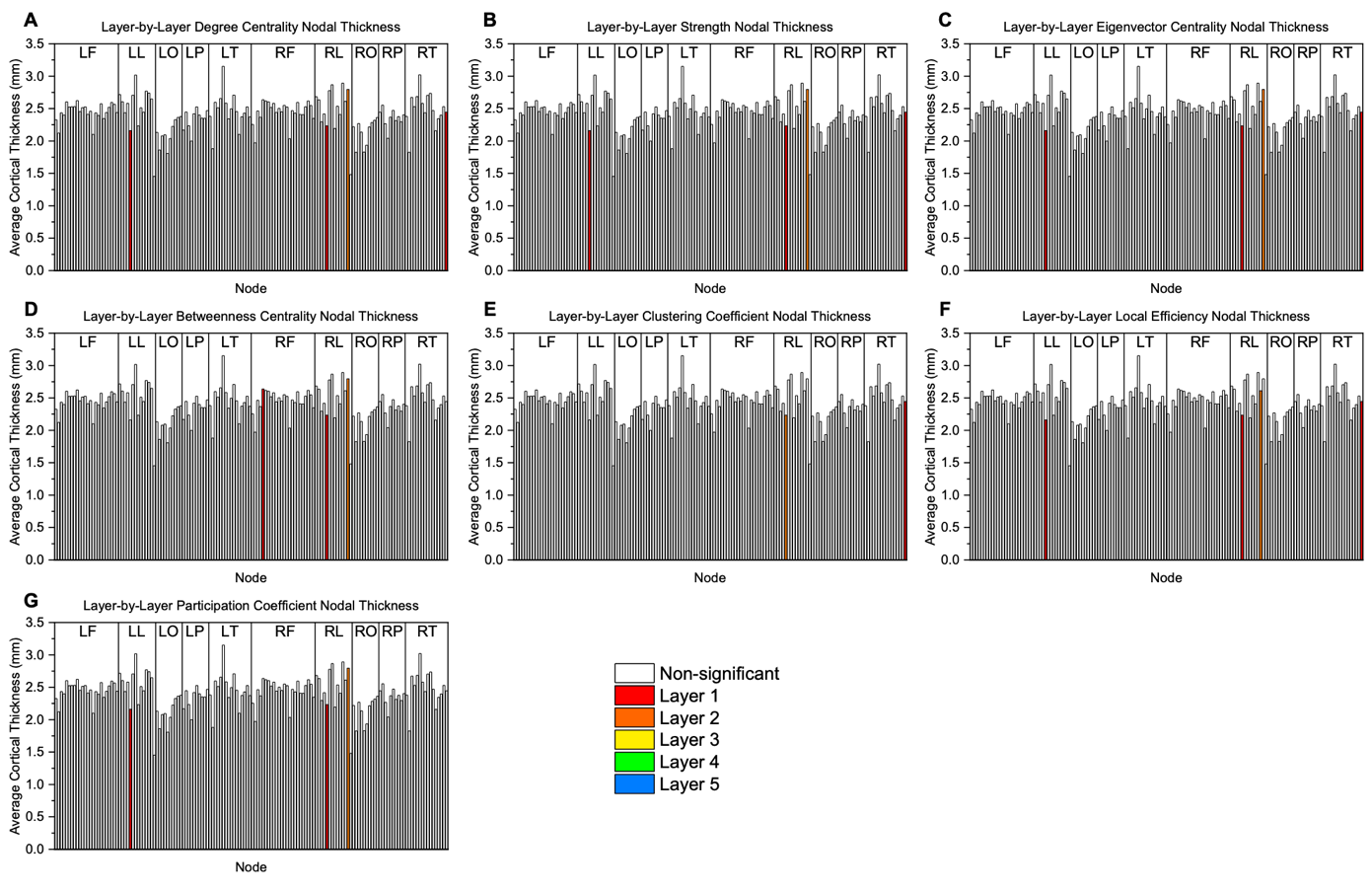

**Figure S10.** Thickness values grouped by brain region for the layer-by-layer analysis: **(A)** degree centrality, **(B)** strength, **(C)** eigenvector centrality, **(D)** betweenness centrality, **(E)** clustering coefficient, **(F)** local efficiency, and **(G)** participation coefficient. The colored bar represents a significant node and the layer with the highest value for the node. Significance was calculated using a one-way ANOVA with an FDR correction (alpha = 0.01) to account for multiple comparisons (Groppe, 2023; Holm, 1979). LF: left frontal; LL: left limbic; LO: left occipital; LP: left parietal; LT: left temporal; RF: right frontal; RL: right limbic; RO: right occipital; RP: right parietal; RT: right temporal.

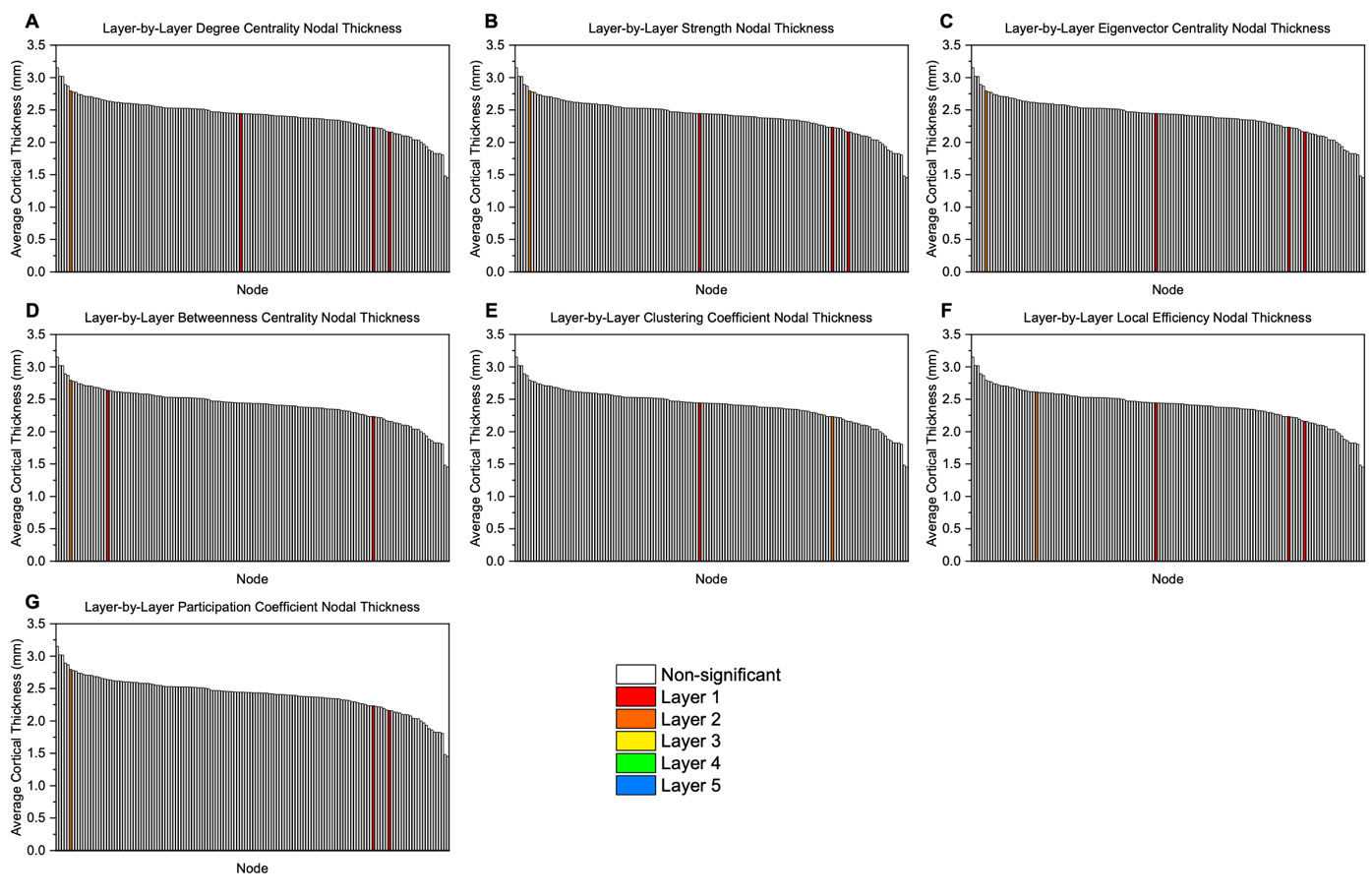

**Figure S11.** Thickness values sorted from largest to smallest value for the layer-by-layer analysis: **(A)** degree centrality, **(B)** strength, **(C)** eigenvector centrality, **(D)** betweenness centrality, **(E)** clustering coefficient, **(F)** local efficiency, and **(G)** participation coefficient. The colored bar represents a significant node and the layer with the highest value for the node. Significance was calculated using a one-way ANOVA with an FDR correction (alpha = 0.01) to account for multiple comparisons (Groppe, 2023; Holm, 1979). LF: left frontal; LL: left limbic; LO: left occipital; LP: left parietal; LT: left temporal; RF: right frontal; RL: right limbic; RO: right occipital; RP: right parietal; RT: right temporal.

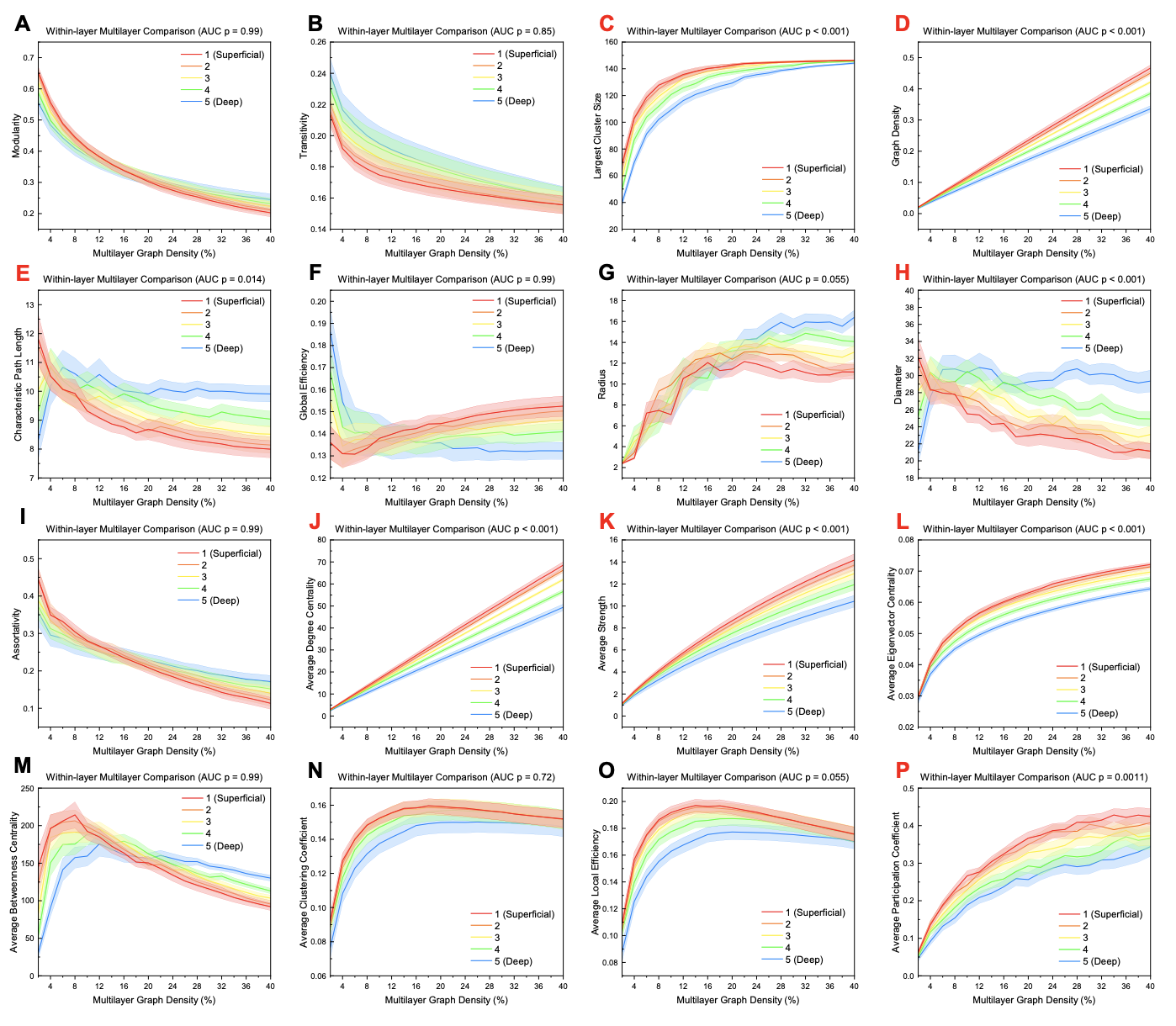

**Figure S12.** Global measures across different thresholds for within-layer analysis. Significance for area-under-the-curve (AUC) values was calculated using a one-way ANOVA with an FDR correction (alpha = 0.05). Red letters indicate a measure that is significantly different between layers.

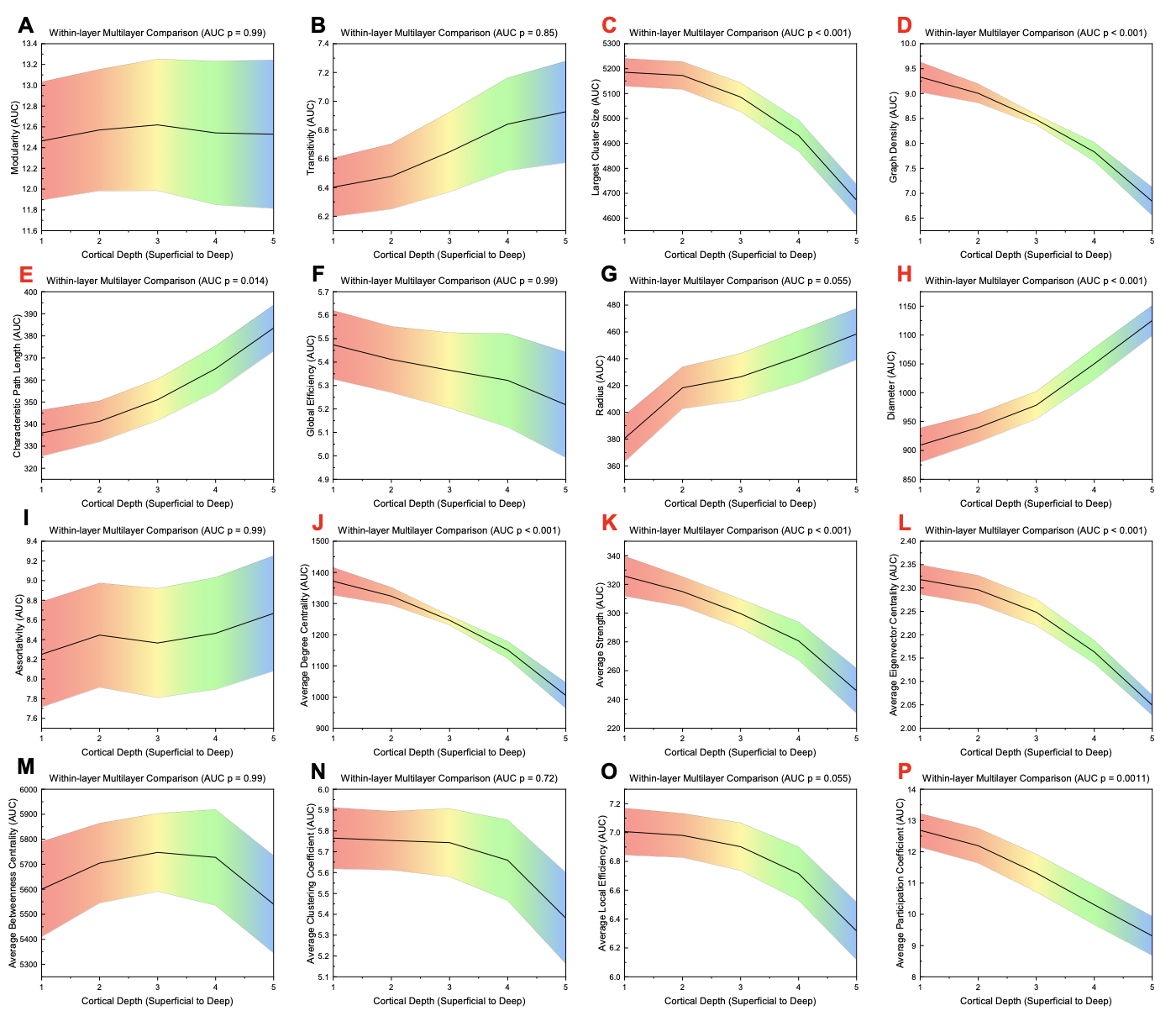

**Figure S13.** Area-under-the-curve (AUC) values across different layers for global measures for within-layer analysis. Significance was calculated using a one-way ANOVA with an FDR correction (alpha = 0.05). Red letters indicate a measure that is significantly different between layers.

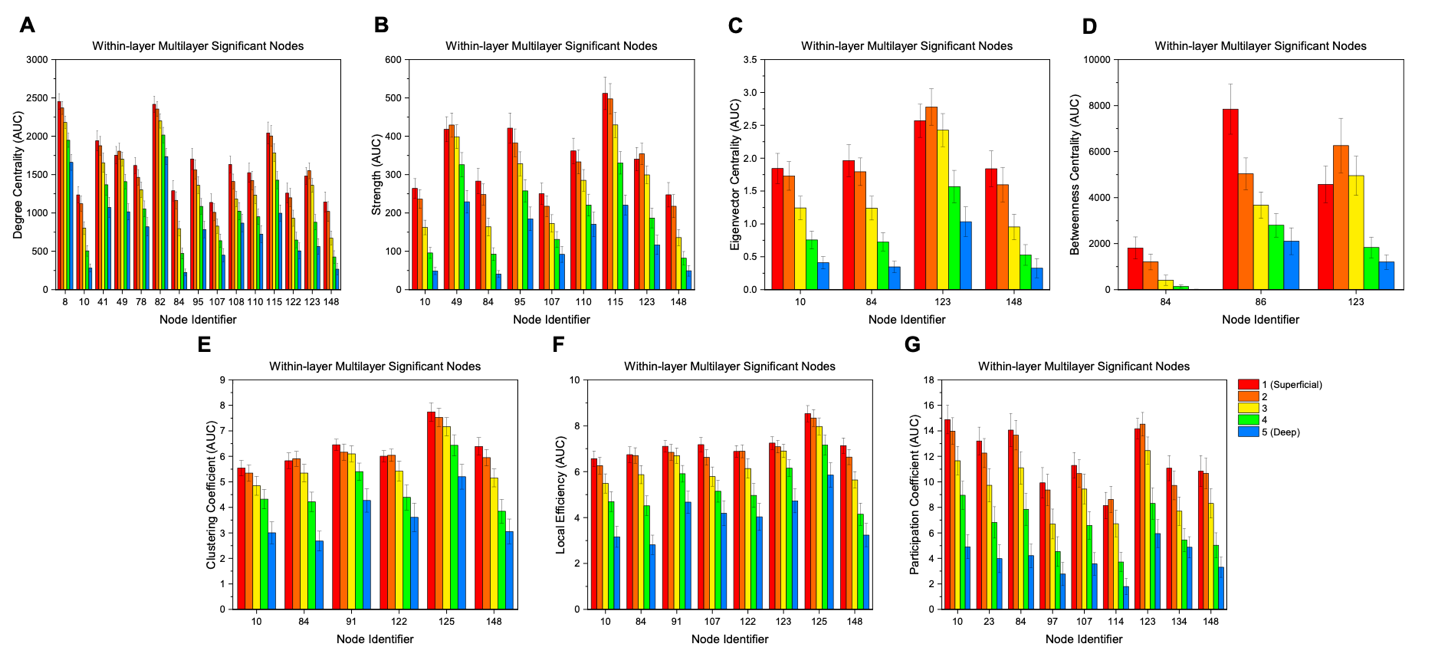

**Figure S14.** Area-under-the-curve (AUC) values for each significant nodal measure for within-layer analysis: **(A)** degree centrality, **(B)** strength, **(C)** eigenvector centrality, **(D)** betweenness centrality, **(E)** clustering coefficient, **(F)** local efficiency, and **(G)** participation coefficient. Significance was calculated using a one-way ANOVA with an FDR correction (alpha = 0.01) to account for multiple comparisons (Groppe, 2023; Holm, 1979). Error bars display standard error. Node identifier numbers can be found in Table S1.

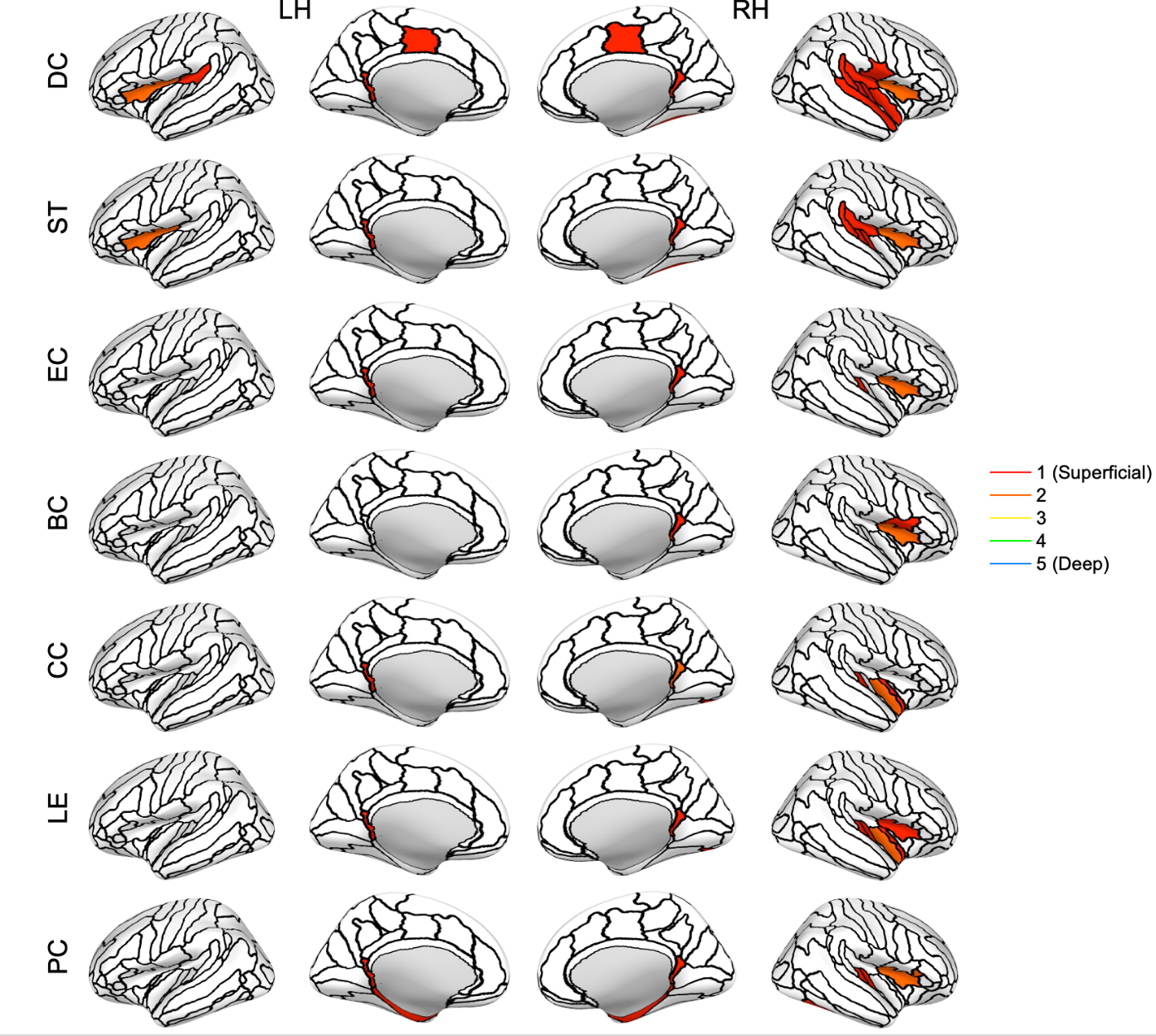

**Figure S15.** Nodes with significant differences between layers for the within-layer analysis (inflated cortical surface for the corresponding hemisphere). Significance was calculated from the area-under-the curve (AUC) values using a one-way ANOVA with an FDR correction (alpha = 0.01) to account for multiple comparisons (Groppe, 2023; Holm, 1979). The colored section represents the layer with the highest value for the node. The nodes are based on the Destrieux atlas in FreeSurfer (Destrieux et al., 2010; Fischl et al., 2004). LH: left hemisphere; RH: right hemisphere; DC: degree centrality; ST: strength; EC: eigenvector centrality; BC: betweenness centrality; CC: clustering coefficient; LE: local efficiency; PC: participation coefficient.

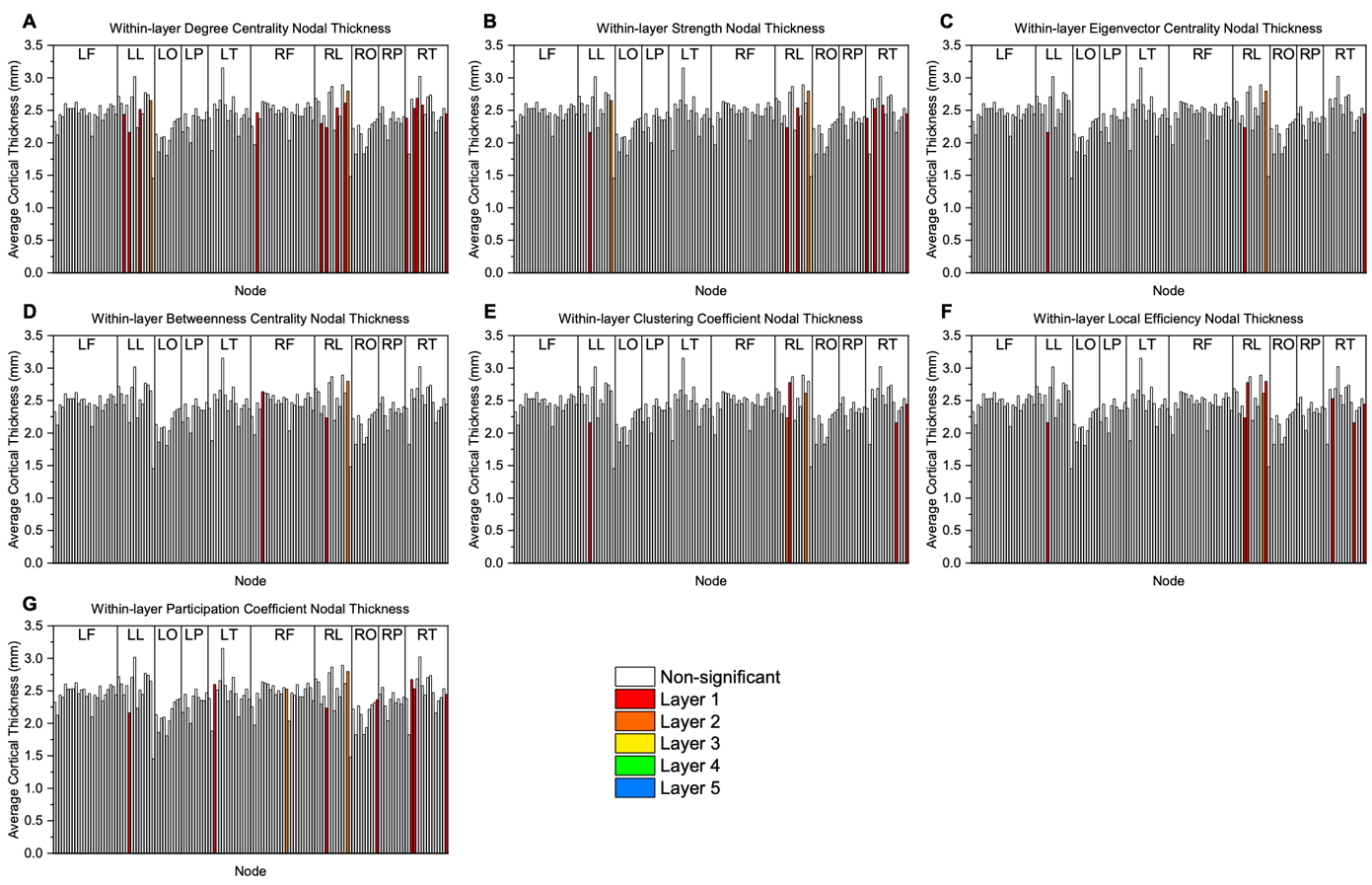

**Figure S16.** Thickness values grouped by brain region for the within-layer analysis: **(A)** degree centrality, **(B)** strength, **(C)** eigenvector centrality, **(D)** betweenness centrality, **(E)** clustering coefficient, **(F)** local efficiency, and **(G)** participation coefficient. The colored bar represents a significant node and the layer with the highest value for the node. Significance was calculated using a one-way ANOVA with an FDR correction (alpha = 0.01) to account for multiple comparisons (Groppe, 2023; Holm, 1979). LF: left frontal; LL: left limbic; LO: left occipital; LP: left parietal; LT: left temporal; RF: right frontal; RL: right limbic; RO: right occipital; RP: right parietal; RT: right temporal.

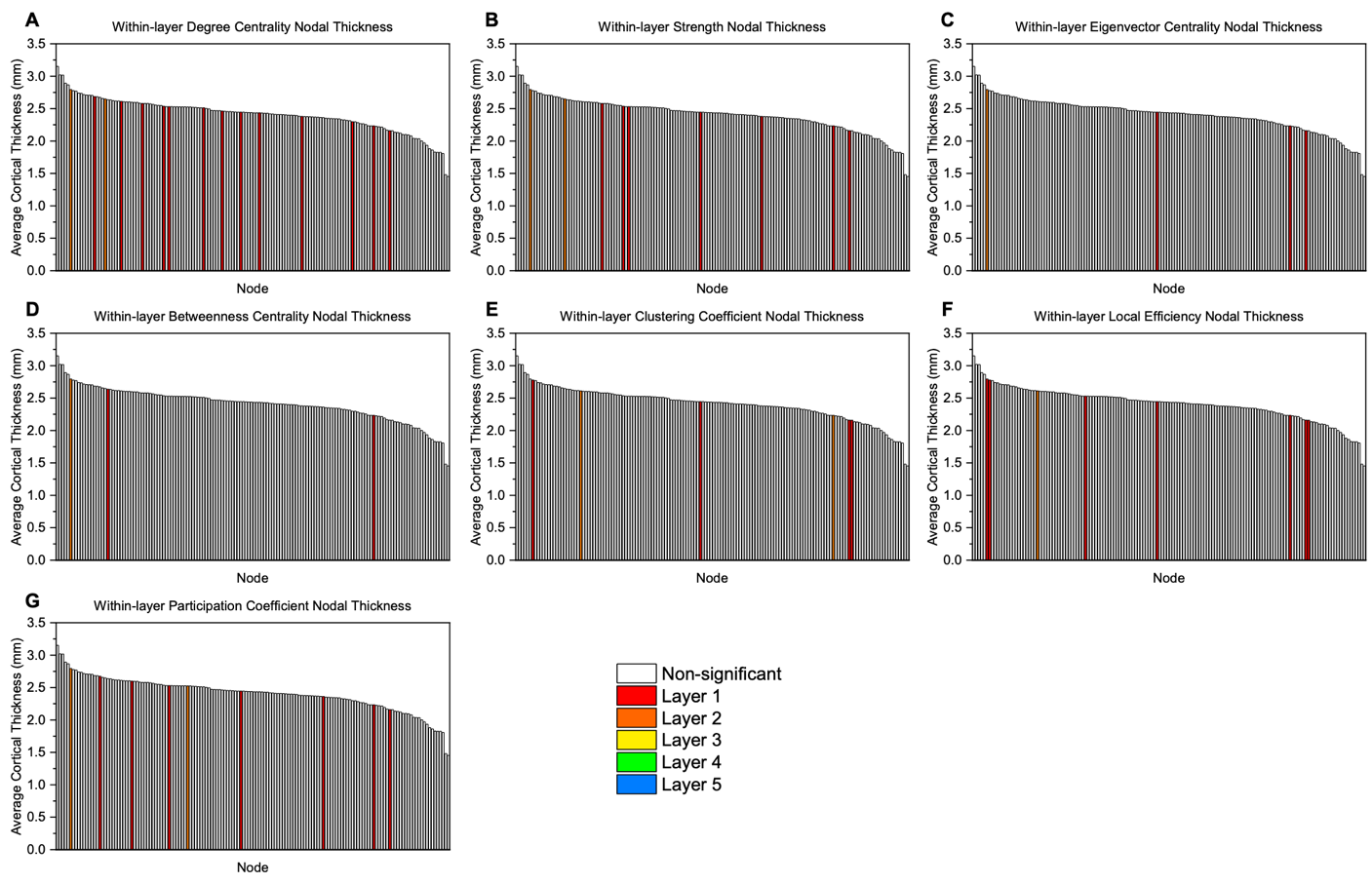

**Figure S17.** Thickness values sorted from largest to smallest value for the within-layer analysis: **(A)** degree centrality, **(B)** strength, **(C)** eigenvector centrality, **(D)** betweenness centrality, **(E)** clustering coefficient, **(F)** local efficiency, and **(G)** participation coefficient. The colored bar represents a significant node and the layer with the highest value for the node. Significance was calculated using a one-way ANOVA with an FDR correction (alpha = 0.01) to account for multiple comparisons (Groppe, 2023; Holm, 1979). LF: left frontal; LL: left limbic; LO: left occipital; LP: left parietal; LT: left temporal; RF: right frontal; RL: right limbic; RO: right occipital; RP: right parietal; RT: right temporal.

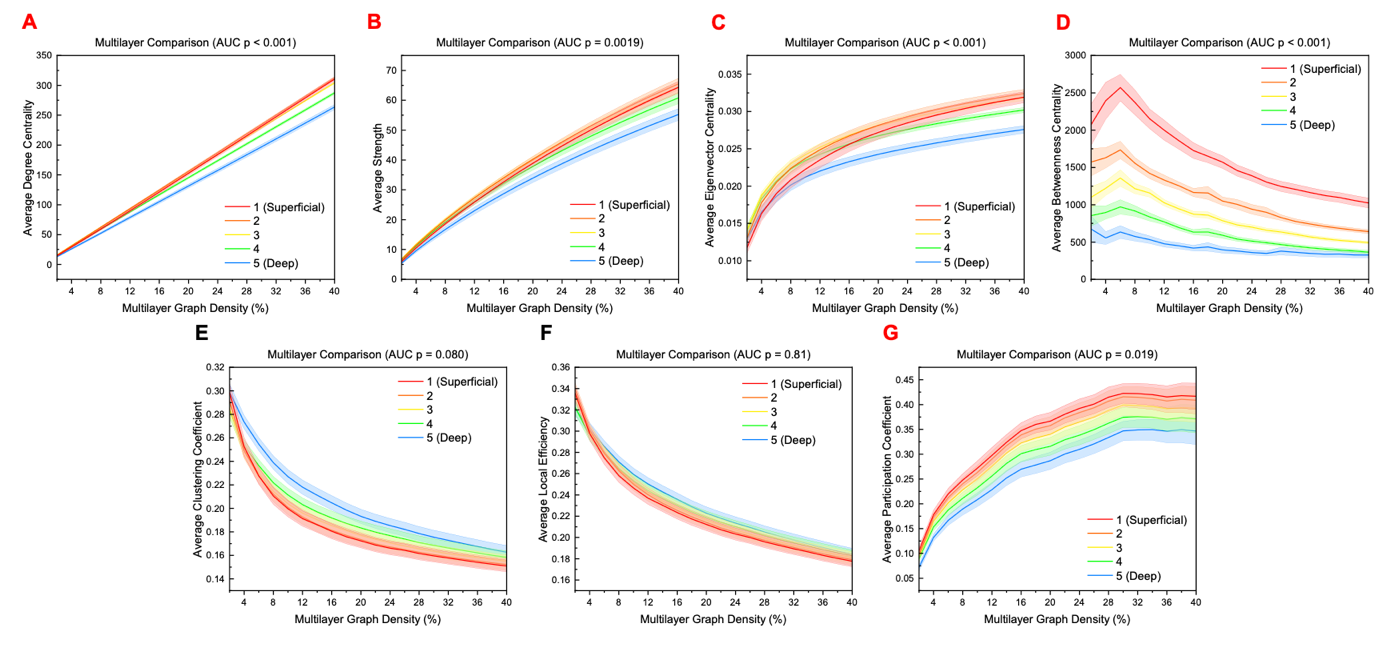

**Figure S18.** Global measures across different thresholds for multilayer analysis. Significance for area-under-the-curve (AUC) values was calculated using a one-way ANOVA with an FDR correction (alpha = 0.05). Only node-averaged global measures are shown since layer-specific global values cannot be extracted from supra-adjacency matrices. Red letters indicate a measure that is significantly different between layers.

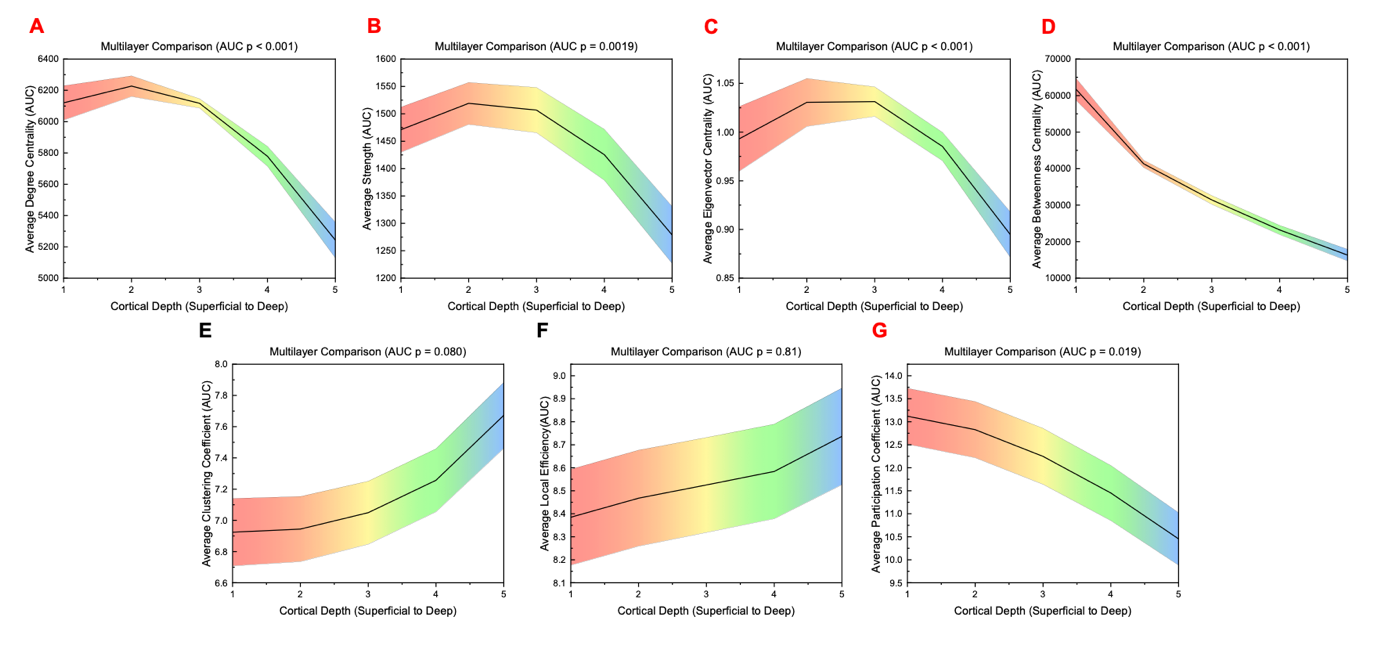

**Figure S19.** Area-under-the-curve (AUC) values across different layers for global measures for multilayer analysis. Significance was calculated using a one-way ANOVA with an FDR correction (alpha = 0.05). Only node-averaged global measures are shown since layer-specific global values cannot be extracted from supra-adjacency matrices. Red letters indicate a measure that is significantly different between layers.

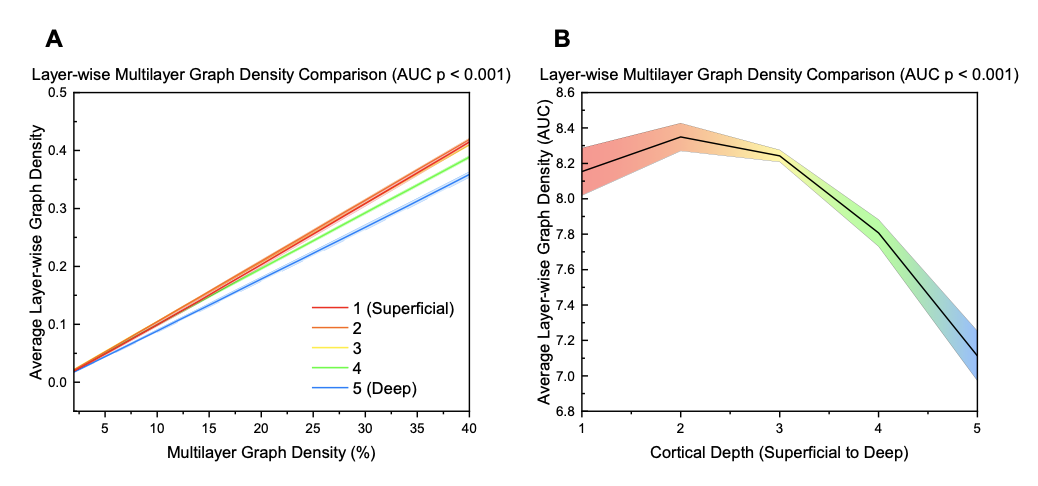

**Figure S20.** **(A)** Layer-wise graph density across different thresholds for multilayer analysis. **B)** Area-under-the-curve (AUC) values across different layers for layer-wise graph density for multilayer analysis. Significance for area-under-the-curve (AUC) values was calculated using a one-way ANOVA. The mean value across participants at each layer is plotted while the shaded region indicates the standard error.

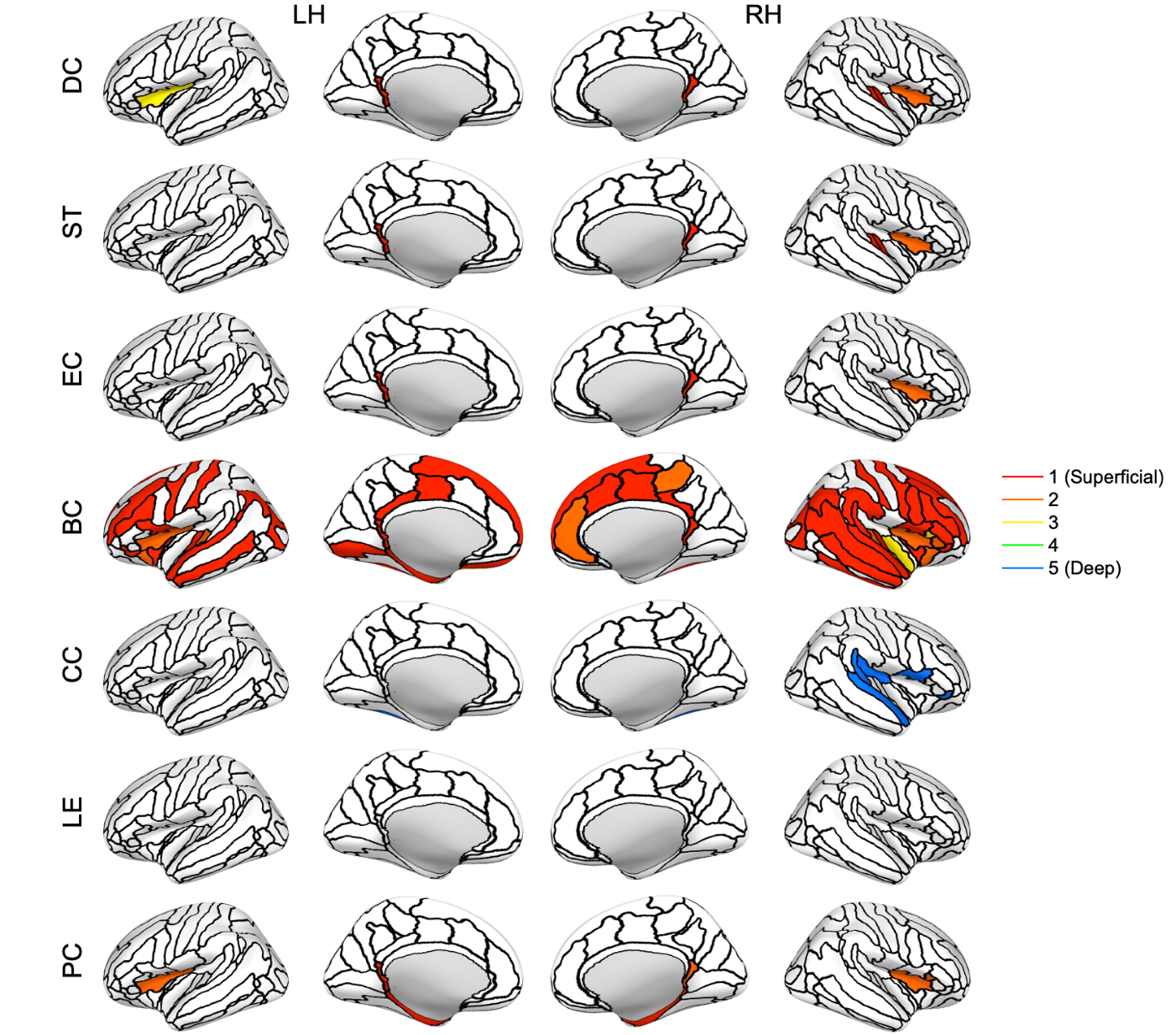

**Figure S21.** Nodes with significant differences between layers for the multilayer analysis (inflated cortical surface for the corresponding hemisphere). Significance was calculated from the area-under-the curve (AUC) values using a one-way ANOVA with an FDR correction (alpha = 0.01) to account for multiple comparisons (Groppe, 2023; Holm, 1979). The colored section represents the layer with the highest value for the node. The nodes are based on the Destrieux atlas in FreeSurfer (Destrieux et al., 2010; Fischl et al., 2004). LH: left hemisphere; RH: right hemisphere; DC: degree centrality; ST: strength; EC: eigenvector centrality; BC: betweenness centrality; CC: clustering coefficient; LE: local efficiency; PC: participation coefficient.

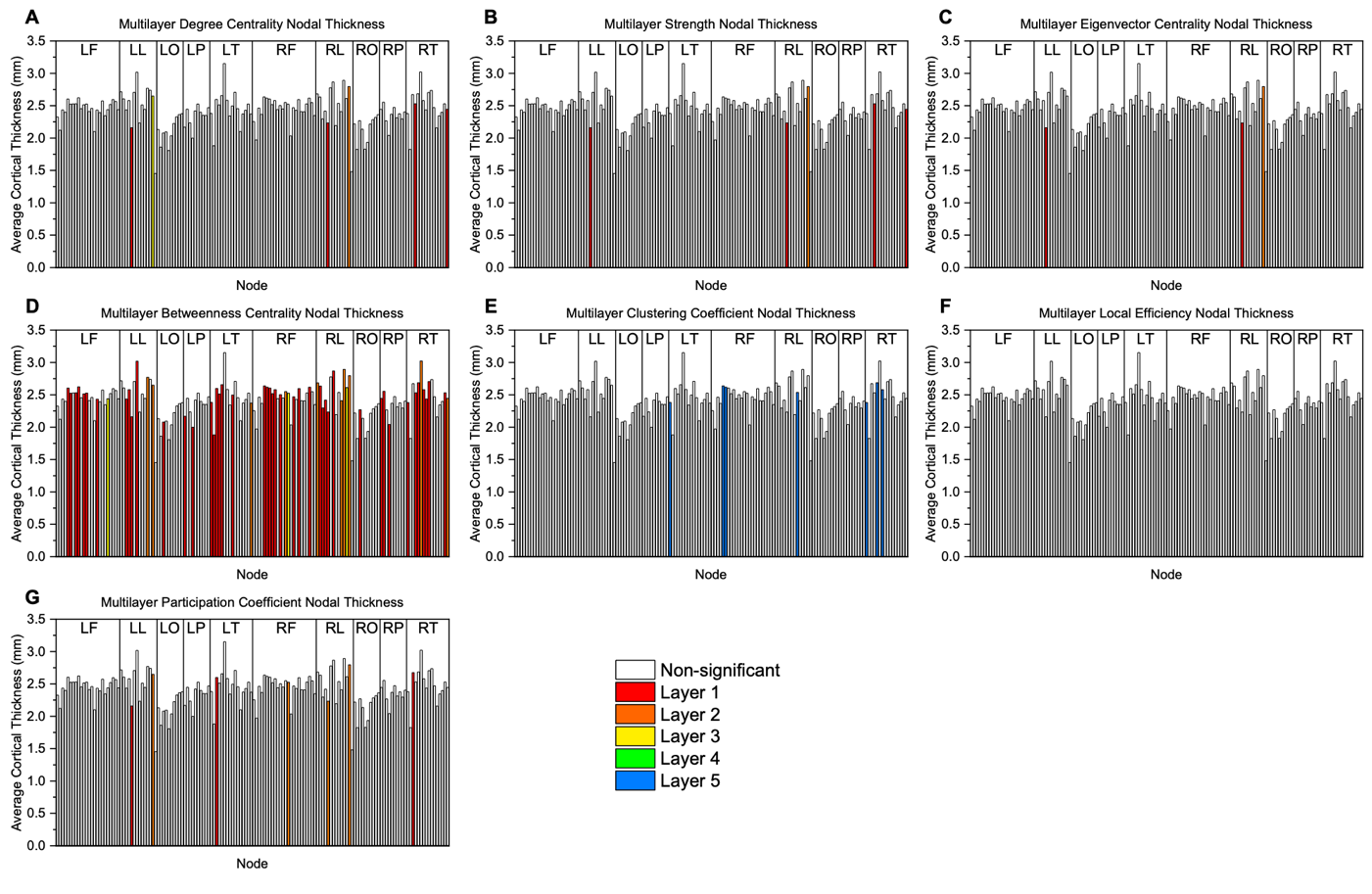

**Figure S22.** Thickness values grouped by brain region for the multilayer analysis: **(A)** degree centrality, **(B)** strength, **(C)** eigenvector centrality, **(D)** betweenness centrality, **(E)** clustering coefficient, **(F)** local efficiency, and **(G)** participation coefficient. The colored bar represents a significant node and the layer with the highest value for the node. Significance was calculated using a one-way ANOVA with an FDR correction (alpha = 0.01) to account for multiple comparisons (Groppe, 2023; Holm, 1979). LF: left frontal; LL: left limbic; LO: left occipital; LP: left parietal; LT: left temporal; RF: right frontal; RL: right limbic; RO: right occipital; RP: right parietal; RT: right temporal.

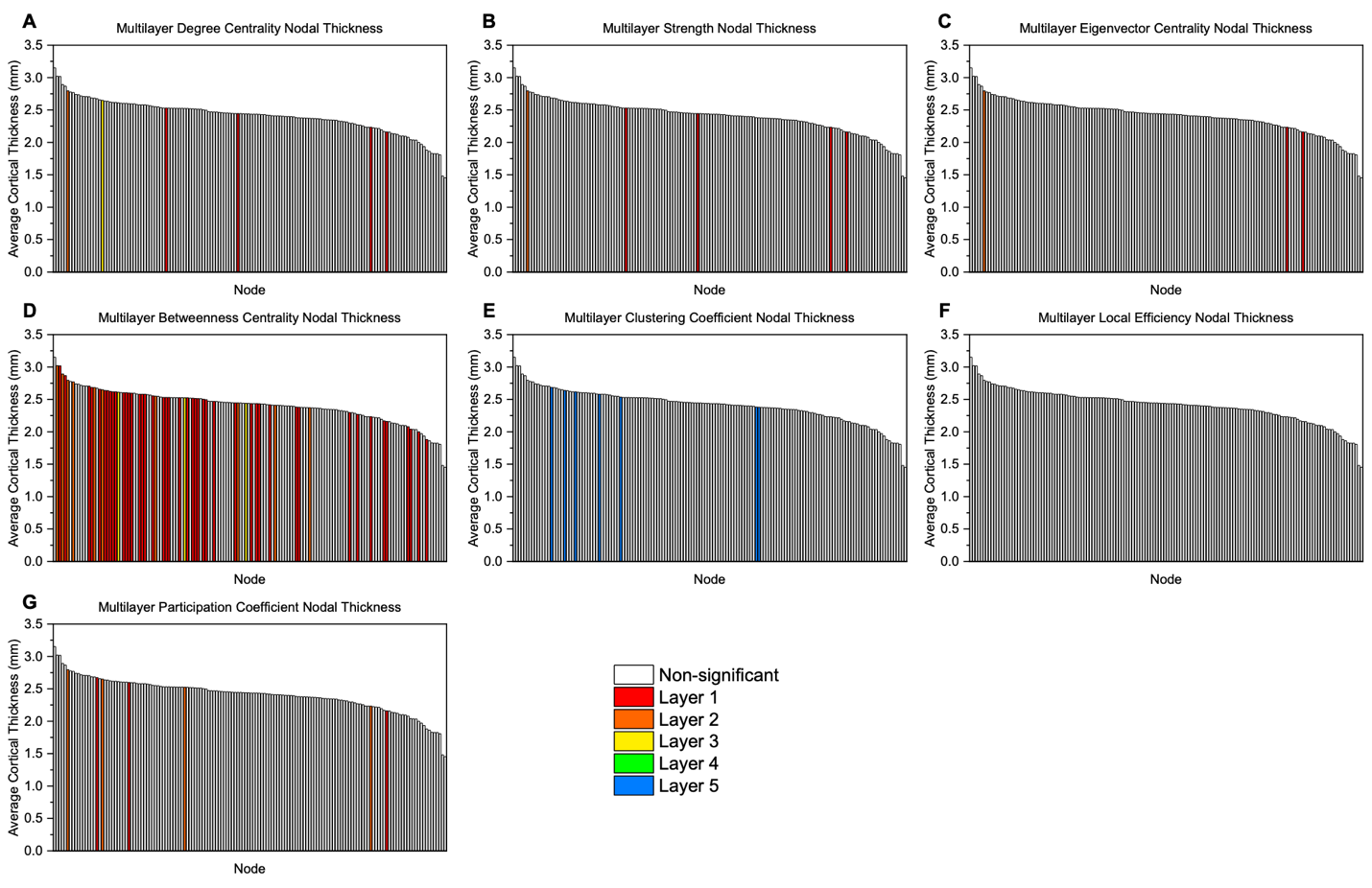

**Figure S23.** Thickness values sorted from largest to smallest value for the multilayer analysis: **(A)** degree centrality, **(B)** strength, **(C)** eigenvector centrality, **(D)** betweenness centrality, **(E)** clustering coefficient, **(F)** local efficiency, and **(G)** participation coefficient. The colored bar represents a significant node and the layer with the highest value for the node. Significance was calculated using a one-way ANOVA with an FDR correction (alpha = 0.01) to account for multiple comparisons (Groppe, 2023; Holm, 1979). LF: left frontal; LL: left limbic; LO: left occipital; LP: left parietal; LT: left temporal; RF: right frontal; RL: right limbic; RO: right occipital; RP: right parietal; RT: right temporal.

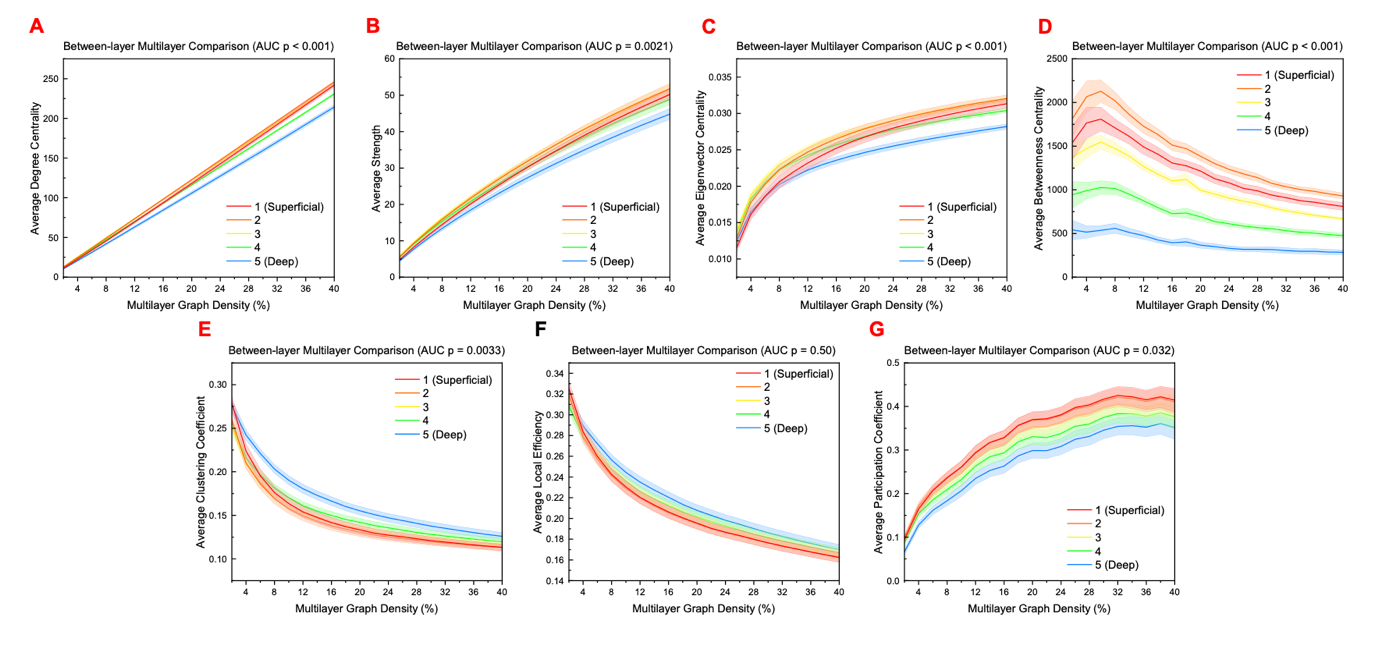

**Figure S24.** Global measures across different thresholds for between-layer analysis. Significance for area-under-the-curve (AUC) values was calculated using a one-way ANOVA with an FDR correction (alpha = 0.05). Only node-averaged global measures are shown since layer-specific global values cannot be extracted from supra-adjacency matrices. Red letters indicate a measure that is significantly different between layers.

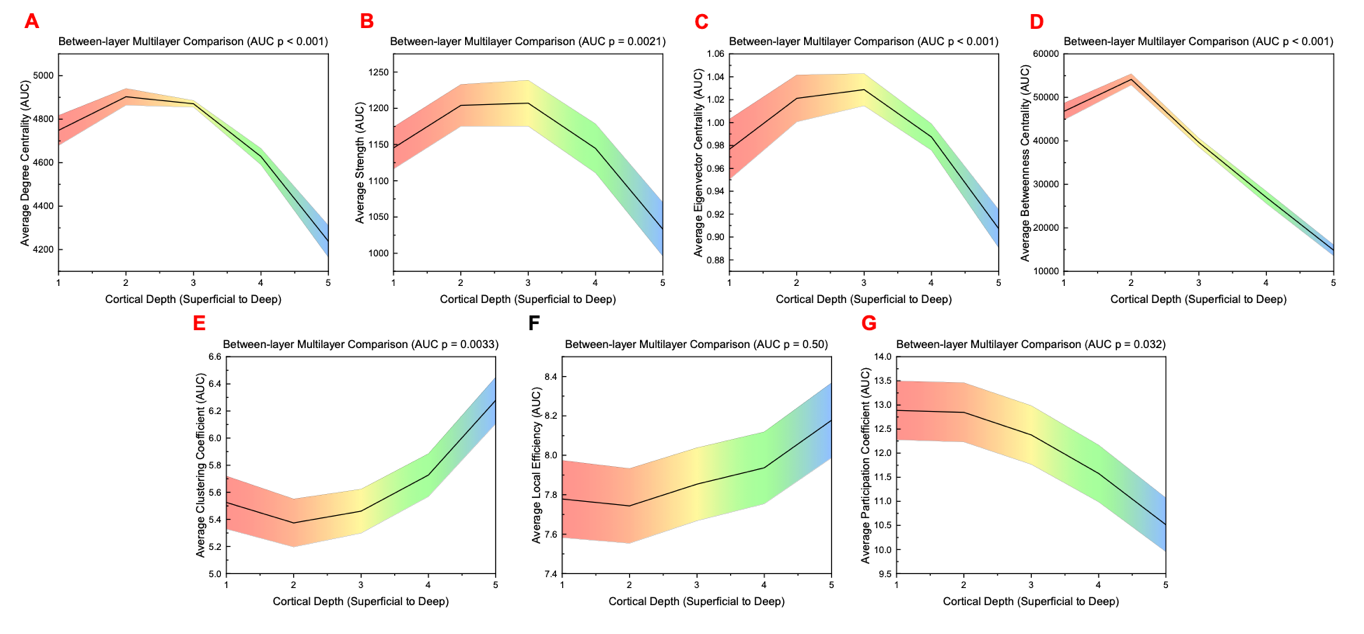

**Figure S25.** Area-under-the-curve (AUC) values across different layers for global measures for between-layer analysis. Significance was calculated using a one-way ANOVA with an FDR correction (alpha = 0.05). Only node-averaged global measures are shown since layer-specific global values cannot be extracted from supra-adjacency matrices. Red letters indicate a measure that is significantly different between layers.

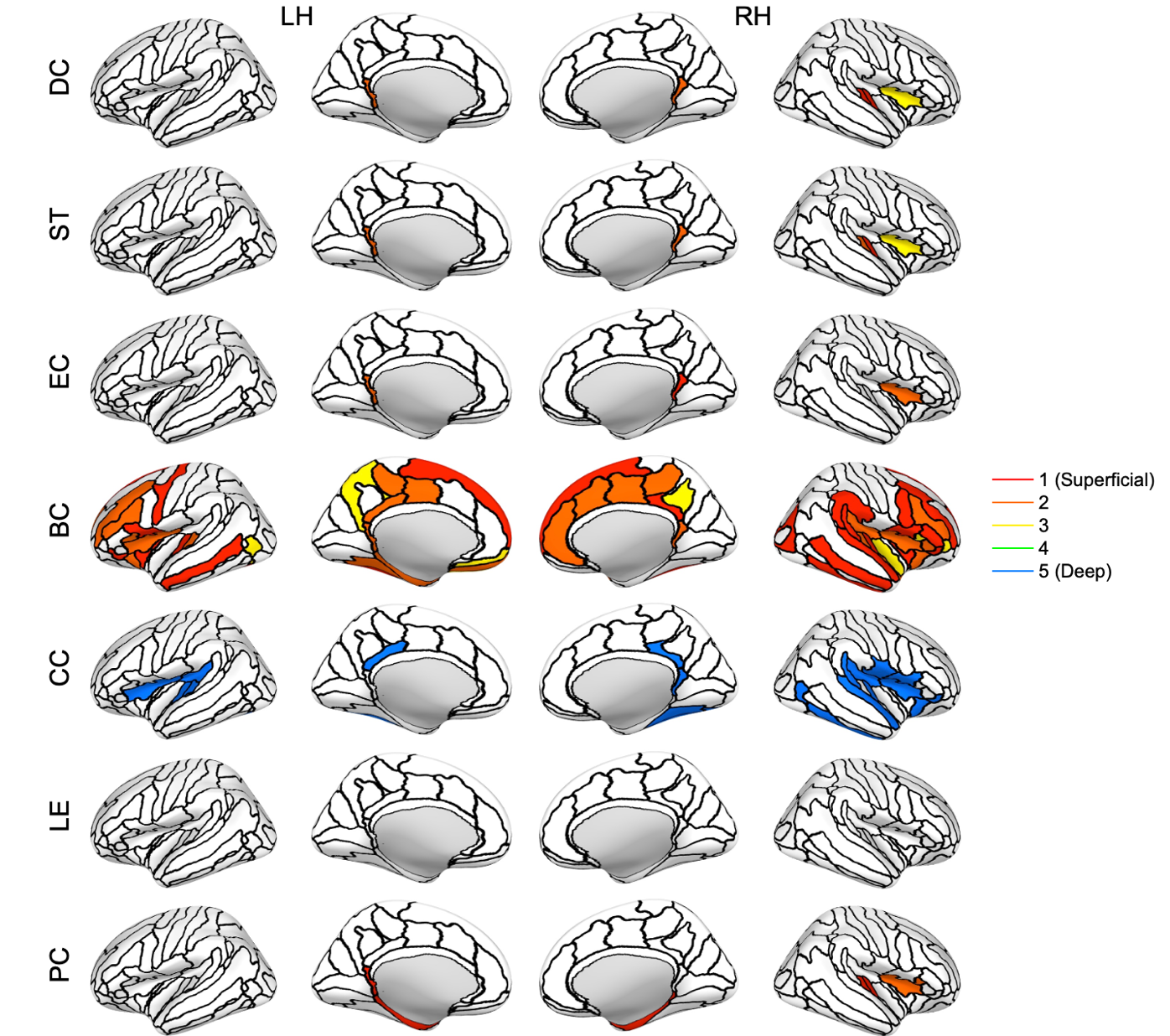

**Figure S26.** Nodes with significant differences between layers for the between-layer analysis (inflated cortical surface for the corresponding hemisphere). Significance was calculated from the area-under-the curve (AUC) values using a one-way ANOVA with an FDR correction (alpha = 0.01) to account for multiple comparisons (Groppe, 2023; Holm, 1979). The colored section represents the layer with the highest value for the node. The nodes are based on the Destrieux atlas in FreeSurfer (Destrieux et al., 2010; Fischl et al., 2004). LH: left hemisphere; RH: right hemisphere; DC: degree centrality; ST: strength; EC: eigenvector centrality; BC: betweenness centrality; CC: clustering coefficient; LE: local efficiency; PC: participation coefficient.

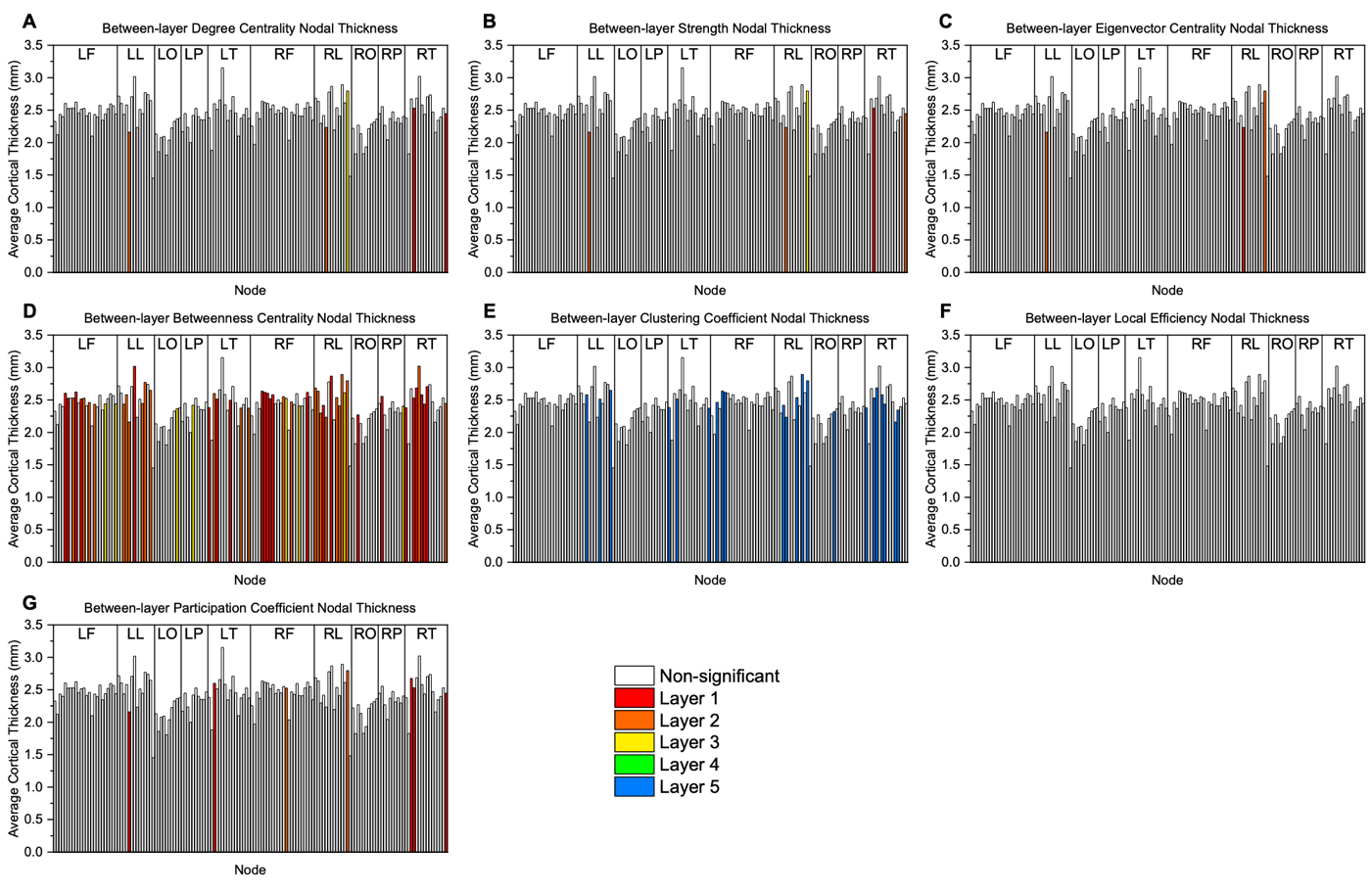

**Figure S27.** Thickness values grouped by brain region for the between-layer analysis: **(A)** degree centrality, **(B)** strength, **(C)** eigenvector centrality, **(D)** betweenness centrality, **(E)** clustering coefficient, **(F)** local efficiency, and **(G)** participation coefficient. The colored bar represents a significant node and the layer with the highest value for the node. Significance was calculated using a one-way ANOVA with an FDR correction (alpha = 0.01) to account for multiple comparisons (Groppe, 2023; Holm, 1979). LF: left frontal; LL: left limbic; LO: left occipital; LP: left parietal; LT: left temporal; RF: right frontal; RL: right limbic; RO: right occipital; RP: right parietal; RT: right temporal.

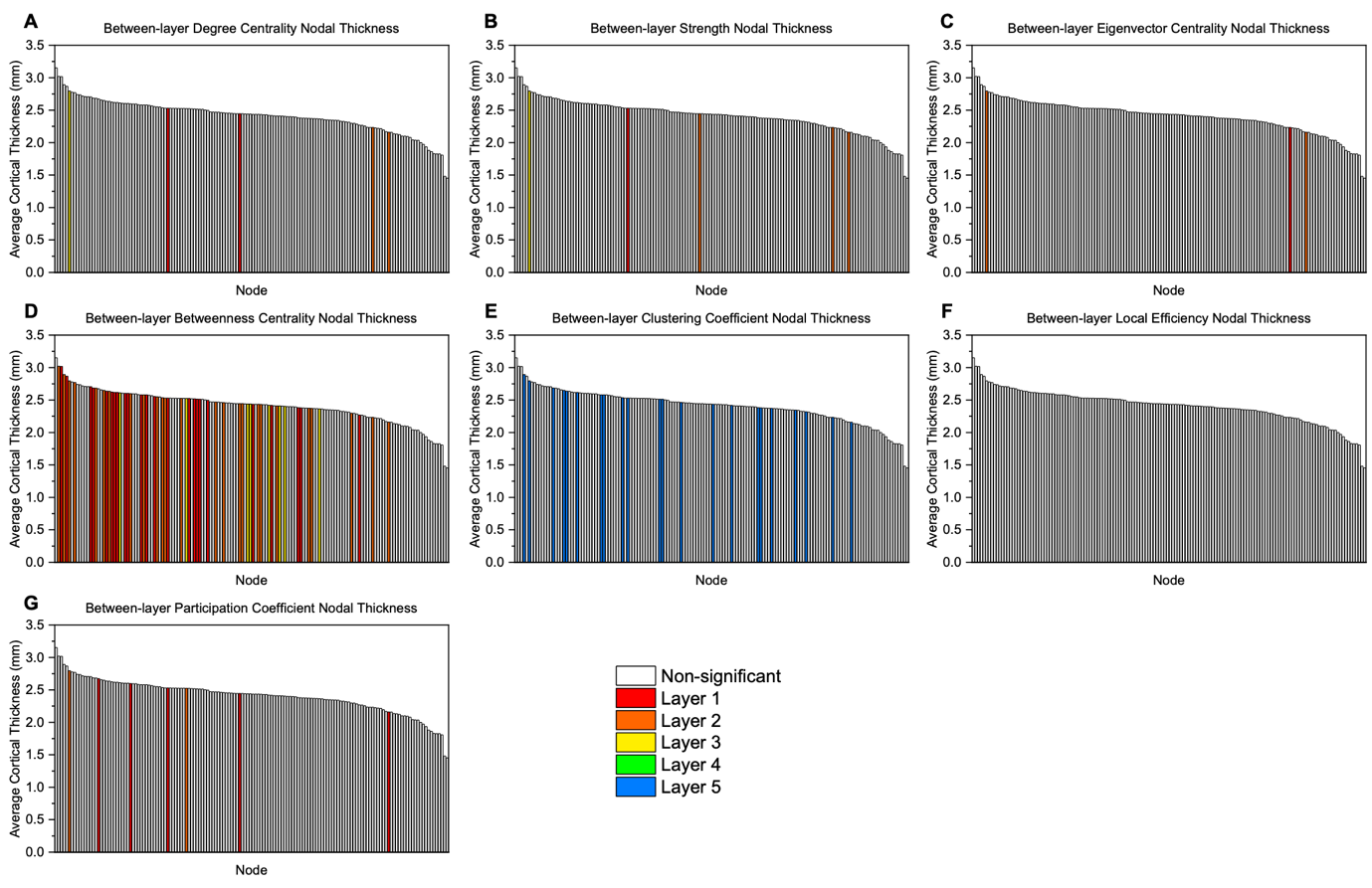

**Figure S28.** Thickness values sorted from largest to smallest value for the between-layer analysis: **(A)** degree centrality, **(B)** strength, **(C)** eigenvector centrality, **(D)** betweenness centrality, **(E)** clustering coefficient, **(F)** local efficiency, and **(G)** participation coefficient. The colored bar represents a significant node and the layer with the highest value for the node. Significance was calculated using a one-way ANOVA with an FDR correction (alpha = 0.01) to account for multiple comparisons (Groppe, 2023; Holm, 1979). LF: left frontal; LL: left limbic; LO: left occipital; LP: left parietal; LT: left temporal; RF: right frontal; RL: right limbic; RO: right occipital; RP: right parietal; RT: right temporal.

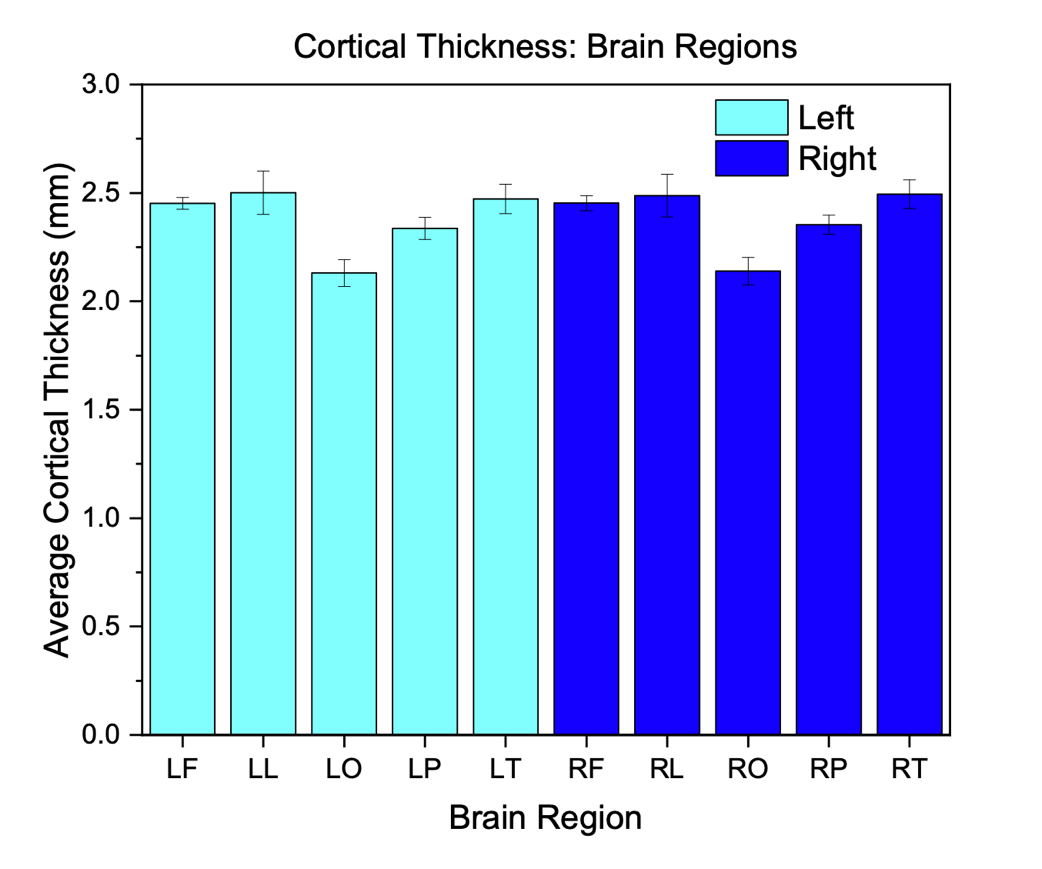

**Figure S29.** Average cortical thickness for each brain region. Error bars denote standard error. Thickness values are an average across all subjects and all nodes within each region (Table 1). LF: left frontal; LL: left limbic; LO: left occipital; LP: left parietal; LT: left temporal; RF: right frontal; RL: right limbic; RO: right occipital; RP: right parietal; RT: right temporal.

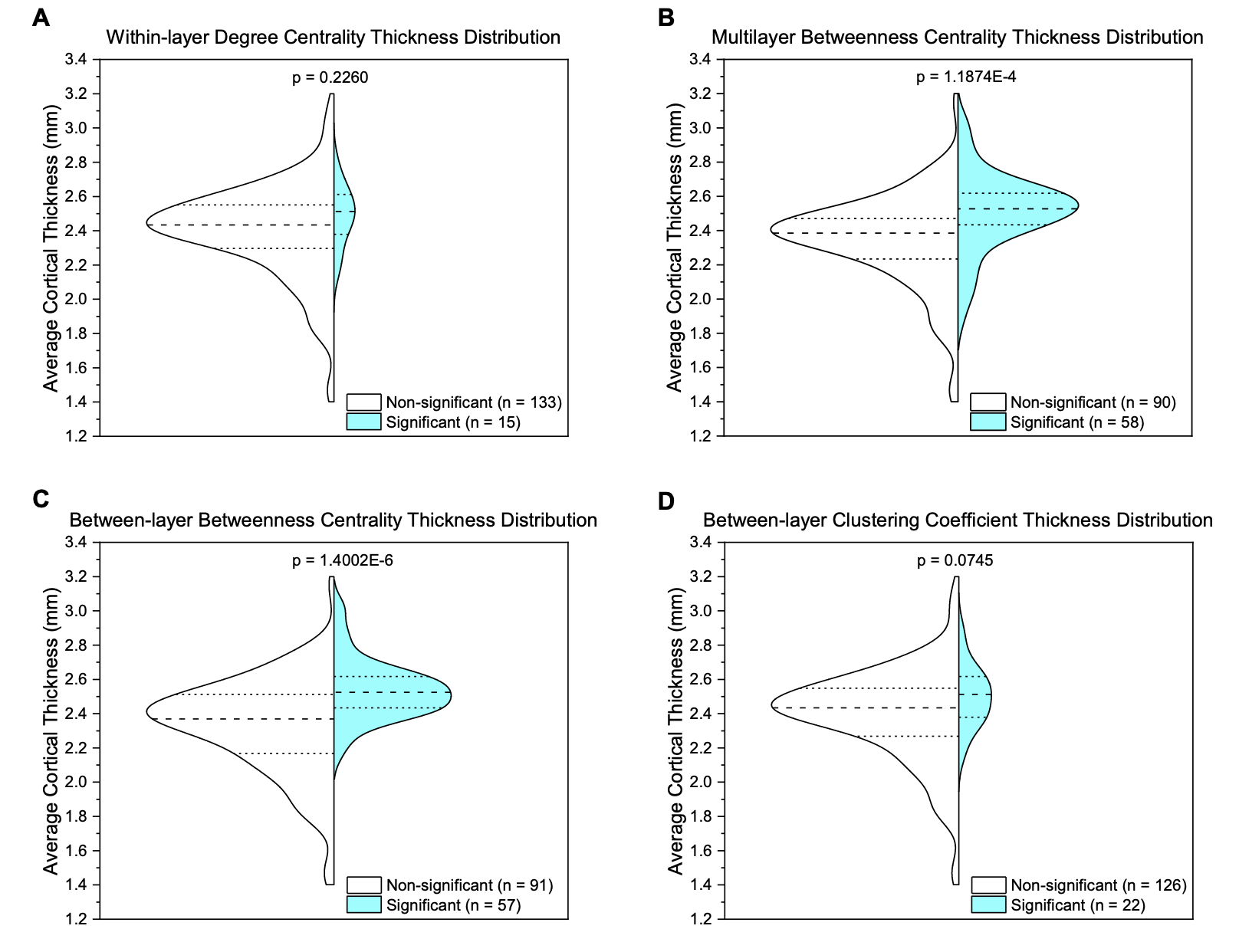

**Figure S30.** Distribution of thickness values for non-significant versus significant nodes for methods with greater than ten significant nodes: **(A)** within-layer degree centrality, **(B)** multilayer betweenness centrality, **(C)** between-layer betweenness centrality, and **(D)** between-layer clustering coefficient. The white distribution plot (left side) represents non-significant nodes while the teal distribution plot (right side) represents significant nodes. Significance was calculated using a t-test [MATLAB-ttest2].
